# Benchmarking SNP-Calling Accuracy Against Known *Citrus* Pedigrees Reveals Pangenome Advantages Over Linear References

**DOI:** 10.64898/2026.04.07.716967

**Authors:** Ryan Kuster, Patrick Sisler, Khushwant Singh, Lu Yin, Shade Niece, Robert Krueger, Chris Dardick, Manjunath Keremane, Chandrika Ramadugu, Margaret Staton

**Author notes:** Corresponding author: Margaret Staton, 2505 EJ Chapman Dr 370 PBB, Knoxville, TN 38996.

## Abstract

**Background:** Pangenomes are a promising new approach to genomics that can reduce reference bias in genotyping, but the reliability of such a data model remains unclear in tracking variation across species. To test the utility of graph-based pangenomes for interspecific breeding, we developed a Minigraph-Cactus super-pangenome representing four *Citrus* species derived from the founder lines of a citrus breeding program. To benchmark SNP calling accuracy using graph and linear-based approaches, we performed whole genome short read sequencing for two sets of pedigreed progeny: 30 F1 hybrids and 244 advanced hybrids from an F1 crossed with a parent not included in the pangenome.

**Results:** The linear approach yielded more SNP calls than the graph-based approach, however, both methods exhibited similar Mendelian Inheritance Error Rates (MIER) in a tool-dependent manner. Reconstruction of parental haplotype blocks in the advanced hybrids revealed a striking improvement in performance in the pangenome graph-based calls, suggesting MIER is vulnerable to error when reference bias influences both parental and progeny genotype calls. Masking of regions diverged from the reference path improved MIER accuracy metrics and haplotype block reconstruction in both the linear and graph-based SNP calls.

**Conclusions:** In non-model systems, inheritance patterns observed from pedigreed hybrids provide a framework for benchmarking variant-calling accuracy using pangenomes. SNP miscalls originating from diverged regions can falsely satisfy MIER filters, thus we recommend haplotype blocks. The inherent structure of the pangenome graph has promising applications for removing regions of unreliable mapping quality, which cannot otherwise be reliably removed using traditional filtering metrics.

## Background

Single linear reference genomes have traditionally been considered sufficient for advancing genomic research for thousands of species, especially given the historic barriers of cost and time. Because a linear reference genome comprises a single haploid sequence, it cannot capture the full genetic diversity present within a population. This creates reference bias, where reads containing variation from the reference genome fail to map properly or remain unmapped. Misaligned reads result in decreased single nucleotide polymorphism (SNP) calling accuracy, especially in regions with increased amounts of structural variation or repeats [1–4].

The reduction of long read sequencing costs and improvement in assembly and scaffolding technology have made it more tractable to generate numerous reference genomes for individuals to better represent diversity within and across species. These linear references can be integrated into graph-based pangenomes that capture both small (SNPs/indels) and large structural variants (SVs) [5,6]. Pangenomes from diverse species have demonstrated the ability to reduce reference bias by increasing the rate and accuracy of read mapping, which in turn, increases SNP calling accuracy [3,4,7–10]. In humans, SNP calling accuracy is benchmarked with gold standard truth sets such as the Genome in a Bottle (GIAB) consortium [3,9,11]. For

other systems lacking ground truth variant sets, such as pig (Miao et al. 2025) and tomato (Zhou et al 2022), simulated reads have demonstrated increased SNP calling accuracy when using a graph-based pangenome compared to a linear reference genome. Though largely absent from non-human pangenome studies, SNP calling accuracy can also be measured by leveraging trios (maternal-paternal-offspring samples) or pedigreed populations, where Mendelian inheritance patterns of individual SNPs or haplotype blocks can reveal incorrectly called SNPs [12–14].

Super-pangenomes, which span more than one species, provide a framework for analyzing genomic evolution across clades, modeling admixture and introgression in wild populations, and tracking variation within interspecific breeding programs [7,15–17]. Because read mapping across species using a single linear reference exhibits higher rates of reference bias than within-species read mapping, a pangenome graph-based approach may offer substantial benefits in reducing reference bias, improving variant calling, and enabling accurate genotyping of SVs. However, high heterozygosity and extensive structural variation are challenging to model as graphs. To make read mapping and genotyping more computationally tractable, these graphs are usually clipped, i.e., sequence segments that do not align well are removed [8,9,11,18]. The reliability to call SNPs from hybrid individuals in super-pangenomes, particularly in regions of graph clipping or structural complexity, has been insufficiently characterized thus far. While larger variants such as structural variants and indels become tractable for genotyping with pangenomes, SNPs remain the primary variant type that can be used in existing software packages for QTL (quantitative trait locus) mapping, genome wide association studies (GWAS), population genetics, and other genetic evaluations.

By incorporating all founder lines into a pangenome graph, breeding programs can capture the full spectrum of genomic variation, creating a comprehensive catalog of complex, diverged loci and an anchor for non-reference sequences. This framework enables the use of cost-effective, low coverage short read sequencing to accurately genotype progeny and track introgression [19–21]. Recent applications of pangenomes to breeding programs have already facilitated new trait locus discovery in wild and domesticated species, demonstrating increased power in detecting genotype-to-phenotype associations [7,10,21,22].

*Citrus* is a timely use case for investigating the utility of graph-based pangenomes in genotyping for interspecific breeding programs. Worldwide citrus production is currently being devastated by Huanglongbing (HLB), also known as citrus greening disease, which is caused by the bacterium *Candidatus* Liberibacter asiaticus (CLas) [23]. To combat this, wild Australian lime species (*C. australasica*, *C. australis*, and *C. inodora*) that exhibit heritable HLB tolerance are being rapidly incorporated into domesticated citrus breeding programs [24–27]. With an estimated evolutionary divergence time of three to five million years [28] and high structural variation within wild Australian lime species [25,29], a pangenome may offer a more accurate platform than a linear reference for identifying and tracking genetic variation across interspecific *Citrus* breeding generations. Here, we construct a *Citrus* super-pangenome using haplotype resolved reference genomes from six founder lines representing four species. We use short read sequencing of interspecific F1 and advanced hybrid samples to address three main objectives. First, we benchmark SNP calling accuracy by comparing a linear reference against a graph-based pangenome across six combinations of read mapping and SNP calling approaches. We then assess the underlying structure of the pangenome graph for correlation of clipping and SNP calling accuracy. We also evaluate the impact of representation bias by determining if SNP calling accuracy is negatively impacted when a parental genome is excluded from the pangenome graph or when founder paths consistently diverge from the reference. Building on these results, we design a masking strategy to filter unreliable SNPs originating from complex graph regions.

## Results

### *Citrus* Super-Pangenome

A Minigraph-Cactus (MC) super-pangenome was constructed from six haplotype-resolved, chromosome scale genome assemblies, totaling 12 haplotypes. This included one individual from each of three wild Australian *Citrus* species (*C. australis*, *C. australasica*, and *C. inodora*) and one individual from each of the three mandarin cultivars of *C. reticulata* (‘Fortune’, ‘Wilking’, and ‘Fallglo’), ranging in total scaffolded chromosome length from 272 Mb to 303.8 Mb ([24–27]**; Supplemental Table 1**). A phylogenetic analysis of single-copy genes from the six genomes plus two more diverged *Citrus* species confirmed the expected phylogeny, with the Australian limes clustering as an outgroup to the domesticated mandarins (**Figure 1**). The *C. reticulata* ‘Fortune’ primary haplotype was used as the reference backbone of the pangenome graph. Our total super-pangenome length was 667.6 Mb and contained 51.6 million nodes with 70.6 million edges and 2,859 paths. To assess how the super-pangenome expanded with each additional haplotype, we summarized node growth and node sharing using Panacus (**Figure 2**) [30]. Core nodes (present in all 12 haplotypes) accounted for 12% (6.2 million), while private nodes (unique to a single haplotype) accounted for 17% (9 million). The largest fraction, 70.7% (36.5 million), was comprised of dispensable nodes (shared by two or more haplotypes, but not all 12).

**Figure 1.**
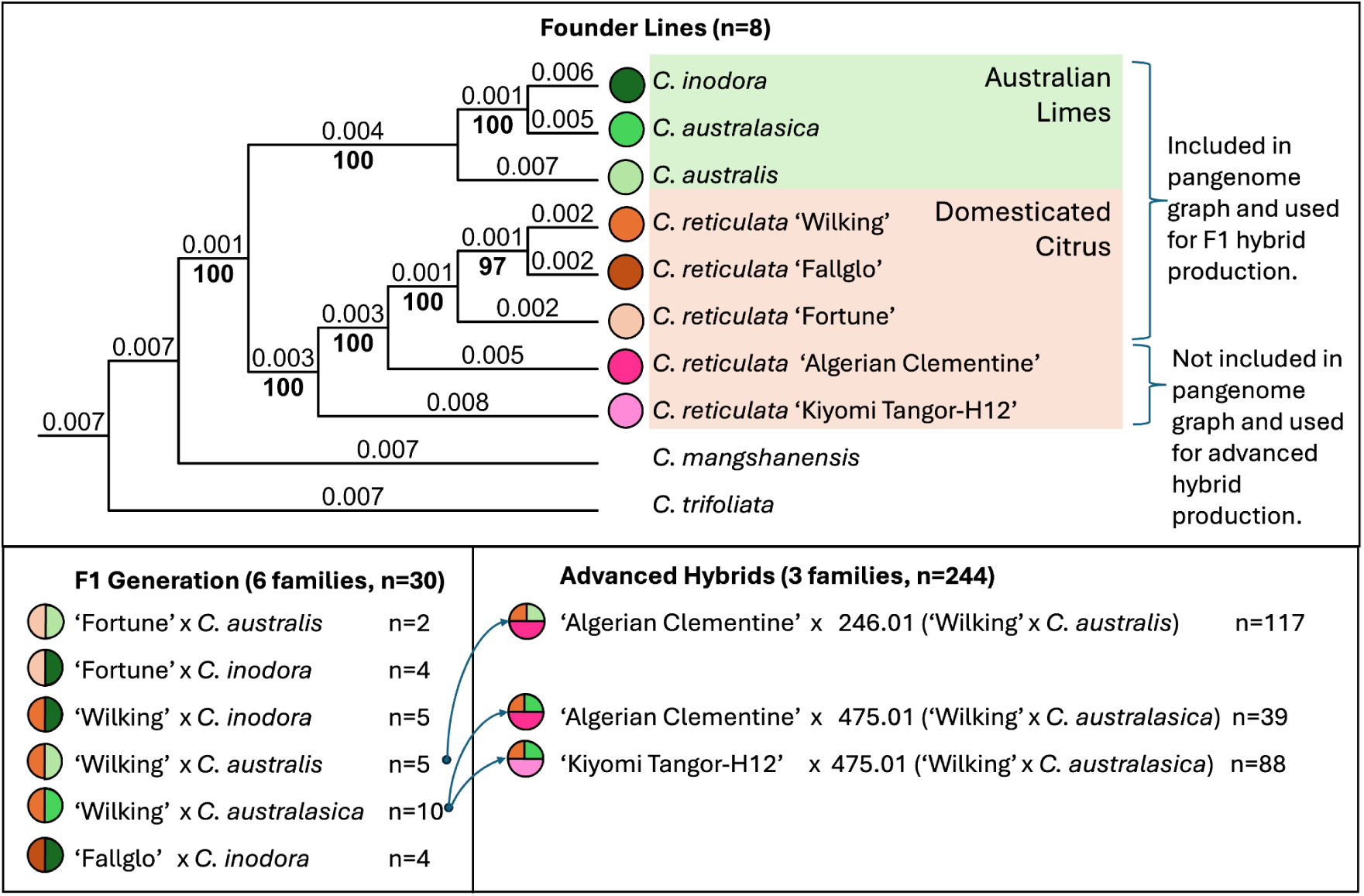
(Top) *Citrus* phylogeny. Bold values at cladogram nodes represent bootstrap values and branch labels represent phylogenetic distance. **(Bottom)** Visual summary of breeding program with F1s on the left and advanced hybrids on the right.

**Figure 2:**
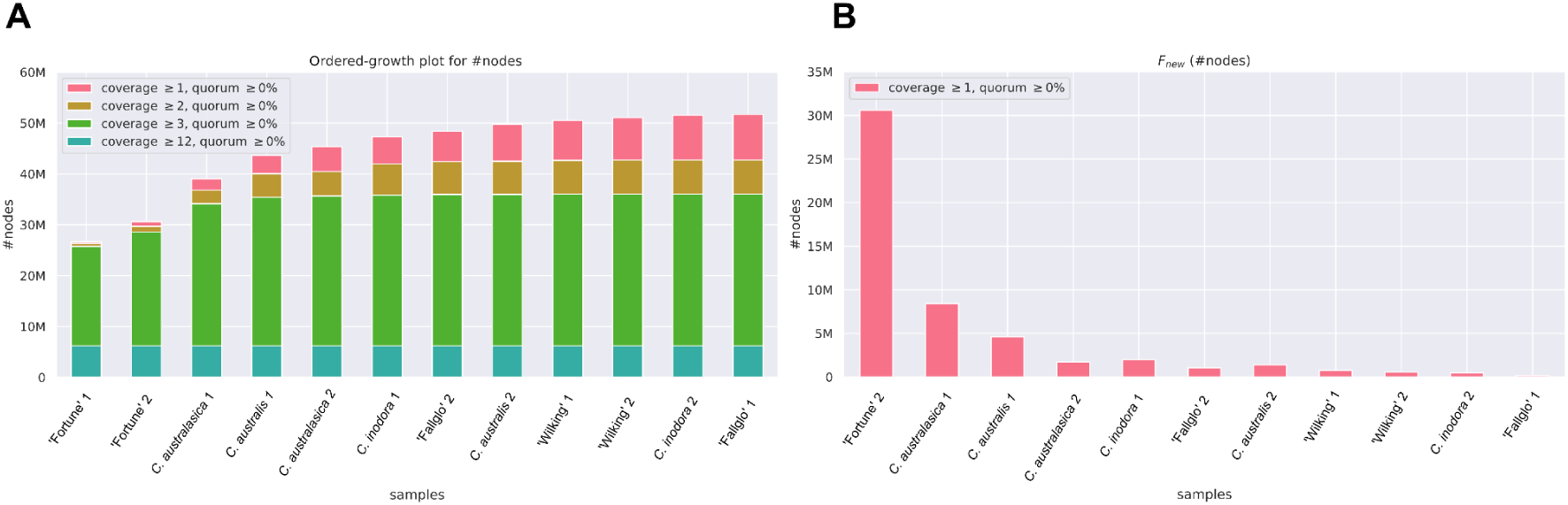
**(A)** Growth curve of nodes in the *Citrus* super-pangenome found in one or more, two or more, three or more, or all twelve haplotypes (core). **(B)** New nodes contributed by the addition of each sample with sample labels of “1” referencing the primary haplotype, and “2” referencing the alternate haplotype.

To limit graph complexity, MC clips individual paths from regions that are highly repetitive or are difficult to align. To quantify the amount of clipped sequences during pangenome construction, we examined each haplotype’s path. While clipping occurred across all haplotypes and scaffolds (**Figure 3A and 3B**, **Table 1**), the amount of clipping varied substantially among haplotypes.

**Figure 3:**
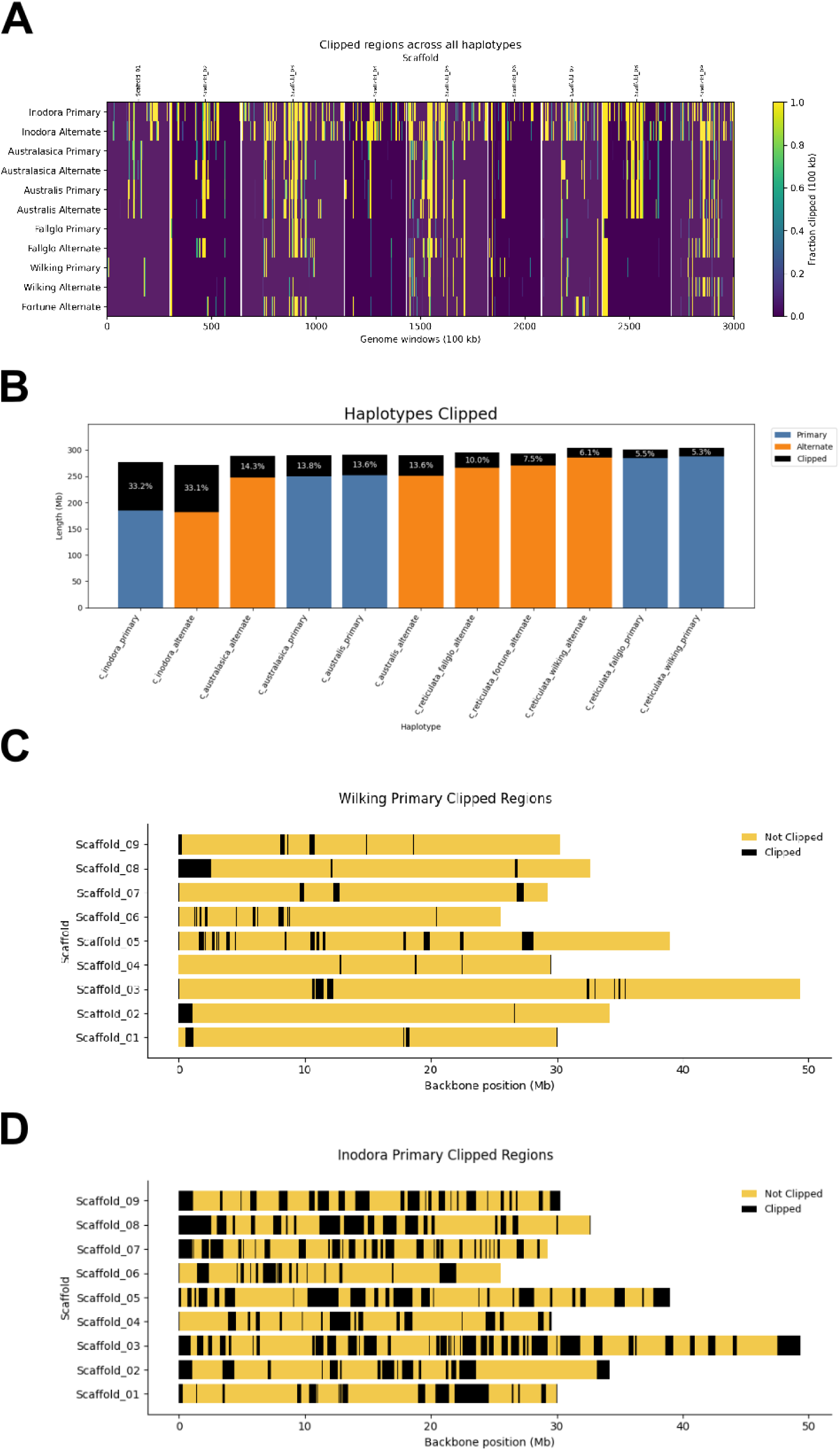
**(A)** Heatmap showing clipped regions from 11 pangenome haplotypes, plotted in the genome coordinates of ‘Fortune’ in 100 kb windows. ‘Fortune’ primary is not included, as the backbone is not clipped during pangenome construction. **(B)** Percent of bases clipped from the pangenome graph by haplotype. **(C)** Clipping by chromosome for the least clipped haplotype, ‘Wilking’ primary. **(D)** Clipping by chromosome for the most clipped haplotype, *C. inodora* primary.

**Table 1.** Number and percent of clipped bases in each haplotype. ‘Fortune’ primary is never clipped as the reference path for the graph.

| Haplotype | Clipped (Mb) | Haplotype Size (Mb) | Percent Clipped |
| --- | --- | --- | --- |
| *C. inodora* Primary | 92.03 | 276.93 | 33.23 |
| *C. inodora* Alternate | 89.94 | 272.04 | 33.06 |
| *C. australasica* Primary | 40.07 | 290.01 | 13.81 |
| *C. australasica* Alternate | 41.31 | 289.02 | 14.29 |
| *C. australis* Primary | 39.68 | 291.56 | 13.61 |
| *C. australis* Alternate | 39.36 | 289.99 | 13.57 |
| 'Fallglo' Primary | 16.48 | 300.99 | 5.47 |
| 'Fallglo' Alternate | 29.49 | 295.33 | 9.98 |
| 'Fortune' Alternate | 21.99 | 293.01 | 7.51 |
| 'Wilking' Primary | 15.98 | 303.77 | 5.26 |
| 'Wilking' Alternate | 18.41 | 303.76 | 6.06 |

Australian founder lines were clipped more frequently than mandarin lines, with total clipped bases ranging from ∼5% for the ‘Wilking’ primary haplotype to over 33% for the *C. inodora* primary haplotype (**Figure 3C and 3D, Supplemental Figures 1-11**). Within assemblies of one individual, the amount of sequence clipping varied by <1% between haplotypes for all except the ‘Fallglo’ assembly, where the primary haplotype was clipped at 5.5% while the alternate haplotype was clipped at 10.0%.

### SNP Benchmarking using F1s

To better understand the accuracy and performance of SNP calling in pangenome versus linear reference approaches, we analyzed short-read whole genome sequencing data from 30 interspecific F1 hybrids and their corresponding parental lines (**Figure 1**). Across these samples, the mean depth was 39x (ranging from 23.5x to 57.5x; **Supplemental Table 2**). These F1 individuals originated from crosses between the mandarin and Australian lime founders in the MC super-pangenome. To benchmark variant calling quality, we evaluated six variant calling approaches in total. Three linear reference-based approaches used BWA-MEM for read alignment, followed by variant calling by BCFtools, GATK, or DeepVariant (hereafter identified with the prefix “linear-”). Three graph-based pangenome approaches used vg giraffe alignment, followed by either vg pack + vg call (referred to as “vg-pack-and-call”) or vg surjection to the linear reference for processing with BCFtools or GATK (hereafter identified with the prefix “vg-surject”).

For the linear-based alignment, reads were mapped to the ‘Fortune’ primary genome using BWA-MEM with an average mapping rate of 95.3%. For the graph-based alignment, mapping with vg giraffe achieved a similar average mapping rate of 95.2%. Surjected read mapping yielded 57.07% average mapping rate to the reference path with more variation per sample than the other two approaches (**Figure 4A**). Lower mapping rates after surjection are expected, as this step discards any reads where graph alignments do not have a valid linear coordinate in the reference path.

**Figure 4.**
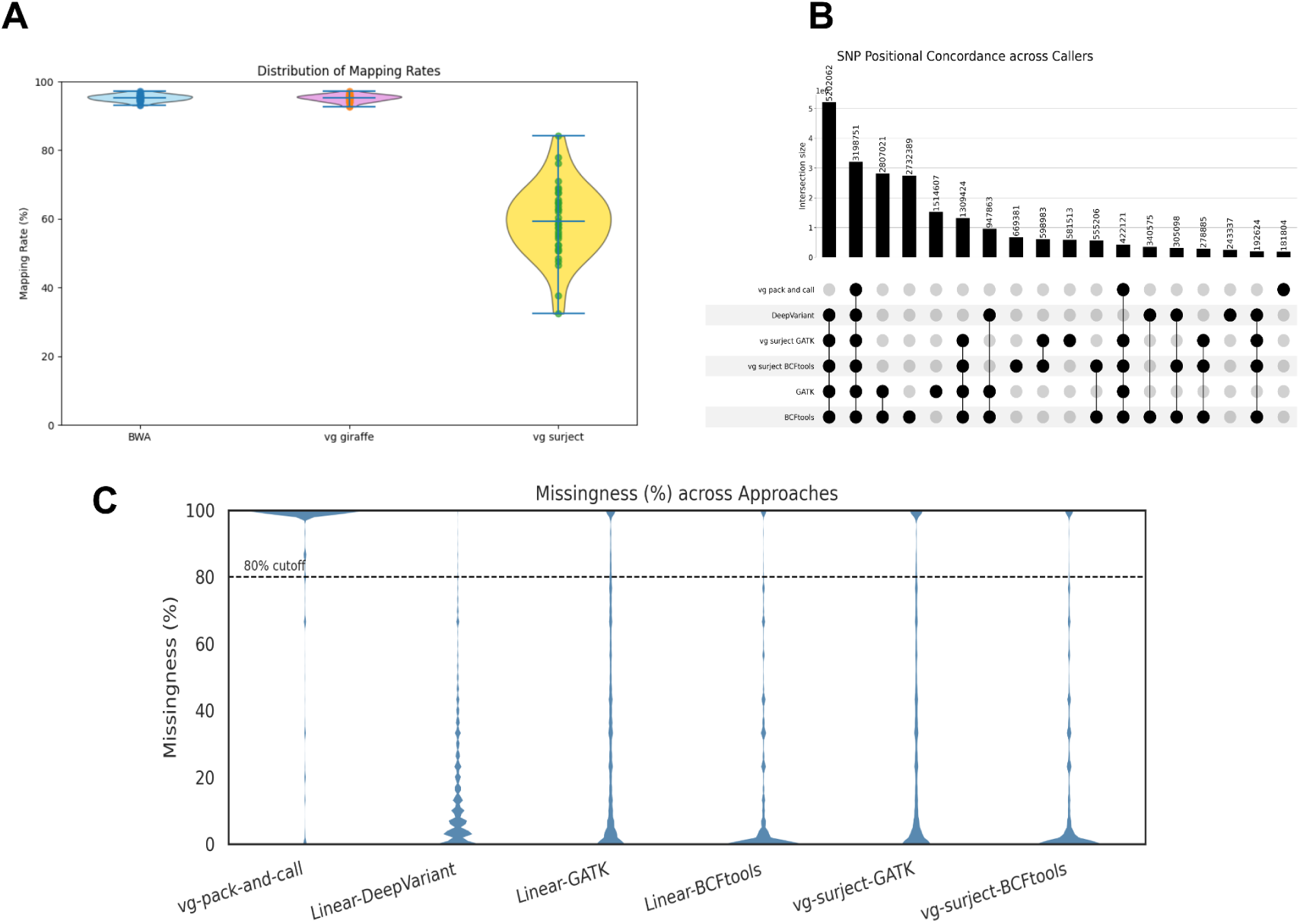
Results from six approaches for SNP calling comparing a linear reference and a pangenome. **(A)** Mapping rates of the whole genome sequencing data from 30 F1’s and six parental lines when aligning with BWA-MEM to the linear reference of ‘Fortune’ primary haplotype assembly (“BWA”), vg giraffe to super-pangenome graph (“vg giraffe”), and vg surject, which uses the vg giraffe mappings and projects mappings to the linear reference path of ‘Fortune’ primary (“vg surject”). **(B)** Upset plot of shared SNP positions across the six SNP calling approaches. **(C)** Missingness of SNP calls in each approach. Dotted line marks an 80% cutoff.

In the F1s, after filtering, the resulting number of SNPs across callers varied substantially, ranging from ∼4.9 million (vg-pack-and-call) to ∼19.1 million (linear-BCFtools) (**Table 2**).

**Table 2.** Performance summary table of SNP calling approaches.

| Caller | SNPs | Missing<br>Genotype Calls<br>Percentage | MIER<br>(exclude<br>missing) | MIER<br>(include<br>missing) |
| --- | --- | --- | --- | --- |
| **F1 Generation (n = 30)** |  |  |  |  |
| **vg-pack-and-call** | 4,914,458 | 59.50% | 1.24% | 92.66% |
| **linear-DeepVariant** | 10,958,948 | 8.60% | 22.01% | 34.79% |
| **linear-GATK** | 16,274,623 | 12.60% | 9.54% | 38.77% |
| **vg-surject-GATK** | 12,686,117 | 12.00% | 9.25% | 39.40% |
| **linear-BCFtools** | 19,127,522 | 4.70% | 7.81% | 23.81% |
| **vg-surject-BCFtools** | 13,921,411 | 5.10% | 8.35% | 26.39% |

| **Advanced Hybrid Generation (n = 244)** |  |  |  |  |
| --- | --- | --- | --- | --- |
| **linear-BCFtools** | 20,019,741 | 10.31% | 7.13% | 16.71% |
| **vg-surject-BCFtools** | 14,886,746 | 11.38% | 7.10% | 17.67% |
| **Advanced Hybrid Generation After Masking (n = 244)** |  |  |  |  |
| **linear-BCFtools** | 11,386,766 | 7.56% | 5.75% | 12.92% |
| **vg-surject-BCFtools** | 11,330,756 | 7.27% | 5.80% | 12.60% |

Linear-BCFtools had the highest number of unique SNPs (2,732,389), and vg-pack-and-call had the fewest unique SNPs (181,804). For both vg-surject-BCFtools and vg-surject-GATK, using surjected reads yielded fewer SNPs than the linear approach. Vg-pack-and-call yielded the fewest SNPs overall. A total of 3,198,751 shared SNPs were identified based on genomic position among all six approaches (**Figure 4B**). Notably, by excluding vg-pack-and-call from the comparison, the number of shared SNPs increased 62.61% to 5,202,062.

For each pair of callers, we assessed genotype concordance call per sample at shared SNP positions (i.e., both callers identified the sample as 0/0, 0/1 or 1/1) (**Figure 5**). Among comparisons excluding vg-pack-and-call, genotype concordance was consistently high (>93%), with particularly strong concordance between vg-surject approaches and their linear counterparts using the same SNP caller. In contrast, comparisons involving vg-pack-and-call showed lower concordance rates with all other SNP callers, ranging from 57.67% to 68%.

**Figure 5.**
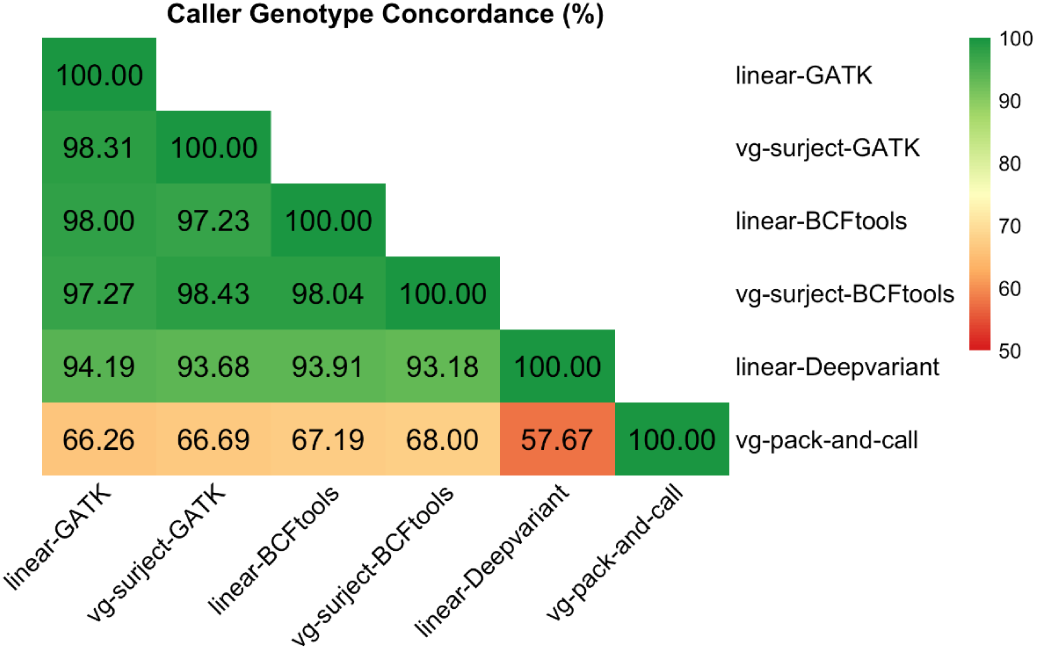
Agreement of genotype calls of F1s at shared SNP positions across the six SNP calling approaches.

Additionally, we quantified the proportion of missing genotypes produced by each caller’s entire SNP callset (**Figure 4C**, **Table 2**). Missing genotype rates were the lowest for BCFtools followed by DeepVariant, then GATK, while vg-pack-and-call produced substantially higher levels of missing genotypes. The hybrid surjection based call approaches performed similarly to their linear counterparts. For example, linear-GATK showed 12.56% missing genotypes compared to 11.98% for vg-surject GATK. Likewise, linear-BCFtools showed 4.71% missing genotypes compared to 5.14% using surjection.

To further assess our SNP-calling approaches, we assessed the expected Mendelian inheritance error rate (MIER) for each caller aggregated across the 30 parent-F1 trios (**Table 2**). After excluding missing calls, callers exhibited error rates ranging from 1% to 22%, with vg-pack-and-call performing the best and linear-DeepVariant performing the worst. Linear-BCFtools, linear-GATK, and their vg-surject counterparts had a moderate error rate ranging from 8% to 10%.

To evaluate reference bias in read mapping across our approaches, we analyzed allelic ratios at putative heterozygous sites identified directly from read pileups where an unbiased expectation is 0.5. Putative heterozygous sites were defined as positions with a minimum coverage of 10x and a minimum minor allele frequency of 25%, following the removal of low quality and secondary alignments. Across the 30 F1 hybrids, reads aligned to the linear reference produced a mean alternate allele fraction of 0.4569, ranging from 0.413 to 0.470. In contrast, reads aligned to the pangenome reference using vg giraffe were notably closer to the unbiased expectation, yielding a mean alternate allele fraction of 0.489, ranging from 0.427 to 0.495 (**Figure 6**).

**Figure 6.**
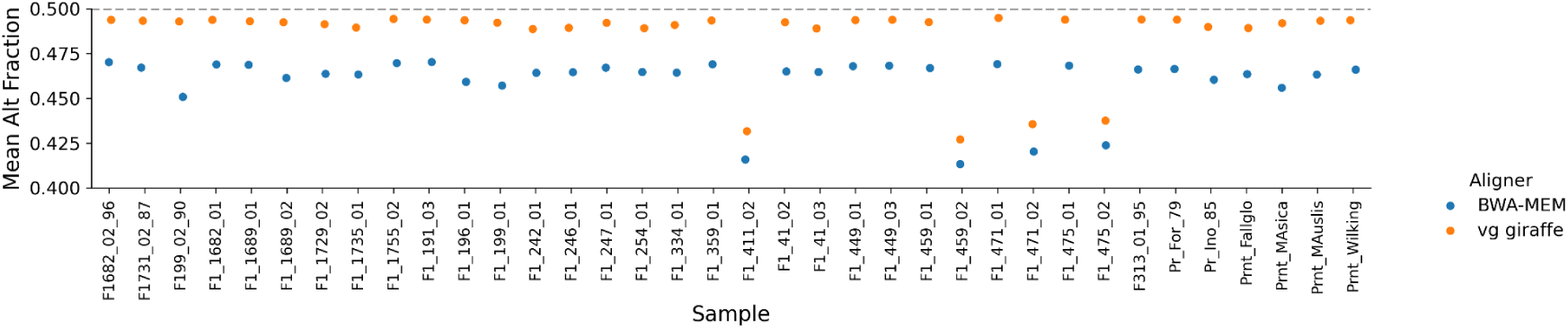
Plot comparing the mean alternate allele frequency across 30 F1 and six parental lines present on the pangenome when aligned with BWA-MEM versus vg giraffe. In an unbiased scenario, allelic ratios at putative heterozygous sites should center at 0.5.

### Graph Clipping Correlation with MIER in the F1s

We hypothesized that sequence clipping, particularly within the highly clipped Australian lime lineages, may introduce errors in SNP calling for pangenome-based approaches. To investigate this, we correlated MIER from the vg-surject-BCFtools approach with the number of clipped paths in the pangenome. F1s were grouped by their Australian lime parent (*C. inodora*, *C. australis*, or *C. australasica)*, then SNPs were categorized by whether they occurred in regions with zero, one, or two clipped Australian lime haplotypes (**Figure 7A**). Kruskal-Wallis tests revealed a significant difference in MIER across clipping categories for all three species (H*_inodora_* = 15,339; H*_australasica_ =* 77,292; and H*_australis_ =* 65,332; *p* < 0.001). Post-hoc Dunn’s tests (p < 0.05) indicated that, for all species, sites with zero versus two clipped haplotypes differed significantly. For *C. australisica* and *C. indora,* we observed a significant difference between sites with one clipped haplotype versus sites with two clipped haplotypes. For *C. australasica* and *C. australis*, sites with no clipped haplotypes vs one clipped haplotype were also significant. Median MIER revealed differences between *C. inodora*, the most highly clipped haplotypes in the pangenome graph, in comparison to *C. australasica* and *C. australis*. The latter two have median MIER values of 0 in regions with no clipping or one haplotype clipped. In contrast, *C.inodora* exhibited consistently higher medians across all categories (0.0769).

**Figure 7.**
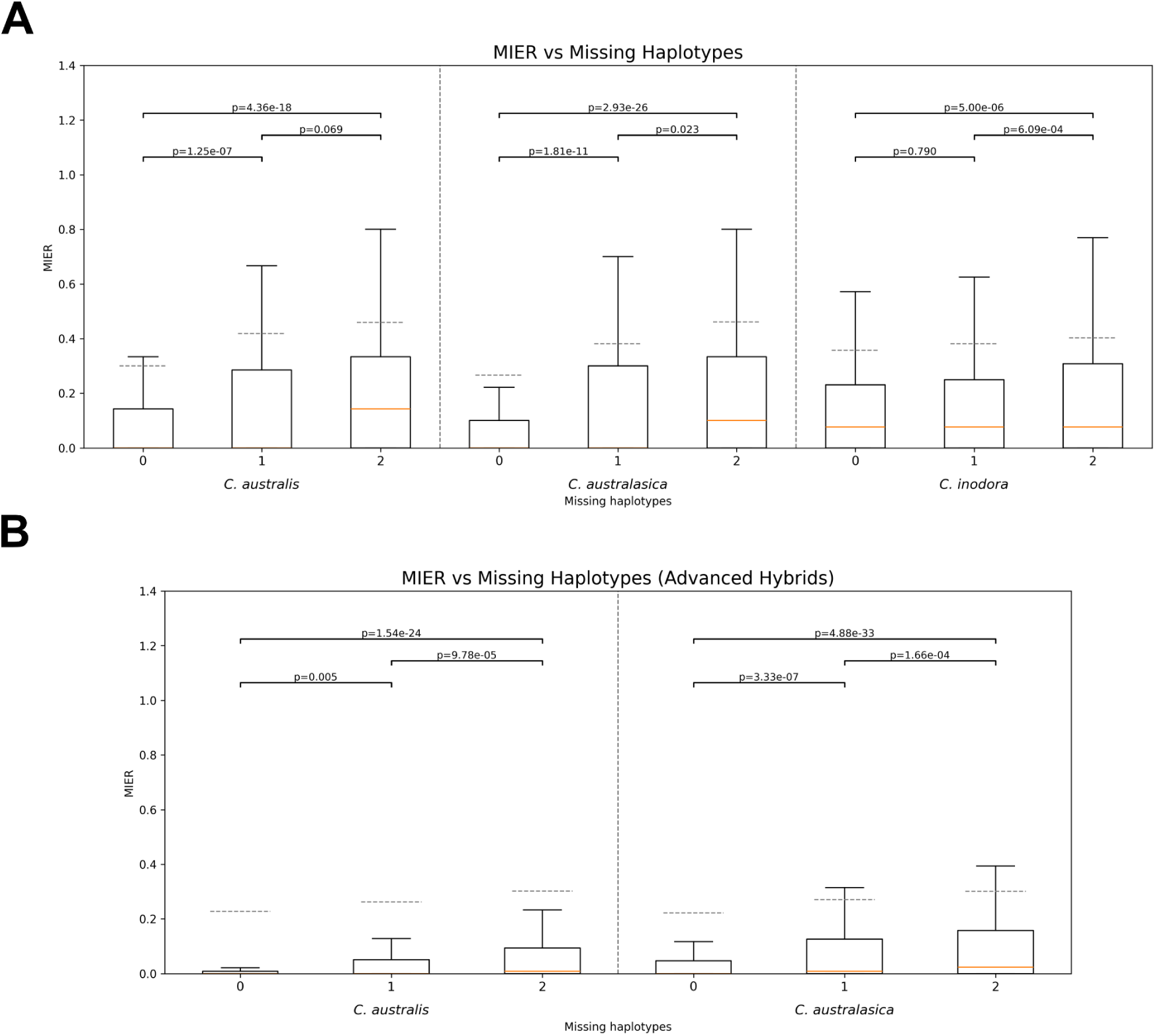
MIER rates increase in regions where parental haplotypes are clipped from the pangenome graph. On the x-axis, *Citrus* trios are grouped by the Australian lime founder line. MIER was calculated for SNPs originating from parts of the pangenome graph where zero, one, or two (both) Australian lime haplotypes were clipped in the super pangenome graph for **(A)** F1 hybrids and **(B)** advanced hybrids. Significant pairwise group differences (*p* < 0.05) were calculated post-hoc using Dunn’s test.

### MIER Benchmarking in Advanced Hybrids (AD)

To assess the performance of SNP calling in individuals with parents absent from the pangenome founder lines, we evaluated a panel of 244 advanced hybrids consisting of three families (**Supplemental Table 2**). These hybrid individuals were derived from an F1 pollen parent and a domesticated *Citrus* seed parent not included in the pangenome graph, ‘Algerian Clementine’ or ‘Kiyomi Tangor-H12’ (**Figure 1**).

Based on the F1 results, we only compared two SNP calling approaches, linear-BCFtools and vg-surject-BCFtools in the advanced hybrids (AD). The vg-surject-BCFtools calls produced fewer SNPs (14,886,746) than linear-aligned calling (20,019,741), with 11,363,316 SNPs sharing the same genomic position between the two sets (**Table 2**). The average missing call rate for the advanced hybrid trios was 10.31% and 11.38% for the linear and surjected calls, respectively. This was over twice the ∼5% missingness rate observed from the same SNP calling approaches in the F1s, likely because of the missing parents in the pangenome graph. Despite this, MIER excluding missing data for the 244 advanced hybrids was comparable to the 8% MIER observed in the F1s, with a marginal improvement in accuracy for vg-surject-BCFtools calls (MIER = 7.10%) over linear-BCFtool calls (MIER = 7.13%).

To determine whether the significant correlation between graph clipping and MIER extends beyond F1 crosses in pangenome-based SNP calling, we performed the same analysis with the advanced hybrids (AD). Despite the reduced contribution of Australian lime in these AD crosses (approximately 25% vs 50% in F1s), we observed similar patterns: MIER increased with more clipped haplotypes (**Figure 7B**). For both *C. australasica* and *C. australis* derived AD, Kruskal-Wallis tests indicated significant differences in MIER across sites with zero, one, and two clipped haplotypes (*p* ≈ 0). Post-hoc Dunn’s tests revealed that all pairwise comparisons among haplotype clipping classes were significant (p < 0.01).

We further examined the genotypes violating Mendelian inheritance for region-specific and genotype-specific patterns of erroneous alignment. Assessment of 100 kb windows indicated a genome-wide tendency for linear-BCFtools calls in trio progeny to yield higher rates of false heterozygous calls (**Figure 8, Supplemental Figures 12-20**). In contrast, vg-surject-BCFtools SNPs exhibited localized spikes in false homozygous calls relative to linear-BCFtools. In addition to the strong correlation of clipping to MIER, we hypothesized that other features of the pangenome graph, specifically unclipped founder paths diverging from the reference backbone, could also explain error patterns. Local graph composition was investigated by examining nodes within 150 bp upstream and downstream of each miscalled SNP along the ‘Fortune’ primary haplotype reference path. For this graph subpath, we then determined the average number of other founder paths (from 0 to 11) that share these nodes, which we defined as “founder presence”. We found moderate positive correlations between correct calls and the founder presence along these 150 bp local subpaths (Spearman’s *r* = 0.45). Analysis of per-node founder presence-absence within the pangenome structure revealed that graph-derived false-homozygous SNPs consistently occur in localized regions where both haplotypes of founder-line parents are absent from the surjected reference path. This spatial correlation between missing graph paths and genotyping error is further evidenced when visualizing these patterns across 100 kb windows (**Figure 9, Supplemental Figures 21-29**).

**Figure 8.**
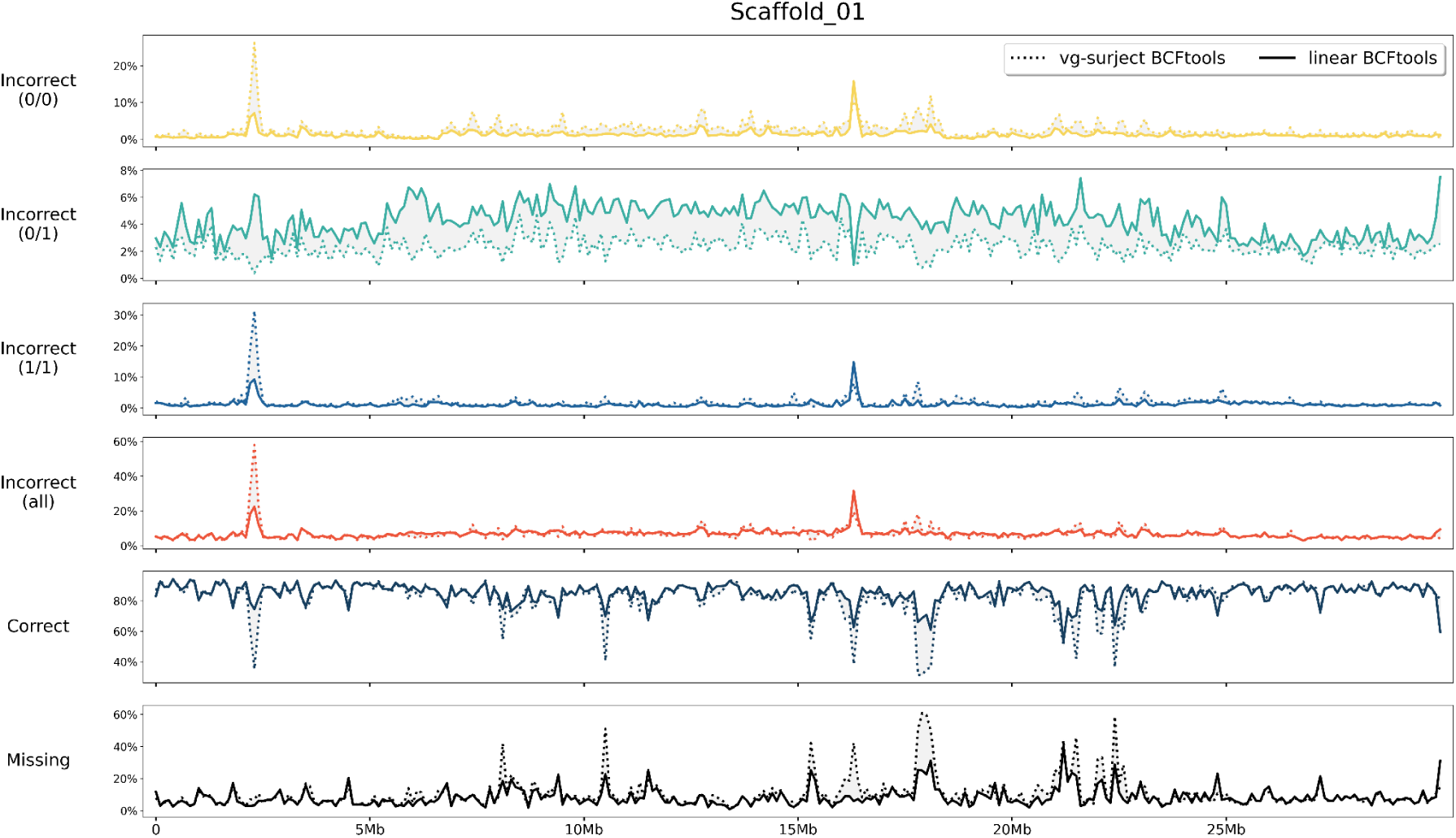
Summary comparison of genotype call types from linear-BCFtools (solid line) and vg-surject-BCFtools (dotted line) across 100 kb windows on chromosome 1 (additional chromosomes in **Supplemental Figures 12-20**). Panels from top to bottom: incorrect homozygous reference (yellow), incorrect heterozygous (green), incorrect homozygous alternate (blue), incorrect total (red), correct (dark blue), missing (black).

**Figure 9.**
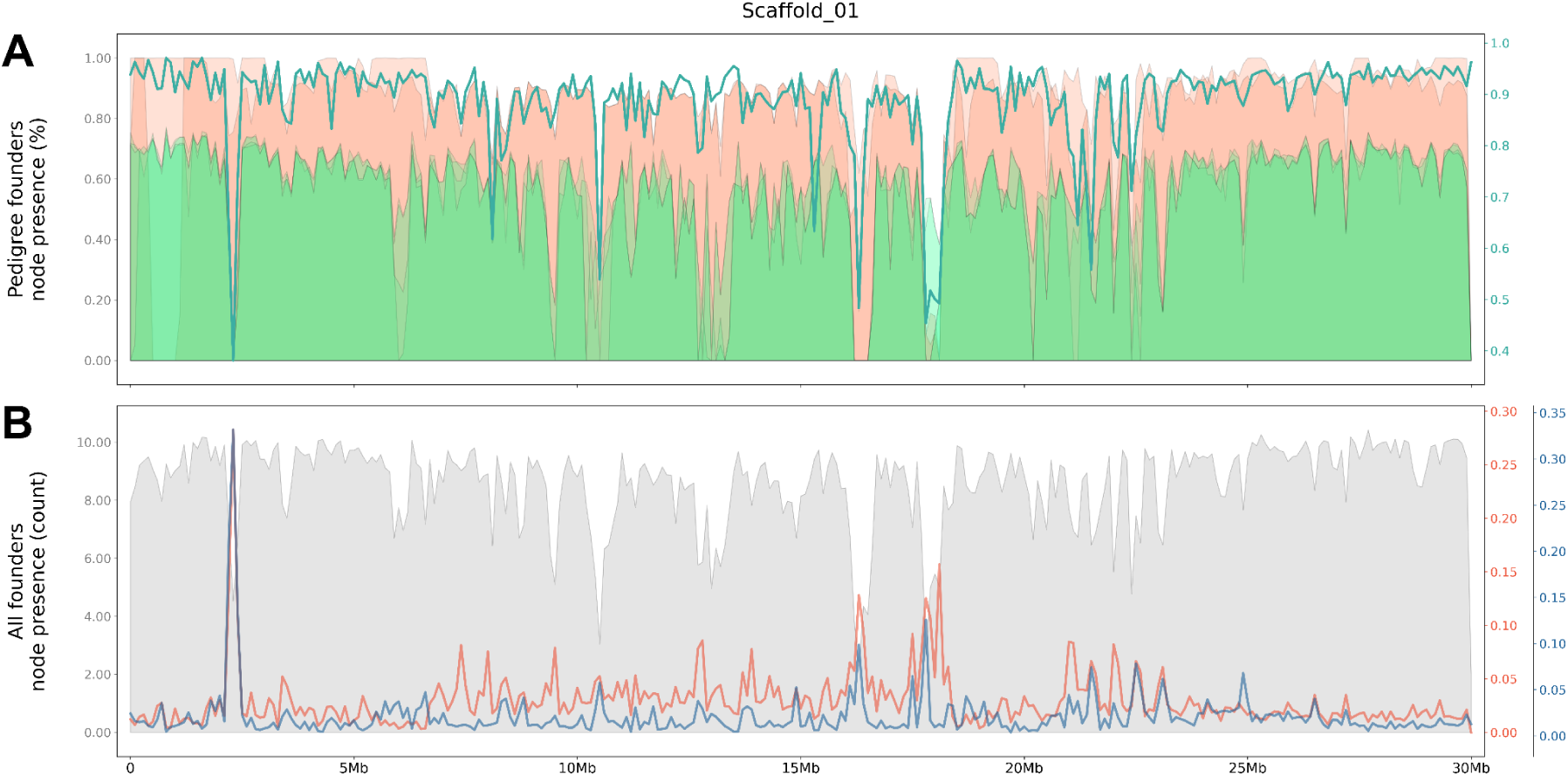
Relationship of founder presence in the graph nodes per 100 kb windows along chromosome 1. **(A)** The node presence of pedigree founders (orange = mandarin; green = Australian lime) vs. the frequency of genotypes per window that are correct (green line). **(B)** The number of founders present per-node (grey) vs. the rates of incorrect homozygous reference calls (red), and incorrect homozygous alternate calls (blue). (Other chromosomes available in **Supplemental Figures 21-29**).

### Advanced Hybrids (AD): Haplotype Blocks

While MIER is a standard benchmark, it is an imperfect metric for SNP calling accuracy. Specifically, concurrent calling failures in both parent and progeny, often stemming from regional mapping artifacts described above, can remain undetected by MIER. To address this, we utilized advanced hybrids to provide an independent metric for SNP accuracy based on heritable haplotype blocks. We defined the haplotype blocks inherited from the Australian lime founders resulting from recombination in the F1s (**Supplemental Figure 30**). Haplotype block detection of the founders was not meaningful in the F1 generation due to each individual simply inheriting one allele from the mother and the other allele from the father and the lack of reliable variant phasing. However, this approach proved highly effective in the advanced hybrid generation, utilizing the two-generation pedigree, high heterozygosity between Australian limes and domesticated *Citrus*, and an expected rate of zero to three recombination events per chromosome per generation.

For each advanced hybrid, SNP calls adhering to Mendelian inheritance were used to predict the parental origin (mandarin or Australian lime) of the F1 contributing allele, thereby visually delineating haplotype blocks (**Figure 10, Supplemental Figures 31-39**). Inconsistencies in parental types within otherwise congruent haplotype blocks revealed loci that were falsely called despite meeting Mendelian expectations.

**Figure 10.**
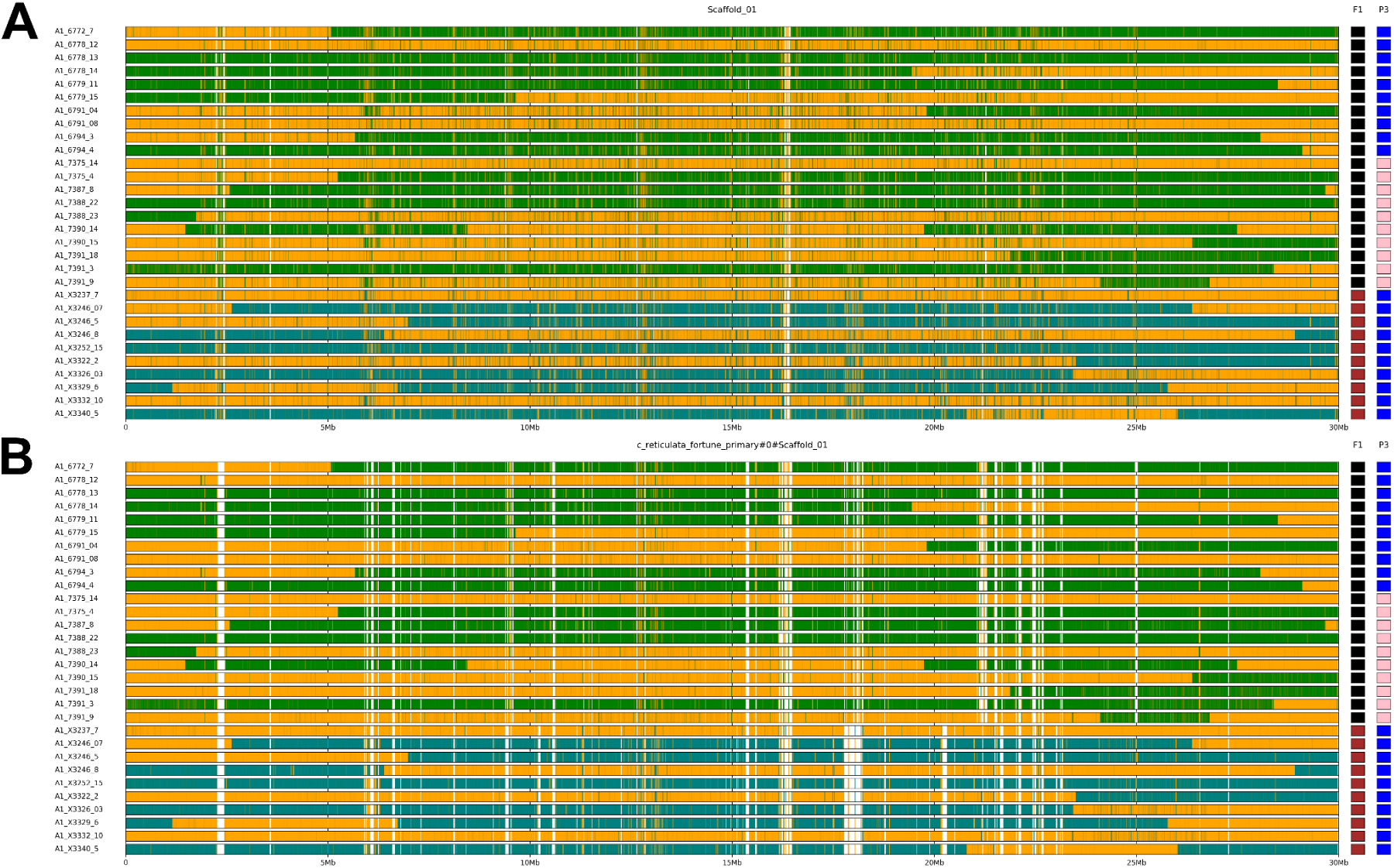
Parental haplotypes reconstructed from **(A)** linear-BCFtools-based variants and **(B)** vg-surject-BCFtools variants. Ten subsets of each of the three hybrid families studied are denoted with the F1 parent (right boxes, black = F1 hybrid 475_01 [‘Wilking’ x *C. australasica*]; brown = F1 hybrid 246_01 [‘Wilking’ x *C. australis*]) and outcrossed parent (right boxes, blue = ‘Algerian Clementine’; pink = ‘Kiyomi Tangor-H12’). Orange blocks indicate the presence of a mandarin grandparent source and the Australian lime contribution is indicated by either green (*C. australasica*) or dark teal (*C. australis*). Miscalled variants within blocks (small regions of opposite color within otherwise congruent color blocks) met Mendelian inheritance expectations. Visualizations of all chromosomes available in **Supplemental Figures 30-38**.

A smoothing algorithm that recodes the SNPs that mismatch the local haplotype context was developed and used to delineate the ground truth haplotype blocks from mandarin and Australian lime derived haplotypes originating in the F1. Using recombination breakpoints inferred from the smoothed blocks, we estimated a recombination rate of 3.02 cM/Mb, which is consistent with previously reported rates for *Citrus* of approximately 3 cM/Mb [31–33]. We first identified haplotype block-discerning SNPs, defined as heterozygous F1 SNPs with homozygous opposite grandparental calls and homozygous outcrossing parents. We examined genotypes from 244 advanced hybrids at the haplotype block-discerning SNPs to compare predicted per-locus parental origins against the parental types in the ground truth, smoothed blocks. Based on this, we found an increase in false calls using the linear-BCFtools approach compared with the vg-surject-BCFtools approach (**Figure 10**). Genome-wide, the vg-surject-BCFtools-based calls produced 454,045,632 SNP genotype calls with a discordance of 2.11% relative to their haplotype block context. In contrast, the linear-BCFtools-based calls produced 393,198,130 SNP genotype calls with a discordance of 5.14% relative to their haplotype block context.

### Advanced Hybrids: Masking

Given our results that clipping and founder path presence both correlate to poor SNP calling accuracy, we hypothesized that the underlying structure of the pangenome graph can be used to filter low quality SNP calls. We performed targeted masking of the reference path where both haplotypes of any parental line of our advanced hybrid population (i.e., *C. australis*, *C. australasica*, or ‘Wilking’) were absent from the reference path for a minimum of six consecutive base pairs. This masking approach removed 3,555,990 SNPs (23.88%) from the vg-surject-BCFtools VCF and reduced the MIER by 1.35 points to 5.8% (**Table 2, Supplemental Table 3**). Applying the same mask to the linear-BCFtools VCF removed 8,632,975 SNPs (43.12%) with a MIER reduction of 1.38 points to 5.75%. Inversely, calls taking place within masked coordinates exhibited higher error rates, with a MIER of 12.4% for the graph-based calls and 9.03% for the linear call sets. The frequency of missing genotypes in F1 trios within these masked regions was more than twice as high using vg-surject-BCFtools, suggesting that F1 parental calls were less frequent in these poor-performing graph regions.

Notably, while the initial call sets (linear-BCFtools and vg-surject-BCFtools) differed by 5.2 million SNPs, the number of SNPs in non-masked regions was nearly identical between the these approaches, with only 56,010 fewer SNPs in the graph VCF and 9,165,272 calls (∼80%) sharing the same genomic position.

Despite only marginal improvements in MIER, the application of masking provided notable visual improvements in the haplotype block for the graph approach relative to the linear approach (**Figure 11**). Strikingly, when we narrowed our haplotype block analysis to those calls within the proposed masking regions, we found 8.18% discordant calls in the vg-surject-BCFtools-based calls (13,023,724 genotypes) juxtaposed with 23.31% discordance in the linear-BCFtools-based calls (53,933,159 genotypes). These results indicate that while linear-based alignments produce a larger proportion of calls within these divergent, masked regions of the reference genome, the calls in these regions are much more frequently incorrect.

**Figure 11.**
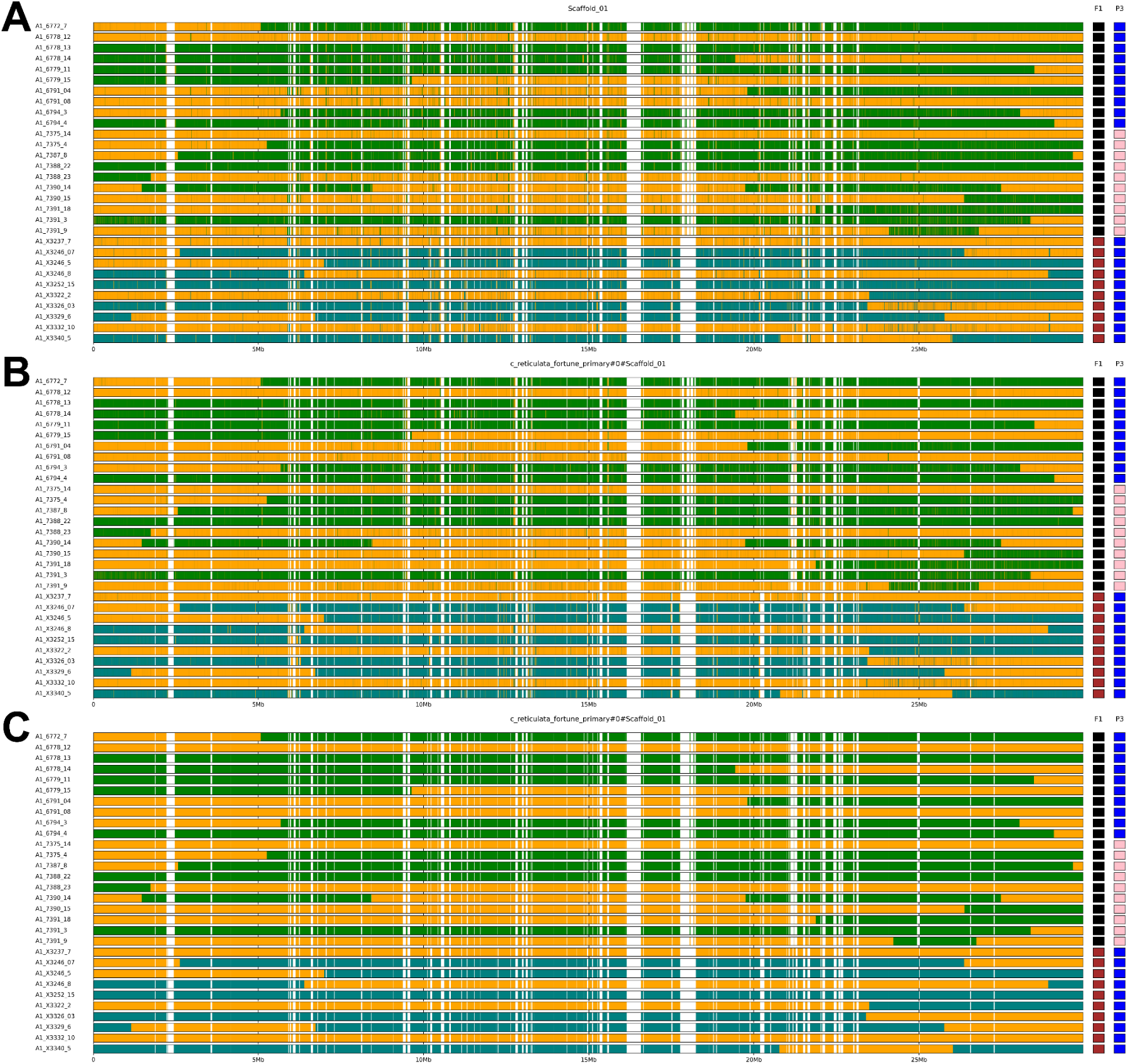
Post-masking parental haplotypes reconstructed from **(A)** linear-alignment based variants, **(B)** graph-alignment based variants, and **(C)** smoothed graph-alignment based variants. Ten subsets of each of the three hybrid families studied are denoted with the F1 parent (black = 475_01; brown = 246_01) and P3 outcrossed parent (blue = ‘Algerian Clementine’; pink = ‘Kiyomi Tangor’). Orange blocks indicate the presence of a mandarin grandparent source and the Australian lime contribution is indicated by either green (*C. australasica*) or dark teal (*C. australis*).

## Discussion

Pangenomes have been widely deployed in human genomics [11], livestock [4,8,34], and crops [7,15,35] but are particularly powerful in interspecific breeding, where genomic complexity is much more prevalent [36]. Interspecific breeding programs enhance crop resilience and genetic diversity by introducing novel traits like disease resistance from related, often undomesticated, species [37]. Increasing variant calling accuracy despite high genomic divergence is necessary for key applications such as QTL mapping. Pangenomes, which incorporate multiple haplotypes across individuals, offer a way to reduce mapping bias and improve the resolution of genomic analysis in breeding contexts.

Previously we have identified sources of citrus huanglongbing resistance in wild Australian limes, spurring the establishment of a breeding program to introgress resistance traits from Australian limes in cultivated citrus [24]. While graph pangenomes are emerging as a promising alternative to traditional linear reference genomes, their reliability for SNP calling, especially in non-model, highly diverged interspecific species, have yet to be reliably characterized. Here, we developed a breeding program-specific pangenome to facilitate analysis of complex hybrids generated and lead to selection of promising individuals in the breeding population. We evaluated this approach for SNP calling by comparing traditional linear, graph-based, and hybrid surjection approaches using F1 and advanced hybrids spanning a diverse lineage of *Citrus* species. One of the central motivations for implementing pangenomes is the expectation of reducing reference bias during read alignment, thus improving downstream genotype accuracy. Consistent with previous studies [3,4,8], alignment to our Minigraph-Cactus super-pangenome using vg giraffe produced a more balanced allele representation than aligning to a linear reference genome using BWA-MEM (**Figure 6**). This suggests that graph pangenomes can more effectively represent aligned reads over linear genomes.

SNP calling using F1s and advanced hybrids revealed substantial variability across the different variant calling approaches, highlighting the sensitivity of SNP discovery in both linear and graph-based representations and calling strategy. Linear and hybrid surjection-based approaches with bcftools and GATK showed high agreement in both SNP positions and genotypes, suggesting that these methods largely recover a shared set of variants. However, linear reference-based BCFtools and GATK consistently identified a larger amount of unique SNPs compared to the hybrid surjection and graph-based approaches. While DeepVariant shared some of the same trends, its results had higher MIER, lower overall SNP count, and lower concordance with other callers. We used the default DeepVariant model trained on human data which is likely not suitable for this type of highly divergent SNP calling analysis without lineage-specific training [38]. In contrast, the purely graph based approach, vg-pack-and-call, produced the fewest SNPs with the highest proportion of missing genotypes compared to the other callers. The vg call tool is built and tested primarily to call SVs, but despite this, the SNPs identified by vg-pack-and-call produced a highly reliable callset by displaying the lowest Mendelian Error Rate (MIER) out of all approaches. This trade-off suggests that vg-pack-and-call is too conservative to capture sufficient SNP variants for most applications, but future graph-based SNP callers have strong potential for improved accuracy.

Graph clipping in our super-pangenome varied across haplotypes. Clipping was more pronounced in haplotypes derived from Australian limes, which are more distantly related to the backbone genome ‘Fortune’, suggesting that these regions were difficult to reliably anchor to the super-pangenome. We also observed that clipping occurred to a similar extent in both primary and alternate haplotype assemblies of the same individual, indicating that clipping is largely driven by sequence similarity rather than haplotype-specific features. Taken together, these results highlight a fundamental trade-off between graph completeness and the ability to reliably utilize pangenomes for tractable downstream analysis. Clipping is currently a necessary computational step for genotyping, but keeping diverged haplotype regions in the graph would improve mapping and genotyping. This could be accomplished by more efficient graphs and read mapping strategies that require no or less clipping. Another promising strategy is to include clipped regions as “bait”, outside the graph structure but still available for read mapping [39]. This strategy gains the advantages of removing reads likely to mismap in the graph while maintaining a computationally tractable graph structure (**Figure 12**).

**Figure 12.**
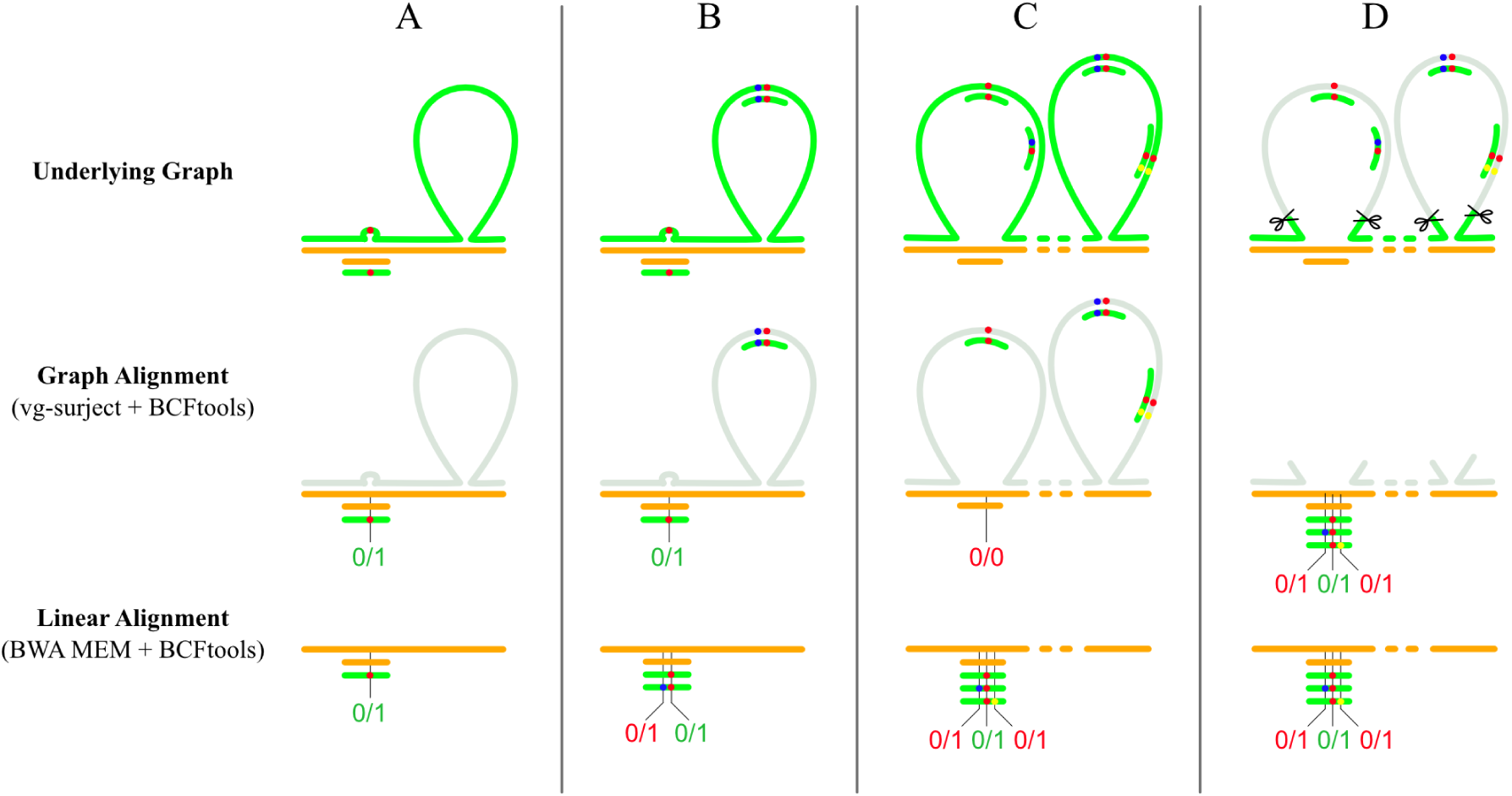
Schematic of hypothetical read mapping scenarios involving hybrid parental reads derived from either of two parental genomes (green or orange). Diverged regions that do not align are visualized as green “bubbles” (top). Where reads cannot be surjected to the reference, those are visualized as grey “bubbles” (middle). SNPs are represented as dots within reads. **(Column A)** SNP calls derived from accurate read mapping to a single origin represented in the full graph (top row), after surjection (middle row) and from linear alignment (bottom row). **(Column B)** Reads that originate from an insertion (green), which would map correctly to the graph (top) and be lost during surjection (middle), may mismap in the linear reference (bottom) to produce incorrect heterozygous calls. **(Column C)** Reads align to the correct target in the full graph. Surjection to a private allele in the reference produces false homozygous interpretation of this locus. Mismapping to the linear reference also creates false calls. **(Column D)** Clipping in the graph (scissors delineate clipped regions) produces a similar set of false heterozygous calls similar to linear B,C,D.

Initial benchmarking assessments of the advanced hybrids suggested that linear-based SNPs outperformed graph-based SNPs due to the 34% increased yield in calls and fewer missing reads, despite only marginal differences in observed MIER. Further investigation into localized regions of error showed that linear-based calls frequently produce erroneous heterozygous calls genome-wide. This phenomenon has been previously documented in populations of closely related *Arabidopsis thaliana* accessions, and these “pseudoheterozygous” calls are likely due to copy number variation and repetitive elements that occur in variable quantities between the reference genome and reads from individual genomes that are aligned to it [40,41]. In contrast, the erroneous graph-based calls occurred in clusters corresponding to the inherent structure of the graph where parental lineages were absent from the selected mandarin parent used for alignment surjection, providing additional context for identifying regions in the reference that are generally problematic for variant calling.

In interspecific hybrid crosses where reads from one parental haplotype can map to the reference but the other cannot due to structural variation, SNP-based genotype calls can produce misleading conclusions about the actual genomic content present (**Figure 12**). In these scenarios, the graph appears to be working as intended by providing recipient sites for reads from expected divergences in lineage; however, the necessity to “surject” or pin those reads back to a fixed reference leads to a flawed interpretation. Specifically, when only one parental haplotype can be observed, it is frequently miscalled as homozygous.

The choice of ‘Fortune’ mandarin as the pangenome backbone raises important considerations regarding the influence of the reference path on SNP calling, particularly for highly diverged taxa such as the Australian limes. While the underlying graph topology remains consistent and thus, the total genetic variation contained within the pangenome graph is preserved, the surjection process inherently anchors observations to the ‘Fortune’ coordinate system. Utilizing an Australian lime species as the reference backbone of the pangenome graph would shift the physical coordinate system of the graph, but the fundamental challenge of reconciling highly diverged regions would remain. We propose that the selection of backbone serves primarily as a coordinate system for the projection of SNPs via surjections, rather than a constraint on the topology of the graph itself. Our masking strategy provides a solution to this challenge by identifying and filtering unreliable regions where the reference path lacks homology with the parental lineages, effectively neutralizing the impact of the backbone choice on genotyping accuracy.

While discarding significant amounts of data may seem counterintuitive, we propose masking these unreliable regions where the selected reference path does not share a degree of homology with the parents used in a breeding program. The improved MIER of both the linear and graph-based calls upon masking suggests that regions difficult to reconcile between assemblies in the pangenome construction may also be regions that are generally challenging for read alignment. The nearly identical SNP counts in both approaches post-masking indicates that regions where founder lines were absent in the reference path may be preferential targets for linear-based alignments and calls, whose reference lacks appropriate recipient targets for reads. Conversely, in the graph, parental calls in regions traversing off-reference “bubbles” may correctly be called as missing, while hybrid progeny from these parents may be incorrectly called by downstream tools where only the parent that is genetically similar to the reference can be mapped (**Figure 12**).

While MIER is a practical benchmarking metric in lieu of a ground-truth SNP reference, the behavior of read-mapping in both the linear and graph-based alignments is susceptible to producing false genotype calls that appear correct under Mendelian assumptions. Examination of unexpected haplotype block transitions that violate expected recombination rates indicates that hybridization between divergent lineages may present unexpected challenges. In the linear alignment, calls consistent with Mendelian inheritance can take the form of reads that multi-map to non-reference accessions mapping to a singular location in the reference path. Similarly, in the graph-based calls, reads that belong to clipped regions will map to unintended targets. In our hybrid crosses, the Australian lime parents are commonly found to be homozygous alternate at haplotype-distinguishing sites. If these sites result from off-target alignments, even from distant locations in the lime genomes, the same pattern will also be present in the progeny which is undetectable using the MIER metric. Although we did not explicitly filter reads with secondary mappings to the chosen reference, doing so can only remove reads that map to multiple locations in the reference path without taking into account reads that originate from multiple sources in the non-reference assembly. Further, removing multi-mapped reads may produce unintended genotypes if only one of the two haplotypes in a cross contains secondary alignments at a locus. A potential approach may be to first catalog reads per founder line with secondary alignments or that map to clipped regions and use this information to further mask VCFs, though this has not been thoroughly tested.

Overall, graph-based alignments combined with standard variant calling protocols appear to provide valuable context, particularly in interspecific crosses. The graph itself provides critical information for identifying reliable regions of the genome for SNP-calling, but additional considerations must be taken into account to avoid misinterpreting variants from highly diverged founder lines.

## Methods

### Phylogenetic Tree

The input set comprised six haplotype assemblies from mandarin (*Citrus reticulata* Blanco) cultivars (‘Fortune’, ‘Fallglo’, and ‘Wilking’) (Singh et al., unpublished data) and six haplotype assemblies from wild Australian lime varieties (*C. australasica* F. Muell., *C. australis* (A.Cunn. ex Mudie) Planch., and *C. inodora* F.M.Bailey) (Table S1) [25], all of which are hosted at the Citrus Genome Database [42]. The primary haplotype from each of the six genomes plus genomes from *C. mangshenensis* S.W. He and *C. trifoliata* L. were provided to OrthoGarden (f9b79e2), which automates extraction of single copy orthogroups to produce a maximum likelihood tree [43].

### Super-Pangenome Construction and Metrics

To capture the genetic variation within our breeding program, we constructed a graph-based *Citrus* super-pangenome consisting of 12 haplotype-resolved, chromosome-scale assemblies from six diploid genome assemblies described above (6 genomes x 2 haplotypes). The pangenome was built using the Minigraph-Cactus (MC) pipeline version 2.9.3 [9], treating wild and cultivated varieties as distinct inputs [9]. As MC requires a linear reference as a backbone for anchoring the graph, we selected the ‘Fortune’ primary haplotype due to its slightly higher contiguity and genome completeness relative to the other assemblies available in our breeding program. The remaining 11 haplotypes were then aligned iteratively to this reference backbone to construct a super-pangenome graph. For all downstream analyses, we used the reduced (clipped) pangenome graph to minimize structural complexity and maintain computational tractability.

Graph level metrics and summaries were generated using ODGI (version 0.9.2)[44]. Basic statistics, including node count, edge count, and path metrics, were obtained via odgi stats using the super-pangenome GFA as input. Cumulative graph growth and node sharing patterns across haplotypes were evaluated and visualized using Panacus (version 0.4.1) [30]. Panacus was run on the same GFA using the histgrowth module with haplotype level grouping enabled and coverage quorum thresholds of 1, 2, 3, and 12, corresponding to different levels of node agreement. This produced node presence tables summarizing how the super-pangenome expanded as haplotypes were added and provided the basis of node classification into core, dispensable, and private categories.

### Graph Clipping

To identify regions that were clipped during pangenome construction, we extracted all walk lines, which describe a haplotype’s paths in GFA format 1.1, from ODGI’s representation of the super-pangenome in .og format. Walk lines were parsed to obtain coordinate intervals for each walk line of every haplotype. These intervals were converted to BED format and separated into one file per haplotype, which represented all continuous genome segments that are present when using a reduced (clipped) MC super-pangenome. For every haplotype, the BED intervals representing non-clipped sequences were first sorted and merged using BEDtools (version 2.31.1) [45] to ensure any overlapping regions were represented as continuous intervals. To identify clipped regions, BEDtools was used to complement, supplying the non-clipped BED file together with a haplotype specific genome length file generated from samtools (version 1.21) [46]. This returned all genomic intervals that were not present in the non-clipped set, corresponding to regions removed during reduced (clipped) MC construction.

Because variant calling was performed relative to the ‘Fortune’ primary haplotype, we converted each haplotype’s clipped regions to the backbone coordinates. We used ODGI extract to walk paths from each haplotype, generating an ordered list of sequence graph nodes. For the ‘Fortune’ primary backbone, this provided the complete ordered list of backbone nodes associated with their respective genomic coordinates. For each non-reference haplotype, each ODGI walk file was parsed to identify all backbone nodes present in each haplotype’s path. We identified shared nodes by subtracting these node intervals from the backbone’s, which defined the precise regions that were not supported by each haplotype. These unsupported regions were visualized as per-haplotype scaffold track plots. In addition, unsupported regions were summarized across all haplotypes using 100 kb backbone windows and visualized as a multi-haplotype heatmap. Data for these plots were processed using pandas (2.2.3), and visualizations were generated using matplotlib (3.10.3).

### Sequencing

Citrus accessions used as parents in the breeding program were sampled from the Givaudan Citrus Variety Collection at the University of California Riverside (UCR) and the hybrids generated were collected from the UCR experimental fields [47,48]. Illumina sequencing was performed on eight parental lines, 30 F1 hybrids and 244 advanced hybrids (**Supplemental Table 2**) using 150 bp paired-end reads at Novogene Corporation (Davis, California, USA). In addition to the six parental lines used for the pangenome above, two domesticated *Citrus* used as advanced hybrid parents were included in the sequencing: *C. reticulata* ‘Algerian Clementine’ and ‘Kiyomi Tangor-H12’. After adapter trimming with fastp [49], the average read length for the samples was 147 bp. Sequences had an average depth of 39x, with a range between 23.54x and 57.45x across the F1 samples based off of the length of the ‘Fortune’ primary assembly, while the advanced hybrids (AD) had an average depth of 47.9x, ranging from 39.9x to 69.1x.

### F1 SNP Calling

To align short reads, we implemented two strategies representing contrasting approaches: (1) a traditional linear reference-based approach and (2) a graph-based pangenome approach. For the linear reference, reads were aligned to the ‘Fortune’ primary haplotype assembly using BWA-MEM (version 0.7.18) [50]. For the pangenome-based method, reads were aligned to the MC super-pangenome using vg giraffe (version 1.64.0) [3].

Following alignment, variant calling was performed separately for each strategy. For linear alignments, three SNP calling approaches were used: GATK HaplotypeCaller (version 4.6.0.0) [51], DeepVariant (version 1.8.0) [52], and BCFtools (version 1.21.0) [46]. HaplotypeCaller was run on individual samples in GVCF mode, and joint genotyping was performed using GenotypeGVCFs. DeepVariant was run using the default whole genome sequencing model, and the resulting calls were joint-genotyped using GLnexus (version 1.4.1) [53]. For BCFtools, variant calling was performed using mpileup with read and allele depth annotations enabled, followed by bcftools call in multiallelic mode. BCFtools calls were finalized by concatenating per-sample VCFs using bcftools concat.

For the graph-based approach, reads were aligned to the MC super-pangenome using vg giraffe, and variant calling was performed using the vg toolkit. Specifically, alignments were converted to pack files using vg pack, and SNPs were called using vg call. The resulting sample VCFs were merged, normalized, and sorted using bcftools merge, norm, and sort.

In addition, implemented a hybrid approach to leverage a pangenome’s broader variant representation with compatibility to traditional callers. In this approach, reads were aligned to the super-pangenome with vg giraffe. Next, vg-surject was used to surject the alignments onto the ‘Fortune’ primary reference path within the super-pangenome to produce coordinate-compatible BAM files. These surjected alignments were subsequently used as input in BCFtools and GATK HaplotypeCaller, following the same practices as the linear approach. For all SNP calling approaches using the linear, graph-based, and hybrid methods, the multisample VCF file was filtered to retain only biallelic SNPs with a PASS filter and a QUAL score ≥ 30.

### Shared SNPs and Genotype Comparison

To compare genotypes across the different callers, we first identified shared SNPs that were called by all approaches. For each caller, SNP position (chromosome:position) was extracted from each VCF file using bcftools query to produce positional lists. Shared SNPs from all callers were merged into a master list and sorted based on position. This list was converted into a presence/absence matrix, where each row represents a SNP and each column represents a variant caller. Cells were marked “True” if a caller contained that SNP and “False” otherwise. By filtering this matrix to retain only rows where all callers are “True” values, we identified SNPs that were shared across all callers. SNP overlap across the six approaches was visualized using Python’s UpSet plot library.

Next, we compared genotype calls using the shared list of SNPs. For each caller, VCF files were subset to only include shared SNPs using bcftools view. Genotype data was extracted from each sample using bcftools query. These genotype tables were then merged into a single table. All genotype calls were normalized by converting phased (|) to unphased (/) and sorting alleles (1/0 to 0/1).

### Genotype Accuracy Benchmarking using Mendelian Inheritance

Mendelian Inheritance Error Rate (MIER) was calculated for every parent-progeny trio in the dataset. Among complete trios, a progeny genotype was flagged as erroneous if it violated Mendelian inheritance: homozygous reference (“0/0”) progeny when any parent was homozygous alternate (“1/1”), homozygous alternate progeny when any parent was homozygous reference, or heterozygous progeny when both parents were identically homozygous. Unless otherwise specified, calls missing for any members of the trio were not included as errors because a missing call could be the desirable behavior in interspecific crosses.

### Reference Bias Estimate

To assess reference bias across our different methods, we measured allele balance at putative heterozygous SNP sites following the approach of Rice et al. (2023) [8]. Sites were defined as heterozygous if they had a minimum depth of 10x and a minor allele frequency of at least 25%. For each site, we calculated the fraction of reads supporting the alternate allele, with an expected unbiased representation of 0.5. Mean alternate allele fractions were then computed per sample for both linear (BWA-MEM) and the pangenome (vg giraffe) alignments.

### Graph Clipping Correlation with MIER

For F1 hybrids grouped by their respective parents (*C. inodora, C. australis,* and *C. australasica)*, we analyzed SNPs derived from vg-surject BCFtools. For each parental group, SNPs were subsetted to sites with a genotype missingness rate of <20% and a minor allele frequency >10%. Previous computed MIER values for these SNPs were converted to bed format using awk. We then used bedtools intersect to associate MIER values with graph regions classified by the number of clipped Australian lime haplotypes (zero, one, or two). To test whether MIER differed amongst these clipped categories, we performed a Kruskal-Wallis test by ranks using the scipy.stats Python library. For pairwise comparisons, 10,000 SNPs were used (randomly downsampled) per group and Dunn’s post-hoc test was used via scikit_posthocs Python library.

### Advanced Hybrids (AD): SNP Calling and MIER Benchmarking

For the advanced hybrid sequencing, two of the above SNP calling pipelines were used: BWA-MEM with BCFtools (“BCFtools”) and vg giraffe to vg surject with BCFtools (“vg-surject-BCFtools”). MIER was calculated using the same script for the advanced hybrids. For each of the two methods, for 100 kb windows across the genome, we calculated rates of incorrect homozygous reference calls, incorrect heterozygous calls, incorrect homozygous alternate calls, incorrect total calls, correct calls, and missing calls.

Additional analyses were performed per SNP to assess correlation of regional graph structure and calling performance. For each node in the GFA, the genomic coordinates for the reference path (*C. reticulata* ‘Fortune’ primary assembly) were obtained as well as the paths traversing reference nodes. Using defined 150-bp subpaths up and downstream from each SNP coordinate along this path, the presence of each founder line was calculated as the percent of base pairs occupied in the subpath of each founder (i.e., the founder path shared those nodes with the reference). Spearman correlation was performed using the founder presence to the MIER-derived statistics to calculate significance. Average presence for individual founders was calculated and visualized across 100 kb blocks using matplotlib [54].

### Advanced Hybrids (AD): Haplotype Blocks

In order to further evaluate the applicability of our variant calls from both graph-based and linear approaches, we attempted to reconstruct the parental contribution of the F1 (mandarin x Australian lime). The F1 haplotype inherited in the advanced hybrid progeny was predicted on a per-trio basis using sites that met Mendelian inheritance pattern expectations in the F1 trio as well as the advanced hybrid trio. SNPs were then further filtered to those sites where F1 parental genotypes were opposing homozygous calls (0/0 x 1/1 or 1/1 x 0/0). The ancestry of the alleles contributing to the F1 types (0/1) was traced via the second generation of “advanced” hybrids (outcrossed to other domesticated mandarin cultivars not present in the graph) using sites where the outcrossing parent was homozygous. These are referred to as “diagnostic SNPs”, which followed Mendelian patterns of inheritance over two generations. Diagnostic SNPs were used to infer the mandarin or Australian lime contribution inherited from the F1, and these typed positions were plotted per advanced hybrid to represent haplotype blocks.

### Advanced Hybrids (AD): Smoothing

Haplotype blocks predicted from masked, graph-based SNP calls were used to predict recombination events in the advanced hybrid population. To smooth the predictions and remove noise, the parental types of diagnostic SNPs in a window of 5000 up and downstream from each site were used to median filter calls that mismatch the local haplotype context. To avoid edge effects near telomeres, this window was scaled such that the flanking regions were equal in SNP number and regions within 500 calls to the end were processed in batch. Based on the recombination loci from the smoothed blocks, the recombination rate was calculated as the average count of crossing over events (2206) for the 244 advanced hybrids in cM (904.1cM) divided by the total chromosome scaffold size for the reference genome in Mb.

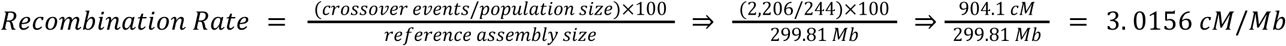

Diagnostic SNPs were then compared against the smoothed haplotype blocks and were determined to be discordant when not in agreement with the local context.

### Advanced Hybrids (AD): Masking

The presence-absence of each founder line for each node in the reference genome path was obtained as described above. Nodes missing in both haplotypes of the F1 parental lines for the hybrid families were used as coordinates for masking based on a minimum consecutive length of missing parental contribution. A parameter sweep of consecutive lengths ranging from 2 to 1024 bp were evaluated. While lengths as low as 3 bp segregated MIER incorrect calls the most consistently, we opted for a minimum length of 6 bp, as it removed the greatest proportion of reference bias, specifically MIER incorrect calls of homozygous reference. Bedtools intersect (v 2.31.1) was used to split the VCF files into the masked and unmasked regions, and the resulting files were calculated using the same MIER summary statistics and haplotype block approaches described above.

## Data availability

Reference assemblies used in this publication can be found at the Citrus Genome Database (https://www.citrusgenomedb.org/). All short read data for parental lines and progeny have been deposited at NCBI SRA under project PRJXXXXXXX. Code and scripts used for MIER analyses are available at https://github.com/ryandkuster/SNP_mendelian_inheritance_citrus.

## Acknowledgements

Funding

This work was supported by USDA NIFA ECDRE grant 2023-70029-41315 and USDA ARS CA 59-8080-0-004.

**Supplemental Figures 1-11.** Chromosome scale clipping for haplotypes in the MC pangenome.

**Supplemental Figures 12-20.** Summary comparison of genotype call types from linear-BCFtools (solid line) and vg-surject-BCFtools (dotted line) across 100 kb windows on chromosomes 1-9.

**Supplemental Figures 21-29.** Relationship of founder presence in the graph nodes per 100 kb windows along chromosomes 1-9.

**Supplemental Figure 30.** Representation of a single chromosome in the F1 and advanced hybrid breeding program.

**Supplemental Figures 31-39.** Recombination frequencies on chromosomes 1-9 derived from parental haplotypes reconstruction in 244 advanced hybrids.

**Supplemental Table 1.** Genome statistics and public availability.

**Supplemental Table 2.** Illumina read statistics and public availability.

**Supplemental Table 3.** Comparison of genotype frequency and average depth in regions of the masked vcf (mask) vs. regions only within masking ranges (mask inverse).

## References

1. Brandt DYC, Aguiar VRC, Bitarello BD, Nunes K, Goudet J, Meyer D. Mapping Bias Overestimates Reference Allele Frequencies at the *HLA* Genes in the 1000 Genomes Project Phase I Data. G3 Genes|Genomes|Genetics. 2015;5:931–41. 10.1534/g3.114.015784

2. Ballouz S, Dobin A, Gillis JA. Is it time to change the reference genome? Genome Biol. 2019;20:159. 10.1186/s13059-019-1774-4

3. Sirén J, Monlong J, Chang X, Novak AM, Eizenga JM, Markello C, et al. Pangenomics enables genotyping of known structural variants in 5202 diverse genomes. Science. American Association for the Advancement of Science; 2021;374:abg8871. 10.1126/science.abg8871

4. Miao J, Wang Q, Zhang Z, Wang Q, Pan Y, Wang Z. Pangenome graph mitigates heterozygosity overestimation from mapping bias: a case study in Chinese indigenous pigs. BMC Biol. 2025;23:89. 10.1186/s12915-025-02194-y

5. Martiniano R, Garrison E, Jones ER, Manica A, Durbin R. Removing reference bias and improving indel calling in ancient DNA data analysis by mapping to a sequence variation graph. Genome Biology. 2020;21:250. 10.1186/s13059-020-02160-7

6. Garrison E, Guarracino A, Heumos S, Villani F, Bao Z, Tattini L, et al. Building pangenome graphs. Nature Methods. 2024;21:2008–12. 10.1038/s41592-024-02430-3

7. Zhou Y, Zhang Z, Bao Z, Li H, Lyu Y, Zan Y, et al. Graph pangenome captures missing heritability and empowers tomato breeding. Nature. 2022;606:527–34. 10.1038/s41586-022-04808-9

8. Rice ES, Alberdi A, Alfieri J, Athrey G, Balacco JR, Bardou P, et al. A pangenome graph reference of 30 chicken genomes allows genotyping of large and complex structural variants. BMC Biology. 2023;21:267. 10.1186/s12915-023-01758-0

9. Hickey G, Monlong J, Ebler J, Novak AM, Eizenga JM, Gao Y, et al. Pangenome graph construction from genome alignments with Minigraph-Cactus. Nature Biotechnology. 2024;42:663–73. 10.1038/s41587-023-01793-w

10. Ahmad B, Su Y, Hao Y, Razzaq T, Arshad R, Zhang Y, et al. Mango pangenome reveals dramatic impacts of reference bias on population genomic analyses. Horticulture Research. 2025;12:uhaf166. 10.1093/hr/uhaf166

11. Liao W-W, Asri M, Ebler J, Doerr D, Haukness M, Hickey G, et al. A draft human pangenome reference. Nature. 2023;617:312–24. 10.1038/s41586-023-05896-x

12. Browning SR, Browning BL. Haplotype phasing: existing methods and new developments. Nat Rev Genet. 2011;12:703–14. 10.1038/nrg3054

13. Nijveen H, Van Kaauwen M, Esselink DG, Hoegen B, Vosman B. QualitySNPng: a user-friendly SNP detection and visualization tool. Nucleic Acids Research. 2013;41:W587–90. 10.1093/nar/gkt333

14. Vanderzande S, Howard NP, Cai L, Da Silva Linge C, Antanaviciute L, Bink MCAM, et al. High-quality, genome-wide SNP genotypic data for pedigreed germplasm of the diploid outbreeding species apple, peach, and sweet cherry through a common workflow. Lightfoot DA, editor. PLoS ONE. 2019;14:e0210928. 10.1371/journal.pone.0210928

15. Cochetel N, Minio A, Guarracino A, Garcia JF, Figueroa-Balderas R, Massonnet M, et al. A super-pangenome of the North American wild grape species. Genome Biology. 2023;24:290. 10.1186/s13059-023-03133-2

16. Raza A, Bohra A, Garg V, Varshney RK. Back to wild relatives for future breeding through super-pangenome. Molecular Plant. 2023;16:1363–5. 10.1016/j.molp.2023.08.005

17. Van Workum D-JM, Mehrem SL, Snoek BL, Alderkamp MC, Lapin D, Mulder FFM, et al. Lactuca super-pangenome reduces bias towards reference genes in lettuce research. BMC Plant Biol. 2024;24:1019. 10.1186/s12870-024-05712-2

18. Leonard AS, Crysnanto D, Mapel XM, Bhati M, Pausch H. Graph construction method impacts variation representation and analyses in a bovine super-pangenome. Genome Biol. 2023;24:124. 10.1186/s13059-023-02969-y

19. Alonge M, Wang X, Benoit M, Soyk S, Pereira L, Zhang L, et al. Major Impacts of Widespread Structural Variation on Gene Expression and Crop Improvement in Tomato. Cell. 2020;182:145–161.e23. 10.1016/j.cell.2020.05.021

20. Wang B, Hou M, Shi J, Ku L, Song W, Li C, et al. De novo genome assembly and analyses of 12 founder inbred lines provide insights into maize heterosis. Nature Genetics. 2023;55:312–23. 10.1038/s41588-022-01283-w

21. Jayakodi M, Lu Q, Pidon H, Rabanus-Wallace MT, Bayer M, Lux T, et al. Structural variation in the pangenome of wild and domesticated barley. Nature. 2024;636:654–62. 10.1038/s41586-024-08187-1

22. Jiao C, Xie X, Hao C, Chen L, Xie Y, Garg V, et al. Pan-genome bridges wheat structural variations with habitat and breeding. Nature [Internet]. 2024; 10.1038/s41586-024-08277-0

23. Jagoueix S, Bove J-M, Garnier M. The Phloem-Limited Bacterium of Greening Disease of Citrus Is a Member of the Subdivision of the Proteobacteria. International Journal of Systematic Bacteriology. 1994;44:379–86. 10.1099/00207713-44-3-379

24. Ramadugu C, Keremane ML, Halbert SE, Duan YP, Roose ML, Stover E, et al. Long-Term Field Evaluation Reveals Huanglongbing Resistance in *Citrus* Relatives. Plant Disease. 2016;100:1858–69. 10.1094/PDIS-03-16-0271-RE

25. Singh K, Huff M, Liu J, Park J-W, Rickman T, Keremane M, et al. Chromosome-Scale, De Novo, Phased Genome Assemblies of Three Australian Limes: Citrus australasica, C. inodora, and C. glauca. Plants. 2024;13:1460. 10.3390/plants13111460

26. Liu J, Singh K, Huff M, Gottschalk C, Do M, Staton M, et al. Deep R-gene discovery in HLB resistant wild Australian limes uncovers evolutionary features and potentially important loci for hybrid breeding. Front Plant Sci. 2025;15:1503030. 10.3389/fpls.2024.1503030

27. Mahmoud LM, Dutt M. Novel citrus hybrids incorporating Australian lime genetics: development of HLB-tolerant citrus rootstocks and physiological changes in ‘Valencia’ sweet orange scions. Front Plant Sci. 2025;16:1614845. 10.3389/fpls.2025.1614845

28. Wu GA, Terol J, Ibanez V, López-García A, Pérez-Román E, Borredá C, et al. Genomics of the origin and evolution of Citrus. Nature. 2018;554:311–6. 10.1038/nature25447

29. Nakandala U, Furtado A, Masouleh AK, Smith MW, Mason P, Williams DC, et al. The genomes of Australian wild limes. Plant Mol Biol. 2024;114:102. 10.1007/s11103-024-01502-4

30. Parmigiani L, Garrison E, Stoye J, Marschall T, Doerr D. Panacus: fast and exact pangenome growth and core size estimation. Bioinformatics. 2024;40:btae720. 10.1093/bioinformatics/btae720

31. Aleza P, Cuenca J, Hernández M, Juárez J, Navarro L, Ollitrault P. Genetic mapping of centromeres in the nine Citrus clementina chromosomes using half-tetrad analysis and recombination patterns in unreduced and haploid gametes. BMC Plant Biology. 2015;15:80. 10.1186/s12870-015-0464-y

32. Tiley GP, Burleigh JG. The relationship of recombination rate, genome structure, and patterns of molecular evolution across angiosperms. BMC Evolutionary Biology. 2015;15:194. 10.1186/s12862-015-0473-3

33. Diaz IA, Ostovar T, Chen J, de Dios EA, Piscatella R, Perez-Alfaro RS, et al. A haplotype-resolved chromatin landscape connects cis-regulatory variants to trait variation in Citrus. BMC Genomics. 2025;26:978. 10.1186/s12864-025-12137-0

34. Li R, Gong M, Zhang X, Wang F, Liu Z, Zhang L, et al. A sheep pangenome reveals the spectrum of structural variations and their effects on tail phenotypes. Genome Res. United States; 2023;33:463–77. 10.1101/gr.277372.122

35. Golicz AA, Bayer PE, Barker GC, Edger PP, Kim H, Martinez PA, et al. The pangenome of an agronomically important crop plant Brassica oleracea. Nature Communications. 2016;7:13390. 10.1038/ncomms13390

36. Della Coletta R, Qiu Y, Ou S, Hufford MB, Hirsch CN. How the pan-genome is changing crop genomics and improvement. Genome Biology. 2021;22:3. 10.1186/s13059-020-02224-8

37. Kashyap A, Garg P, Tanwar K, Sharma J, Gupta NC, Ha PTT, et al. Strategies for utilization of crop wild relatives in plant breeding programs. Theoretical and Applied Genetics. 2022;135:4151–67. 10.1007/s00122-022-04220-x

38. Sapoval N, Aghazadeh A, Nute MG, Antunes DA, Balaji A, Baraniuk R, et al. Current progress and open challenges for applying deep learning across the biosciences. Nat Commun. 2022;13:1728. 10.1038/s41467-022-29268-7

39. Li H, Feng X, Chu C. The design and construction of reference pangenome graphs with minigraph. Genome Biol. 2020;21:265. 10.1186/s13059-020-02168-z

40. Jaegle B, Pisupati R, Soto-Jiménez LM, Burns R, Rabanal FA, Nordborg M. Extensive sequence duplication in Arabidopsis revealed by pseudo-heterozygosity. Genome Biology. 2023;24:44. 10.1186/s13059-023-02875-3

41. Igolkina AA, Vorbrugg S, Rabanal FA, Liu H-J, Ashkenazy H, Kornienko AE, et al. A comparison of 27 Arabidopsis thaliana genomes and the path toward an unbiased characterization of genetic polymorphism. Nature Genetics [Internet]. 2025; 10.1038/s41588-025-02293-0

42. Main D. Citrus Genome Database [Internet]. 2026. https://www.citrusgenomedb.org/

43. Turner JH, Kuster RD, Staton ME, Moulton JK. OrthoGarden: a pipeline for propagating phylogenetic trees for nonmodel organisms from short reads and de novo genome assemblies. Molecular Biology and Evolution. 2026;43:msag053. 10.1093/molbev/msag053

44. Guarracino A, Heumos S, Nahnsen S, Prins P, Garrison E. ODGI: understanding pangenome graphs. Bioinformatics. 2022;38:3319–26. 10.1093/bioinformatics/btac308

45. Quinlan AR, Hall IM. BEDTools: a flexible suite of utilities for comparing genomic features. Bioinformatics. 2010;26:841–2. 10.1093/bioinformatics/btq033

46. Danecek P, Bonfield JK, Liddle J, Marshall J, Ohan V, Pollard MO, et al. Twelve years of SAMtools and BCFtools. GigaScience. 2021;10:giab008. 10.1093/gigascience/giab008

47. Ramadugu C, Keremane M, McCollum TG, Hall DG, Roose ML. Developing resistance to HLB. Citrograph. 2016;7:46–51.

48. Ramadugu C, Keremane M, Lee RF, Hall DG, McCollum T G, Roose ML. Novel citrus hybrids with HLB resistance. Citrograph. 2019;10:60–4.

49. Chen S, Zhou Y, Chen Y, Gu J. fastp: an ultra-fast all-in-one FASTQ preprocessor. Bioinformatics. 2018;34:i884–90. 10.1093/bioinformatics/bty560

50. Li H. Aligning sequence reads, clone sequences and assembly contigs with BWA-MEM [Internet]. 2013. https://arxiv.org/abs/1303.3997

51. Poplin R, Ruano-Rubio V, DePristo MA, Fennell TJ, Carneiro MO, Van der Auwera GA, et al. Scaling accurate genetic variant discovery to tens of thousands of samples. bioRxiv. 2018;201178. 10.1101/201178

52. Poplin R, Chang P-C, Alexander D, Schwartz S, Colthurst T, Ku A, et al. A universal SNP and small-indel variant caller using deep neural networks. Nature Biotechnology. 2018;36:983–7. 10.1038/nbt.4235

53. Yun T, Li H, Chang P-C, Lin MF, Carroll A, McLean CY. Accurate, scalable cohort variant calls using DeepVariant and GLnexus. Bioinformatics. England; 2021;36:5582–9. 10.1093/bioinformatics/btaa1081

54. Hunter JD. Matplotlib: A 2D Graphics Environment. Comput Sci Eng. 2007;9:90–5. 10.1109/MCSE.2007.55

