## SupplementalFigures for "Benchmarking SNP-Calling Accuracy Against Known *Citrus* Pedigrees Reveals Pangenome Advantages Over Linear References"

### Supplemental Figures

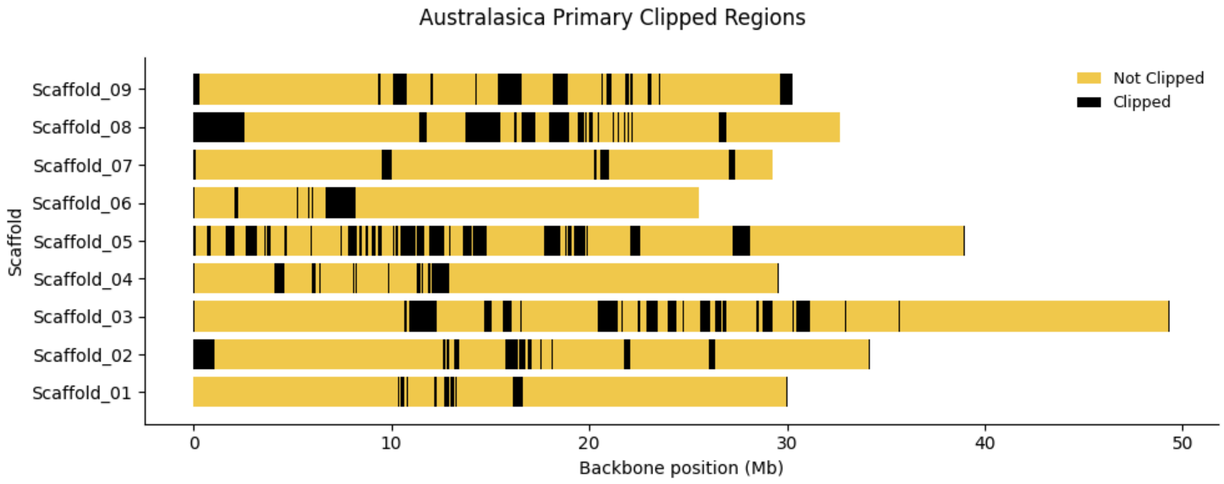


**Figure S1.** Clipped sequences in the *C. australasica* primary haplotype.

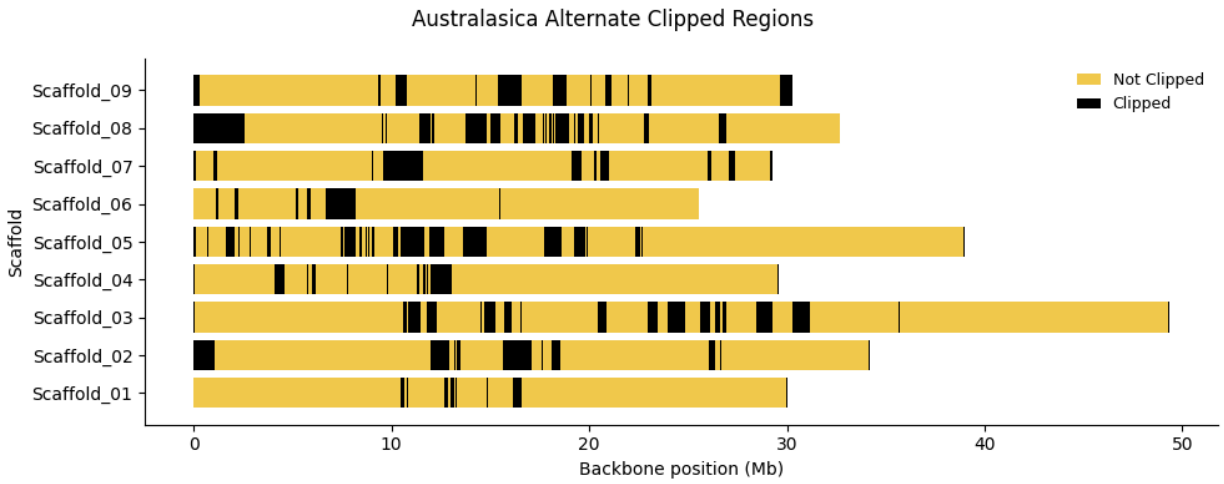


**Figure S2.** Clipped sequences in the *C. australasica* alternate haplotype.

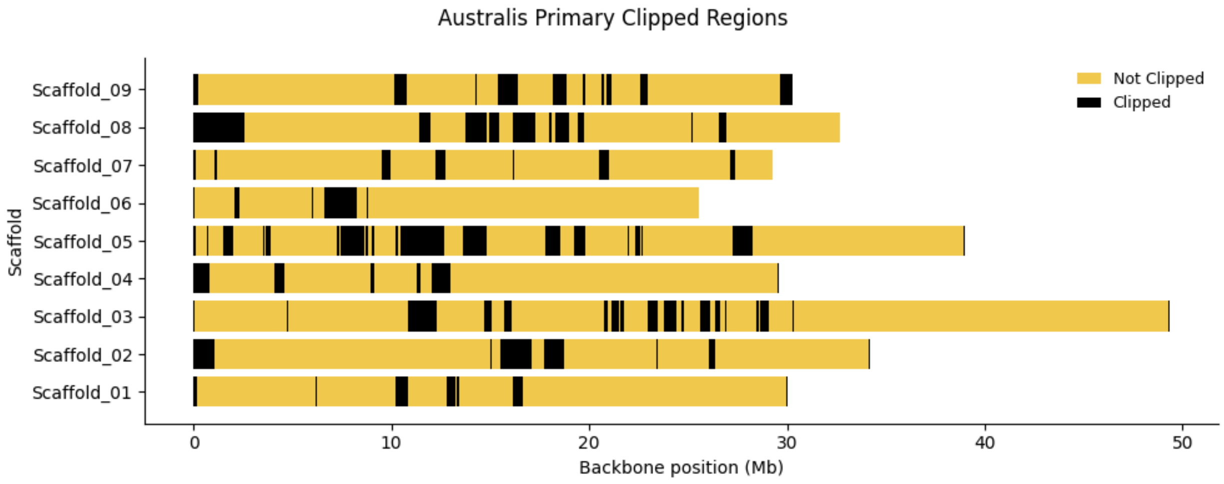


**Figure S3.** Clipped sequences in the *C. australis* primary haplotype.

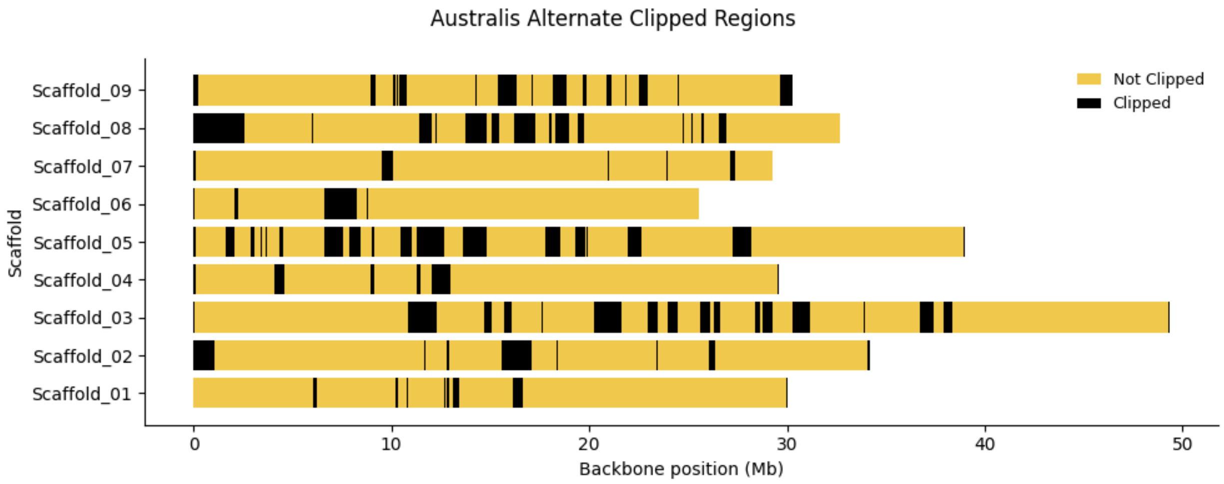


**Figure S4.** Clipped sequences in the *C. australis* alternate haplotype.

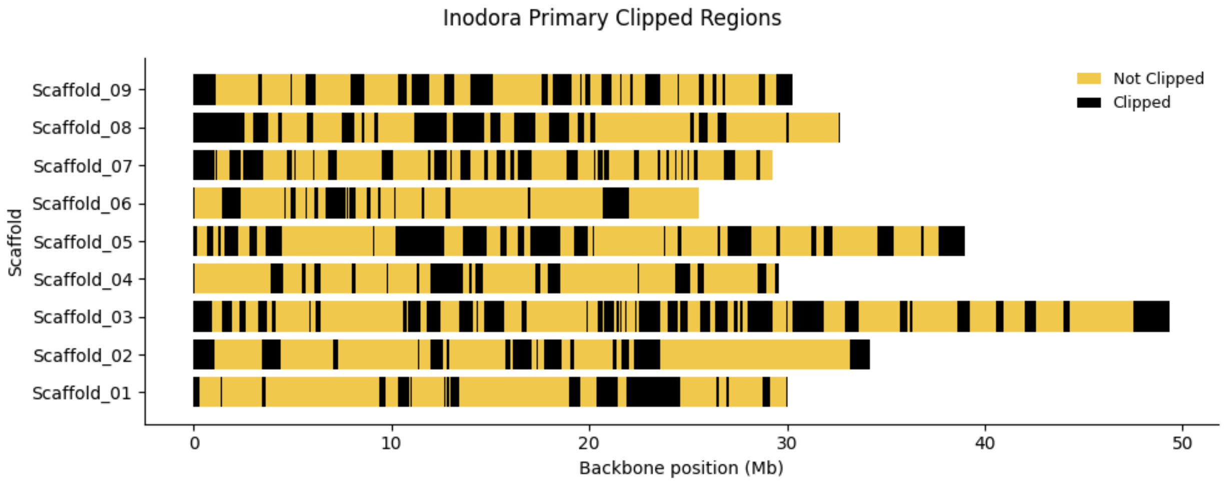


**Figure S5.** Clipped sequences in the *C. inodora* primary haplotype.

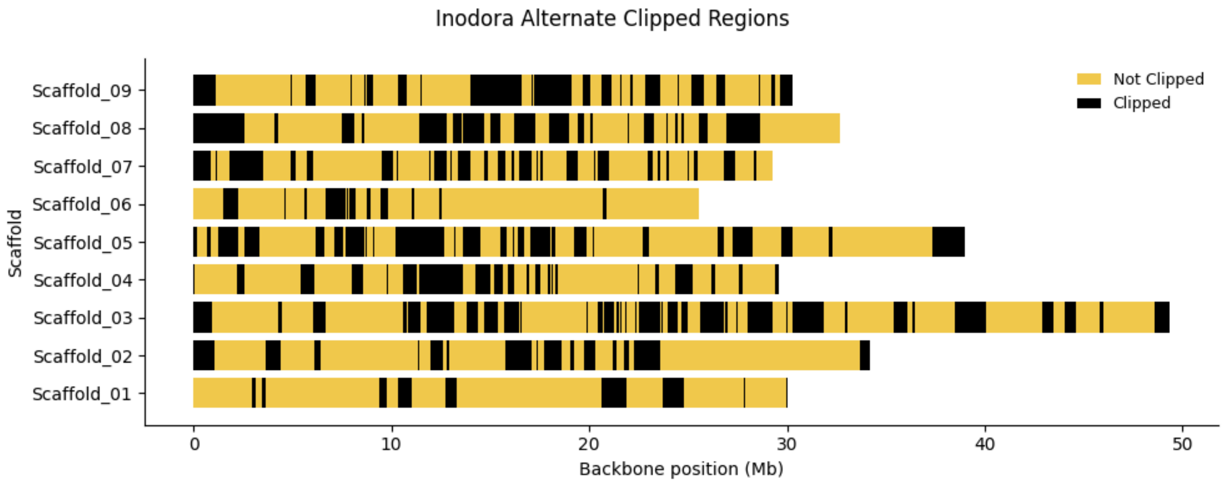


**Figure S6.** Clipped sequences in the *C. inodora* alternate haplotype.

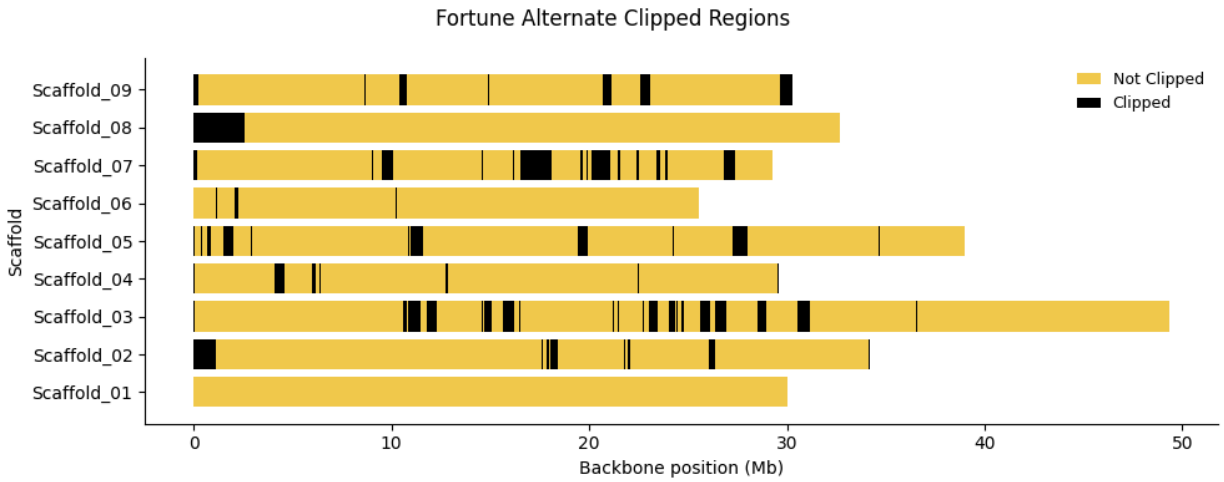


**Figure S7.** Clipped sequences in the *C. reticulata* ‘Fortune’ alternate haplotype.

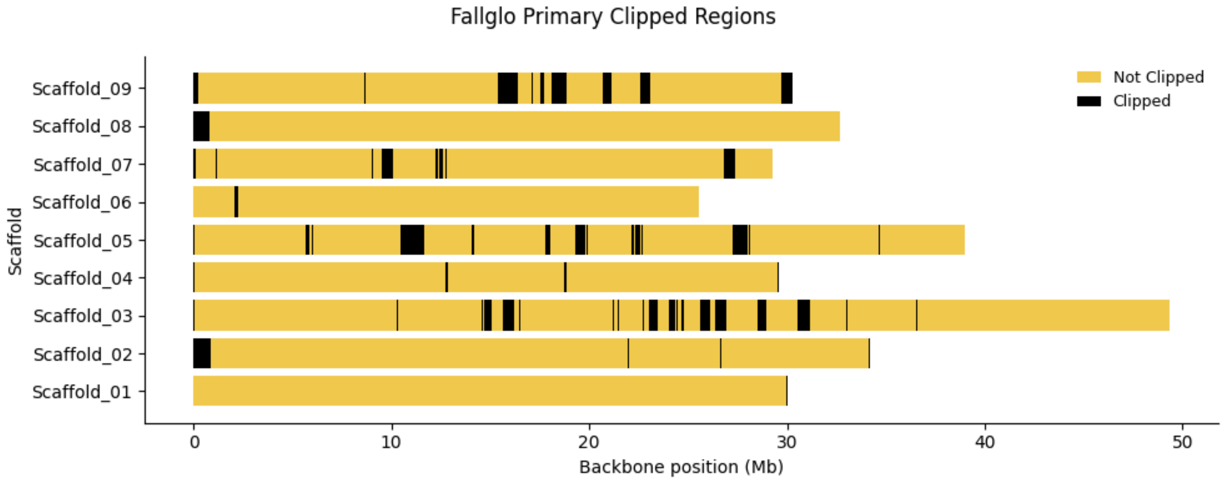


**Figure S8.** Clipped sequences in the *C. reticulata* ‘Fallglo’ primary haplotype.

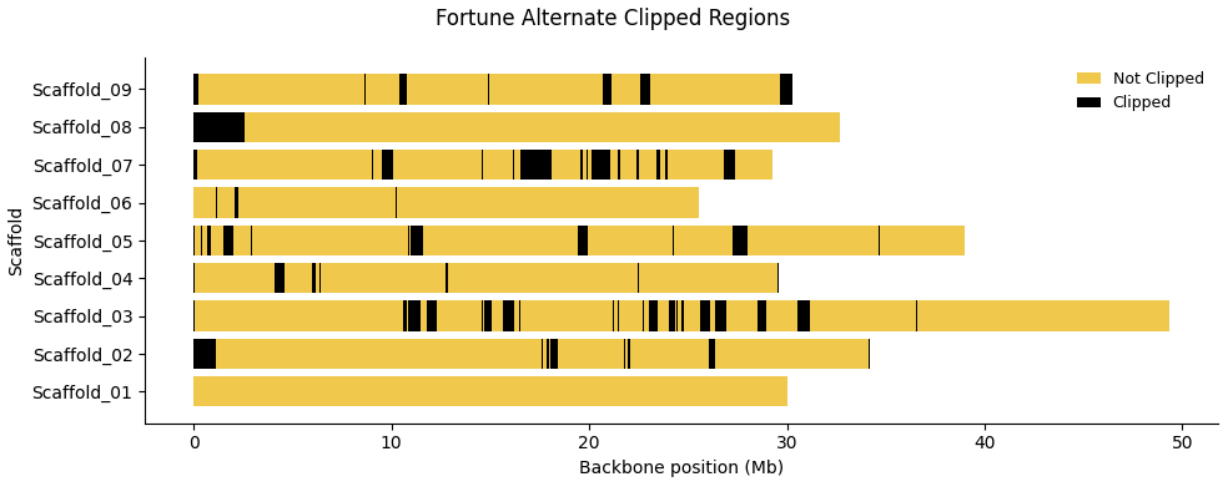


**Figure S9.** Clipped sequences in the *C. reticulata* ‘Fallglo’ alternate haplotype.

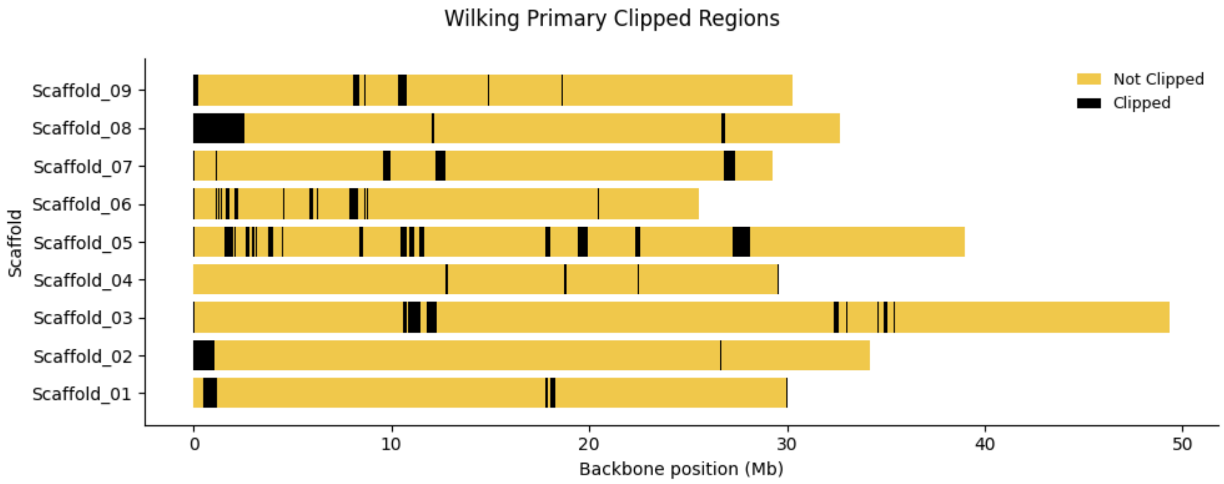


**Figure S10.** Clipped sequences in the *C. reticulata* ‘Wilking’ primary haplotype.

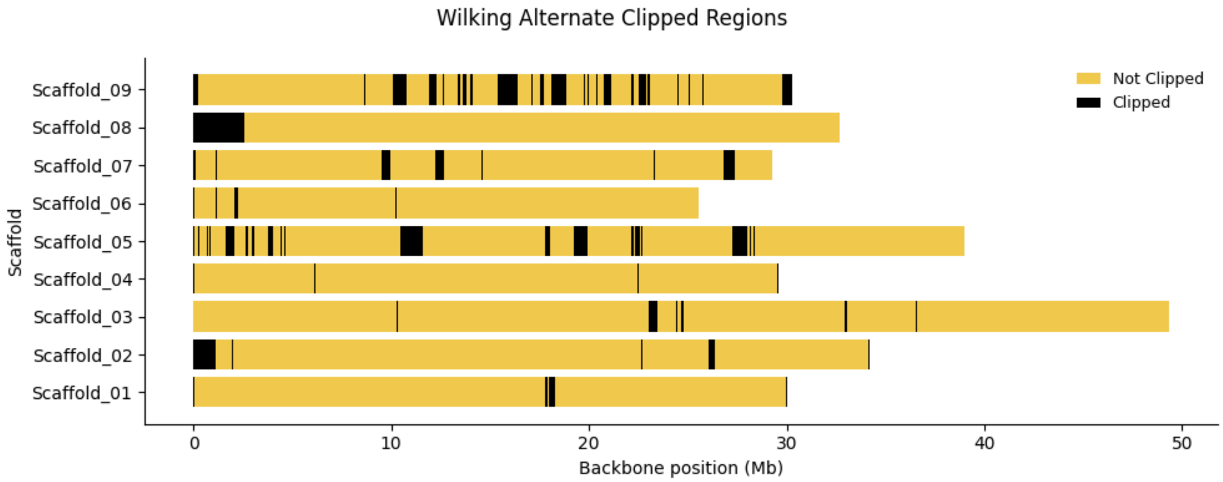


**Figure S11.** Clipped sequences in the *C. reticulata* 'Wilking' alternate haplotype.

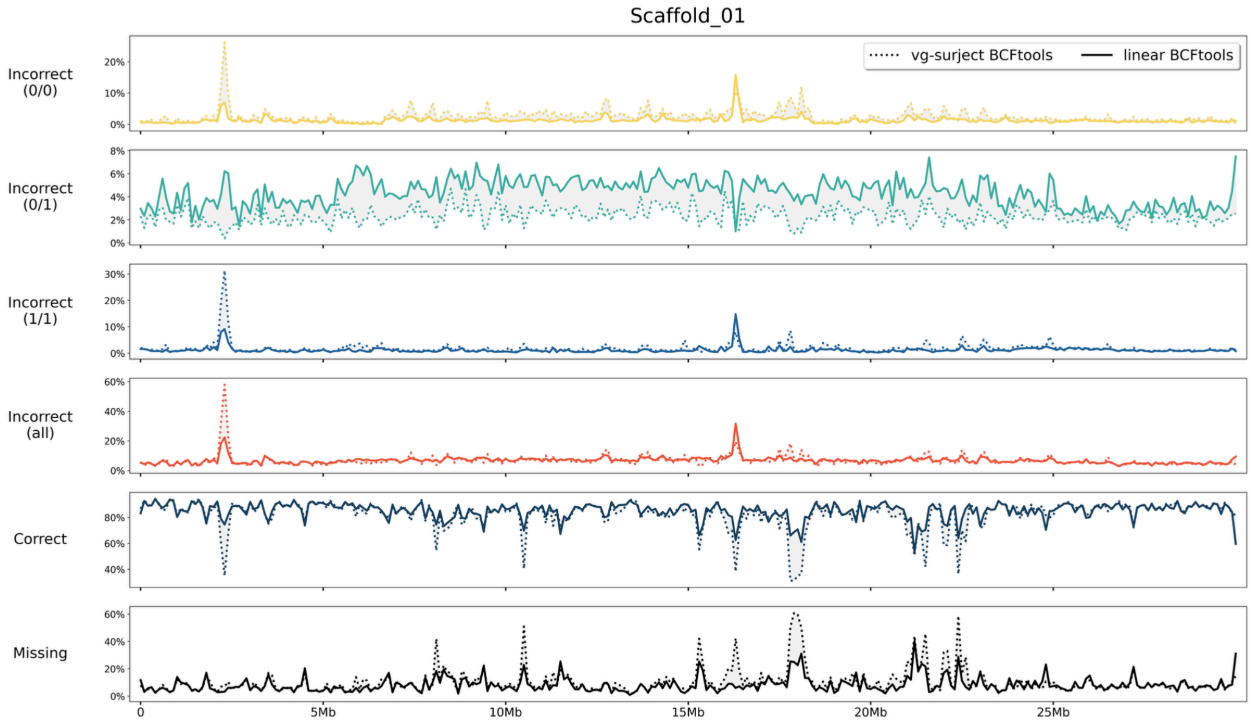


**Figure S12.** Summary comparison of genotype call types from linear-BCFtools (solid line) and vg-surject-BCFtools (dotted line) across 100 kb windows on chromosome 1. Panels from top to bottom: incorrect homozygous reference (yellow), incorrect heterozygous (green), incorrect homozygous alternate (blue), incorrect total (red), correct (dark blue), missing (black).

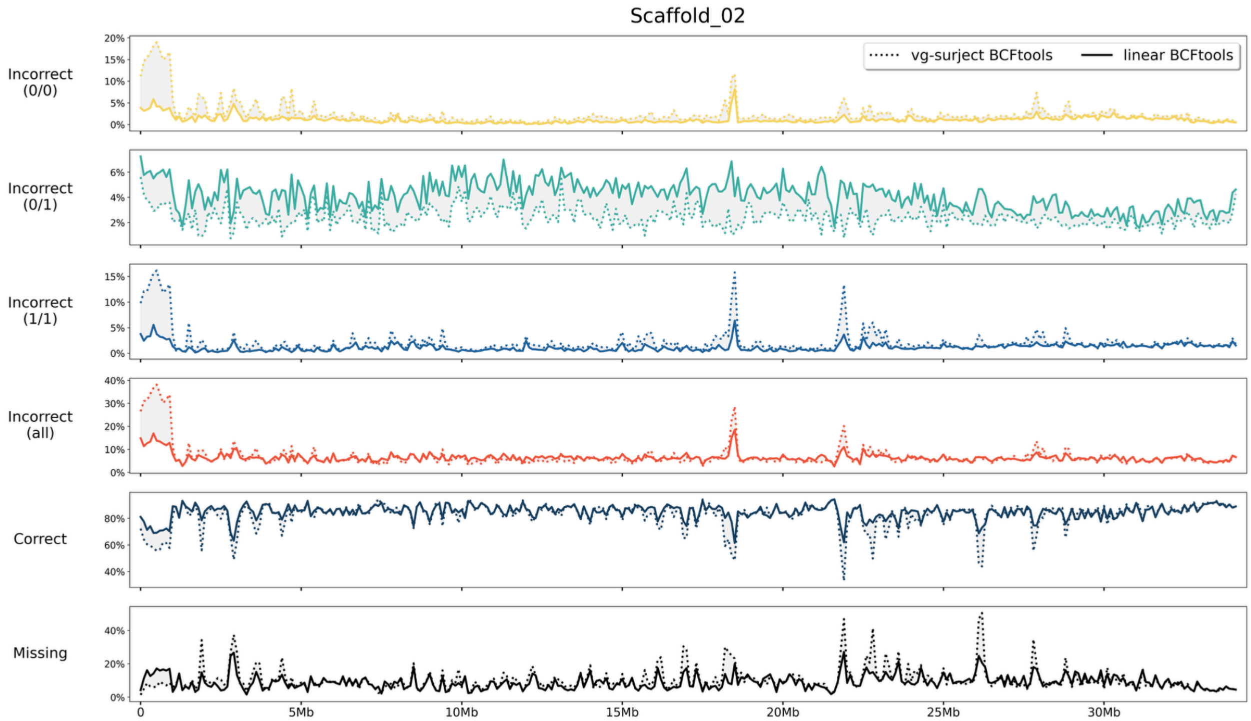


**Figure S13.** Summary comparison of genotype call types from linear-BCFtools (solid line) and vg-surject-BCFtools (dotted line) across 100 kb windows on chromosome 2. Panels from top to bottom: incorrect homozygous reference (yellow), incorrect heterozygous (green), incorrect homozygous alternate (blue), incorrect total (red), correct (dark blue), missing (black).

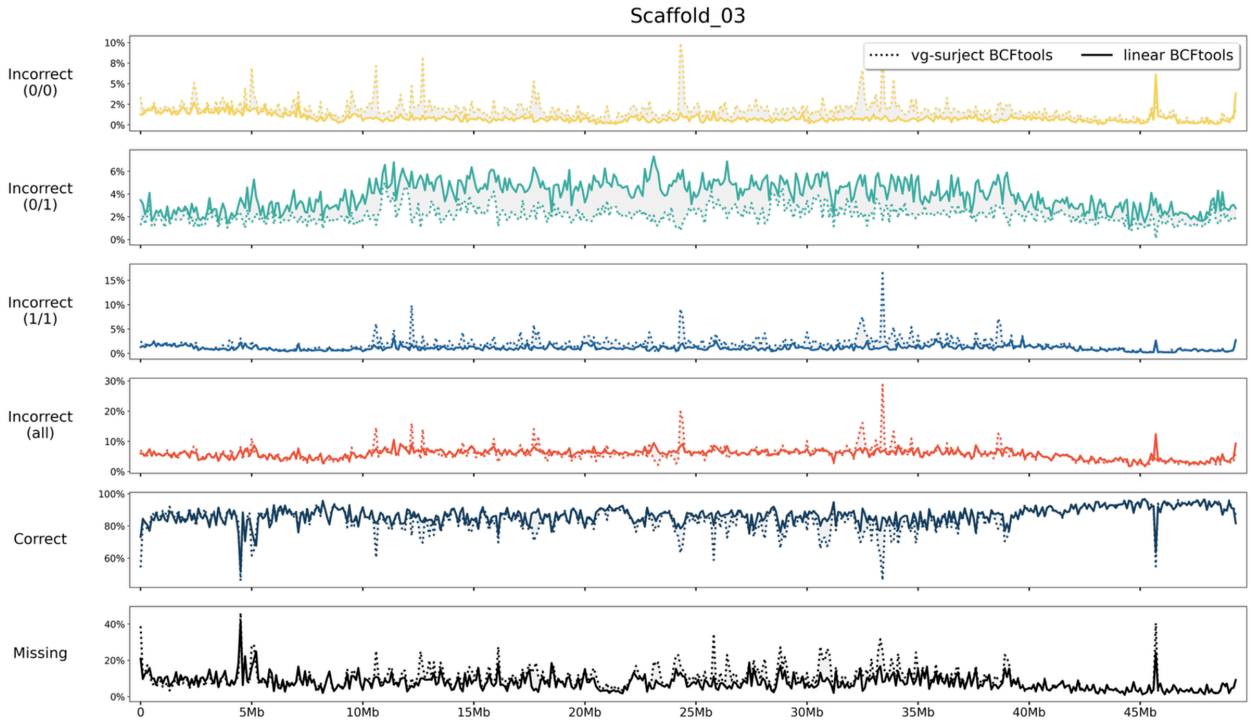


**Figure S14.** Summary comparison of genotype call types from linear-BCFtools (solid line) and vg-surject-BCFtools (dotted line) across 100 kb windows on chromosome 3. Panels from top to bottom: incorrect homozygous reference (yellow), incorrect heterozygous (green), incorrect homozygous alternate (blue), incorrect total (red), correct (dark blue), missing (black).

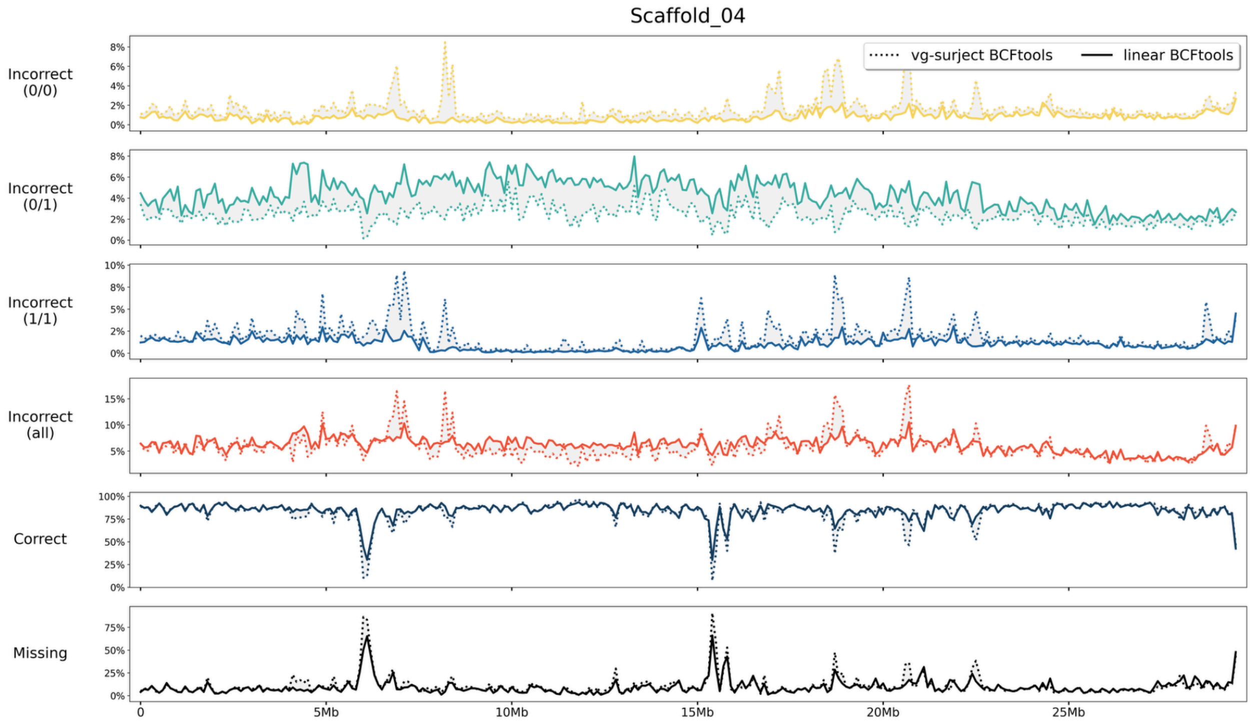


**Figure S15.** Summary comparison of genotype call types from linear-BCFtools (solid line) and vg-surject-BCFtools (dotted line) across 100 kb windows on chromosome 4. Panels from top to bottom: incorrect homozygous reference (yellow), incorrect heterozygous (green), incorrect homozygous alternate (blue), incorrect total (red), correct (dark blue), missing (black).

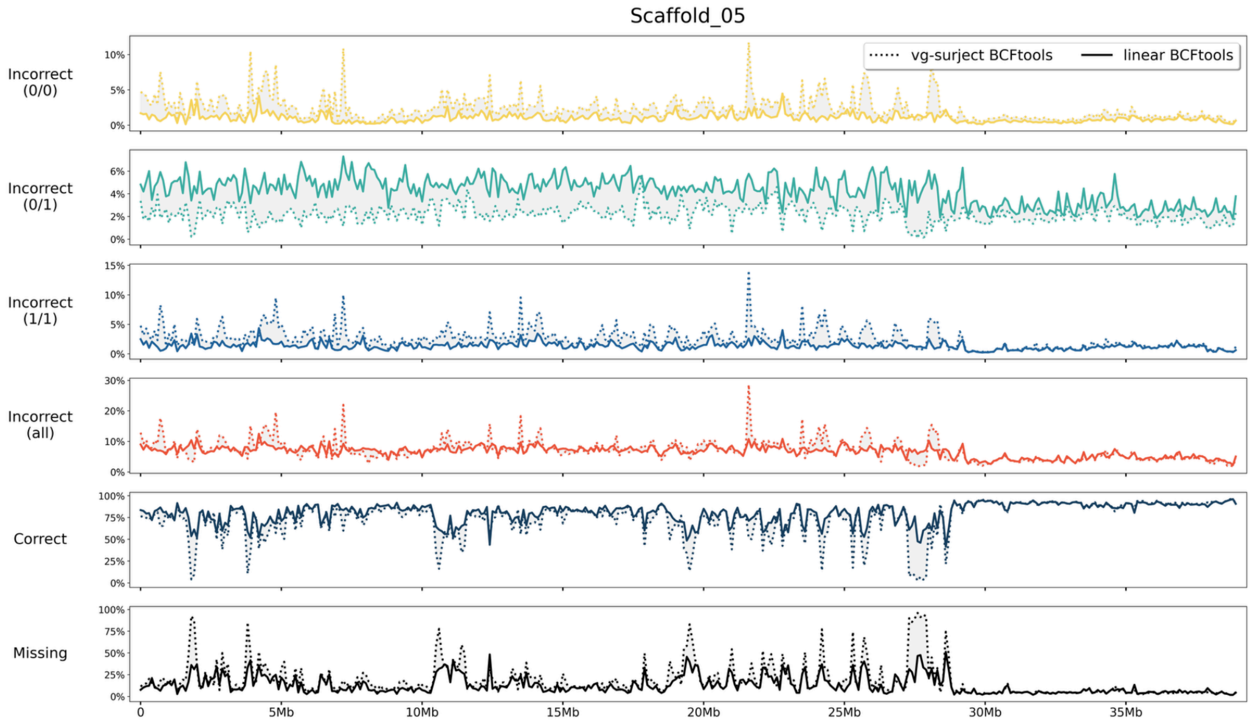


**Figure S16.** Summary comparison of genotype call types from linear-BCFtools (solid line) and vg-surject-BCFtools (dotted line) across 100 kb windows on chromosome 5. Panels from top to bottom: incorrect homozygous reference (yellow), incorrect heterozygous (green), incorrect homozygous alternate (blue), incorrect total (red), correct (dark blue), missing (black).

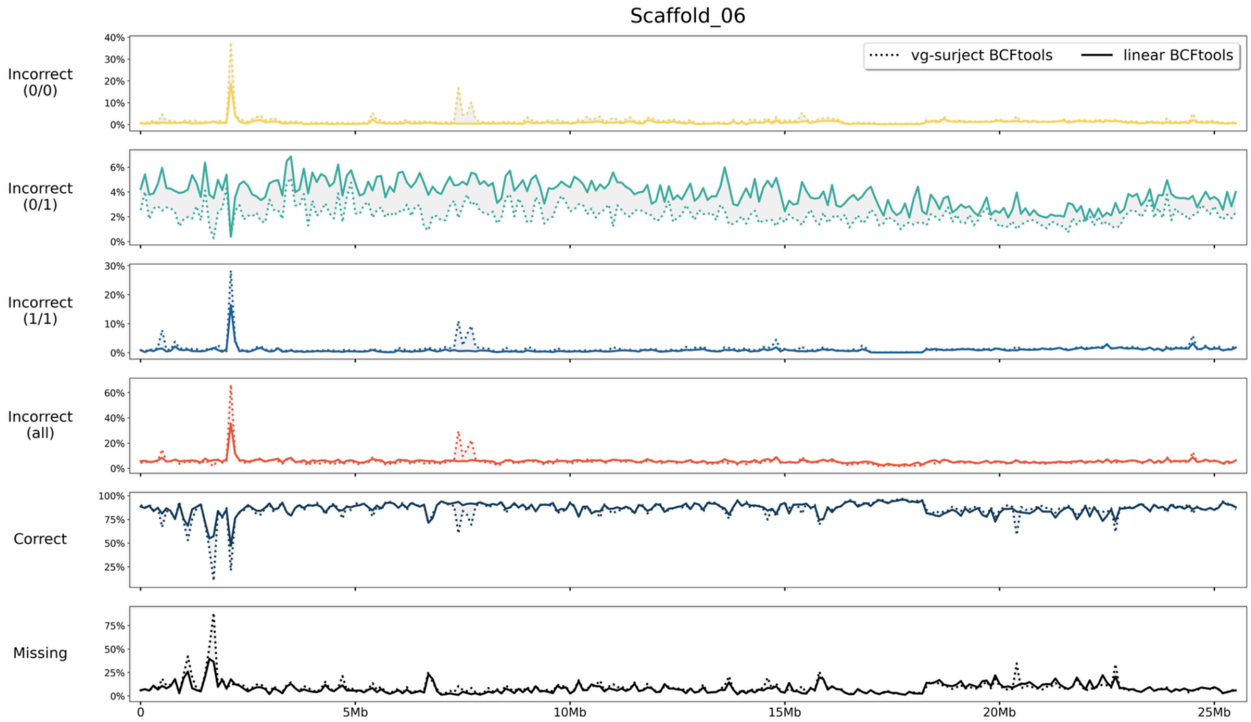


**Figure S17.** Summary comparison of genotype call types from linear-BCFtools (solid line) and vg-surject-BCFtools (dotted line) across 100 kb windows on chromosome 6. Panels from top to bottom: incorrect homozygous reference (yellow), incorrect heterozygous (green), incorrect homozygous alternate (blue), incorrect total (red), correct (dark blue), missing (black).

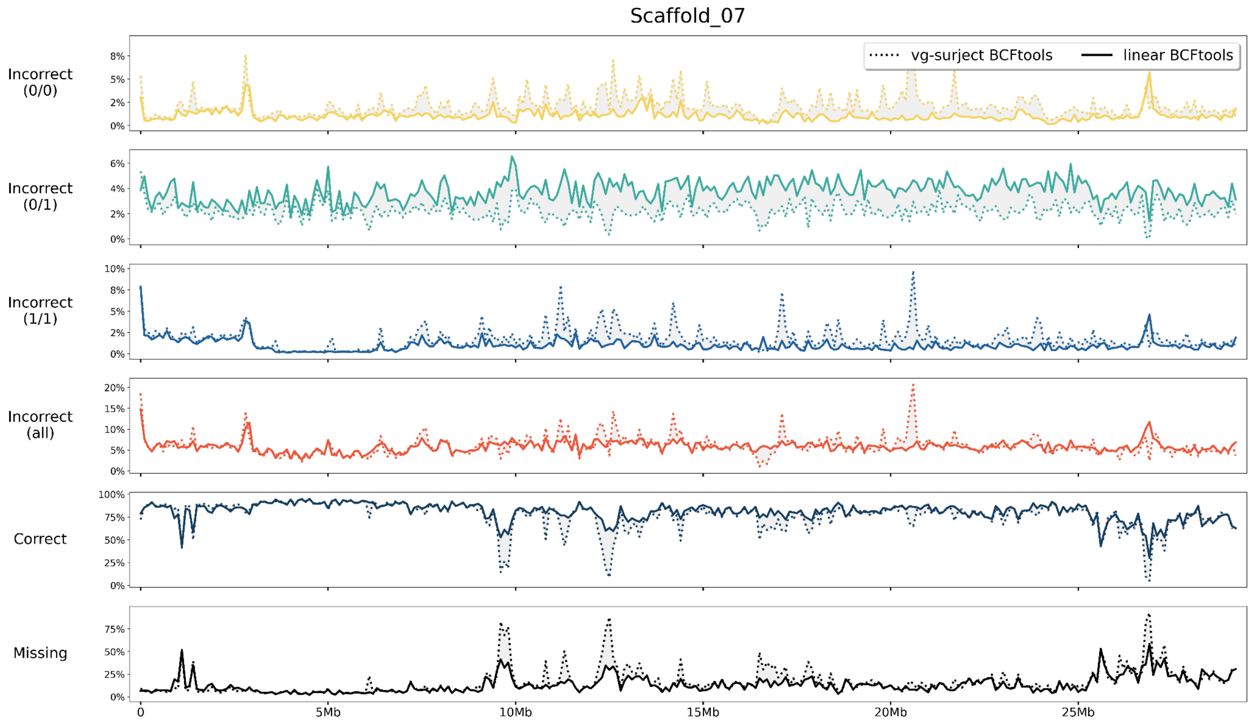


**Figure S18.** Summary comparison of genotype call types from linear-BCFtools (solid line) and vg-surject-BCFtools (dotted line) across 100 kb windows on chromosome 7. Panels from top to bottom: incorrect homozygous reference (yellow), incorrect heterozygous (green), incorrect homozygous alternate (blue), incorrect total (red), correct (dark blue), missing (black).

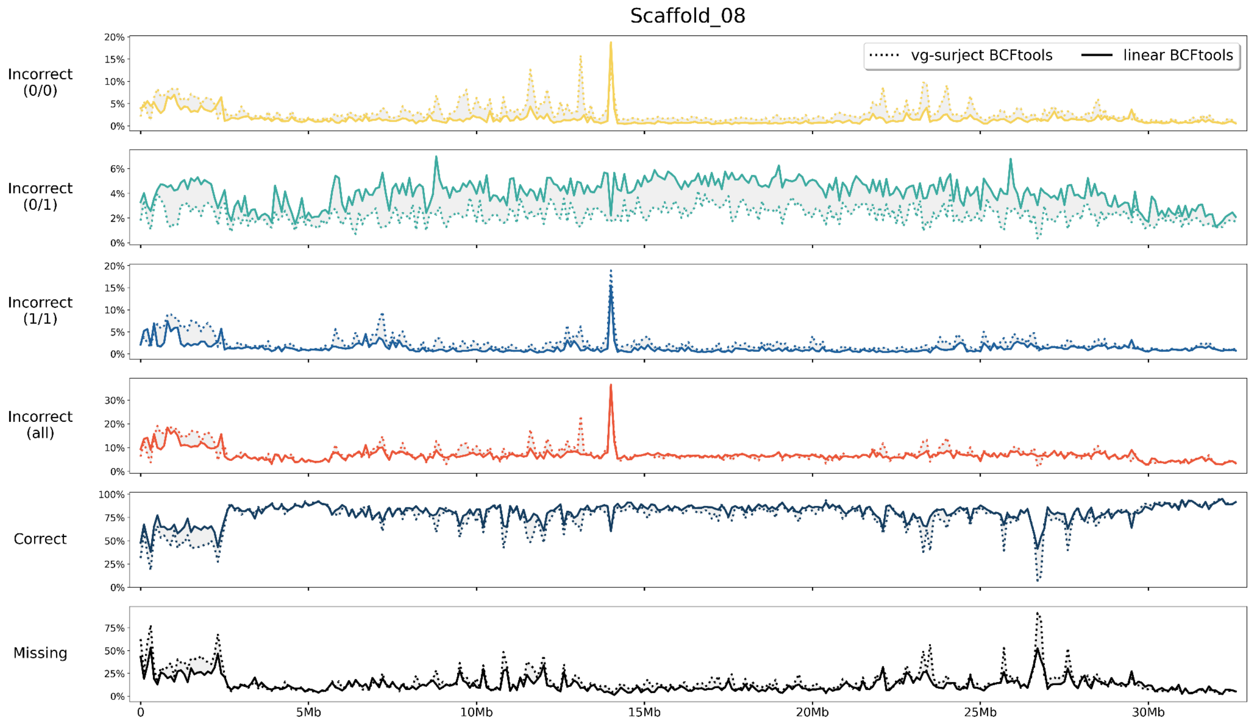


**Figure S19.** Summary comparison of genotype call types from linear-BCFtools (solid line) and vg-surject-BCFtools (dotted line) across 100 kb windows on chromosome 8. Panels from top to bottom: incorrect homozygous reference (yellow), incorrect heterozygous (green), incorrect homozygous alternate (blue), incorrect total (red), correct (dark blue), missing (black).

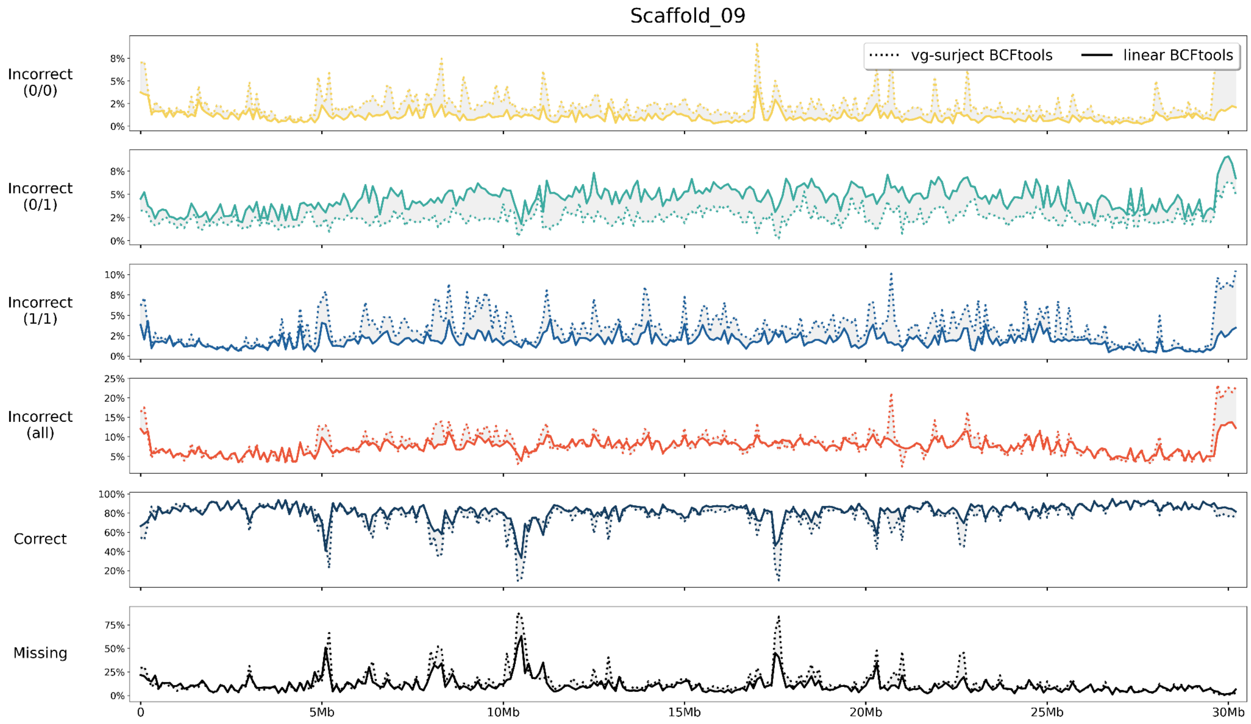


**Figure S20.** Summary comparison of genotype call types from linear-BCFtools (solid line) and vg-surject-BCFtools (dotted line) across 100 kb windows on chromosome 9. Panels from top to bottom: incorrect homozygous reference (yellow), incorrect heterozygous (green), incorrect homozygous alternate (blue), incorrect total (red), correct (dark blue), missing (black).

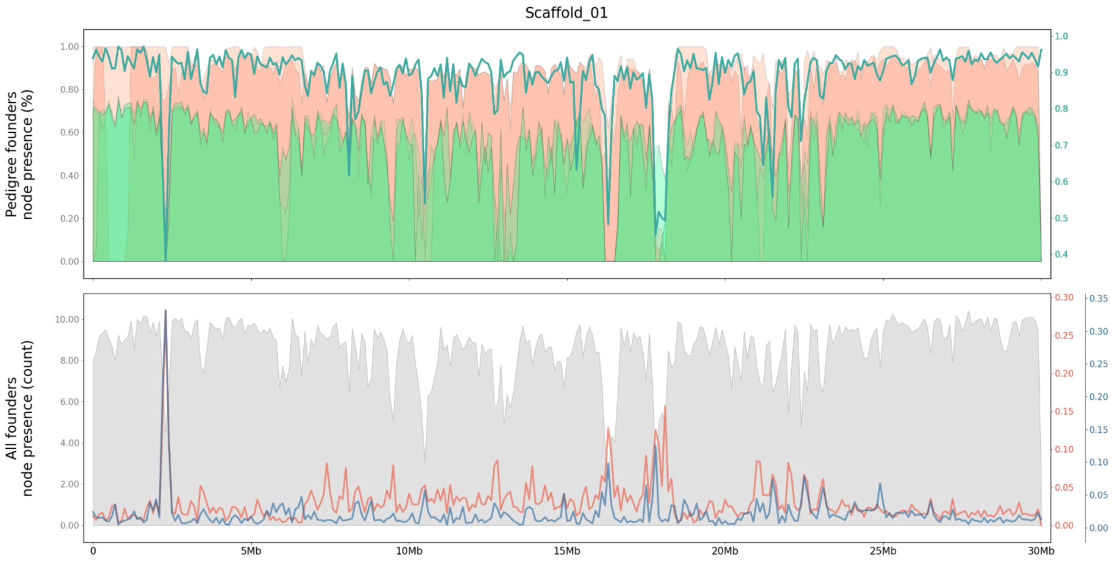


**Figure S21**. Relationship of founder presence in the graph nodes per 100 kb windows along chromosome 1. **(Top)** The node presence of pedigree founders (orange = mandarin; green = Australian lime) vs. the frequency of genotypes per window that are correct (green line). **(Bottom)** The number of founders present per-node vs. the rates of incorrect homozygous reference calls (red), and incorrect homozygous alternate calls (blue).

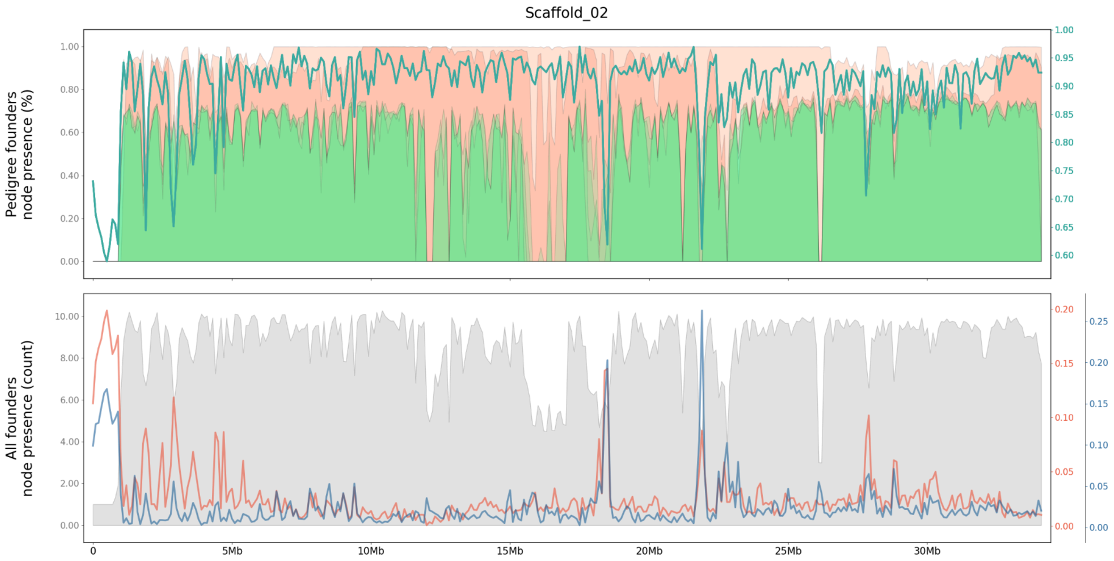


**Figure S22**. Relationship of founder presence in the graph nodes per 100 kb windows along chromosome 2. **(Top)** The node presence of pedigree founders (orange = mandarin; green = Australian lime) vs. the frequency of genotypes per window that are correct (green line). **(Bottom)** The number of founders present per-node vs. the rates of incorrect homozygous reference calls (red), and incorrect homozygous alternate calls (blue).

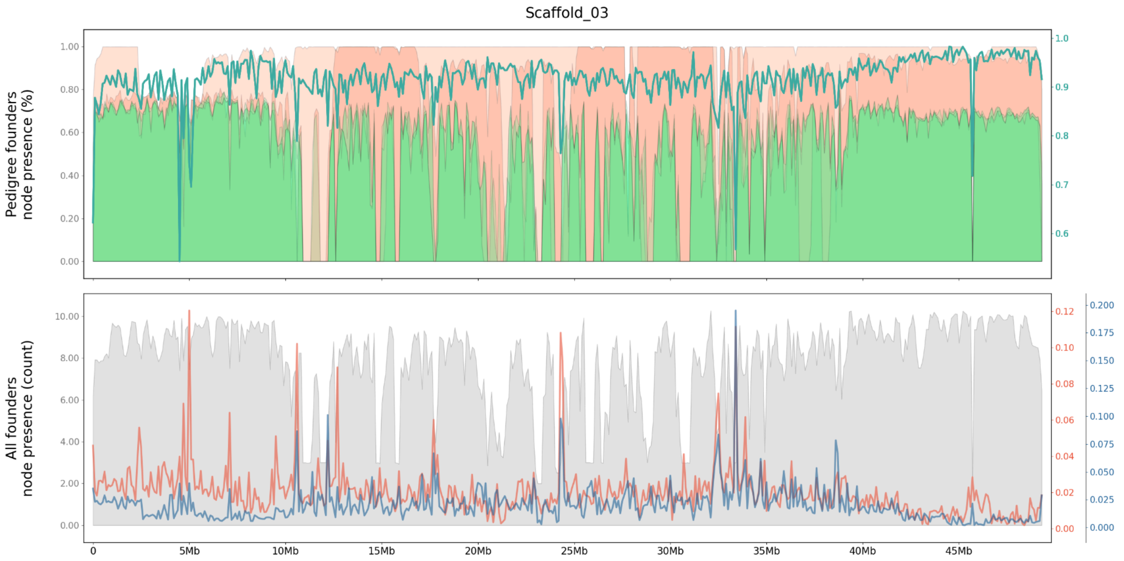


**Figure S23**. Relationship of founder presence in the graph nodes per 100 kb windows along chromosome 3. **(Top)** The node presence of pedigree founders (orange = mandarin; green = Australian lime) vs. the frequency of genotypes per window that are correct (green line). **(Bottom)** The number of founders present per-node vs. the rates of incorrect homozygous reference calls (red), and incorrect homozygous alternate calls (blue).

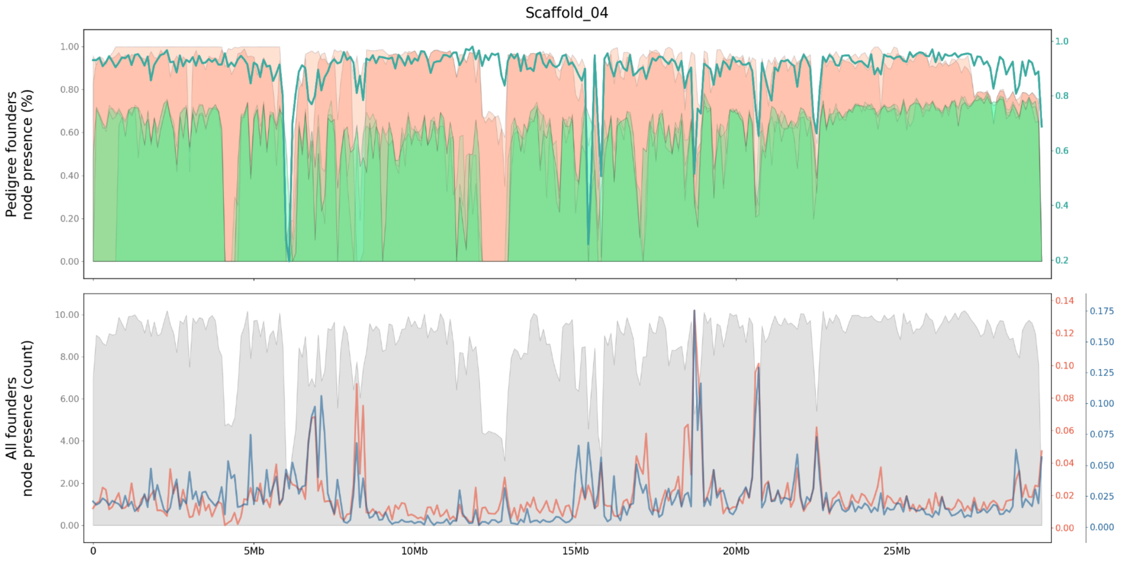


**Figure S24**. Relationship of founder presence in the graph nodes per 100 kb windows along chromosome 4. **(Top)** The node presence of pedigree founders (orange = mandarin; green = Australian lime) vs. the frequency of genotypes per window that are correct (green line). **(Bottom)** The number of founders present per-node vs. the rates of incorrect homozygous reference calls (red), and incorrect homozygous alternate calls (blue).

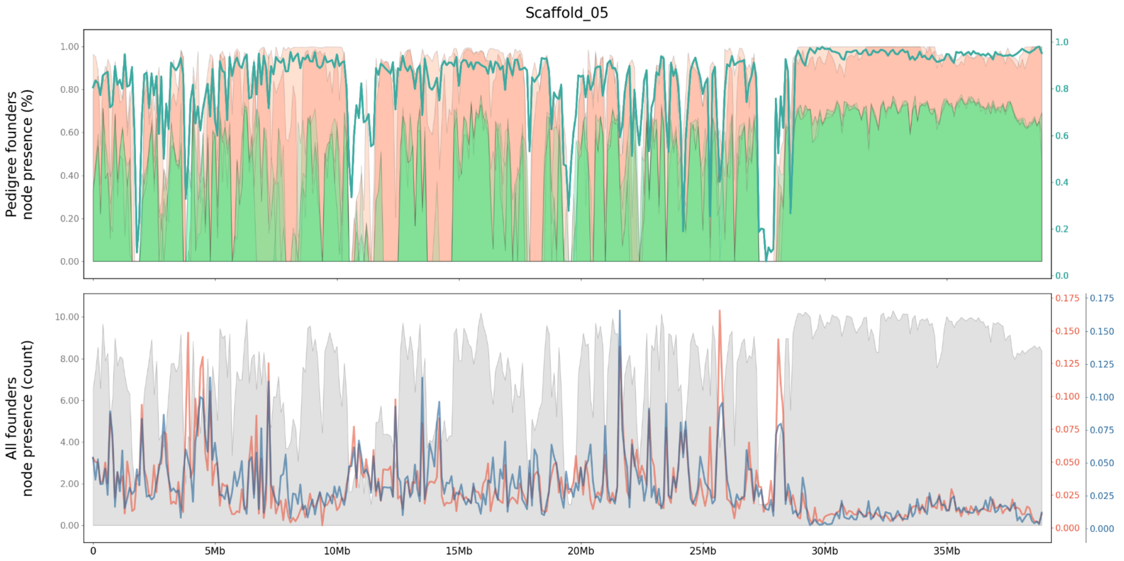


**Figure S25**. Relationship of founder presence in the graph nodes per 100 kb windows along chromosome 5. **(Top)** The node presence of pedigree founders (orange = mandarin; green = Australian lime) vs. the frequency of genotypes per window that are correct (green line). **(Bottom)** The number of founders present per-node vs. the rates of incorrect homozygous reference calls (red), and incorrect homozygous alternate calls (blue).

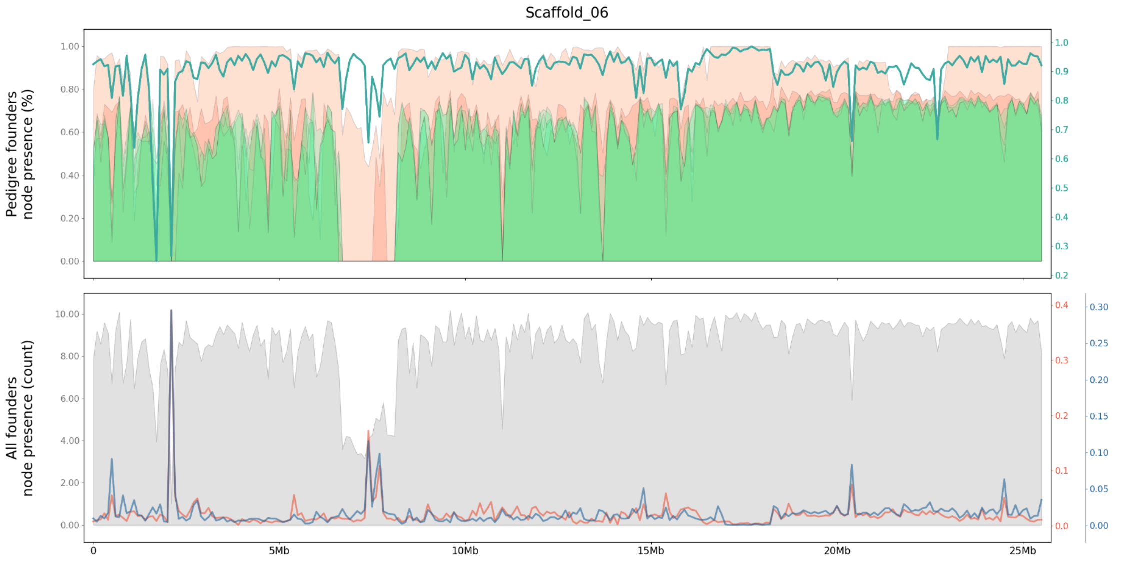


**Figure S26**. Relationship of founder presence in the graph nodes per 100 kb windows along chromosome 6. **(Top)** The node presence of pedigree founders (orange = mandarin; green = Australian lime) vs. the frequency of genotypes per window that are correct (green line). **(Bottom)** The number of founders present per-node vs. the rates of incorrect homozygous reference calls (red), and incorrect homozygous alternate calls (blue).

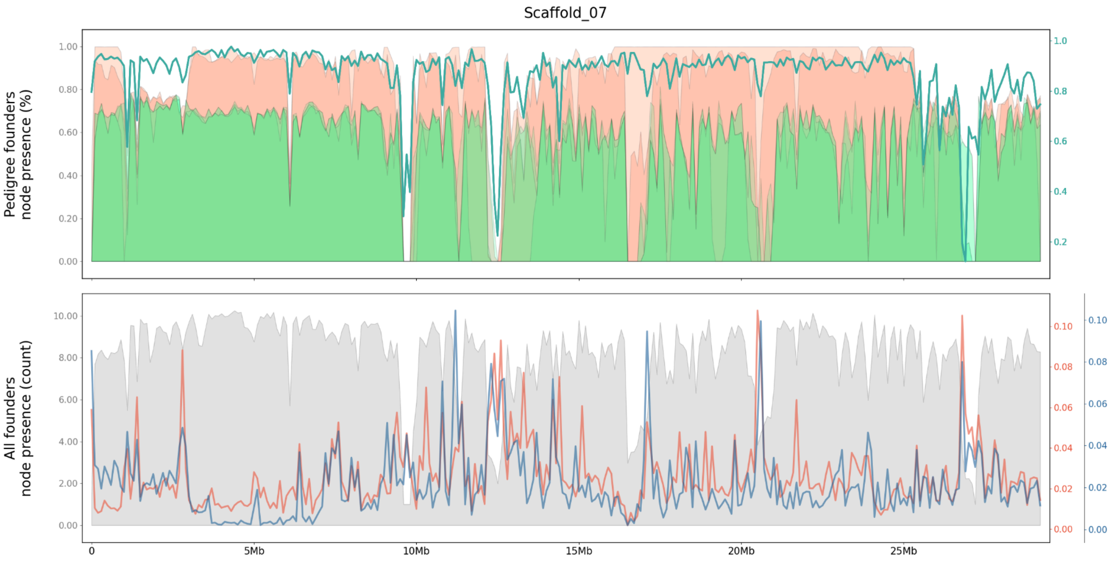


**Figure S27**. Relationship of founder presence in the graph nodes per 100 kb windows along chromosome 7. **(Top)** The node presence of pedigree founders (orange = mandarin; green = Australian lime) vs. the frequency of genotypes per window that are correct (green line). **(Bottom)** The number of founders present per-node vs. the rates of incorrect homozygous reference calls (red), and incorrect homozygous alternate calls (blue).

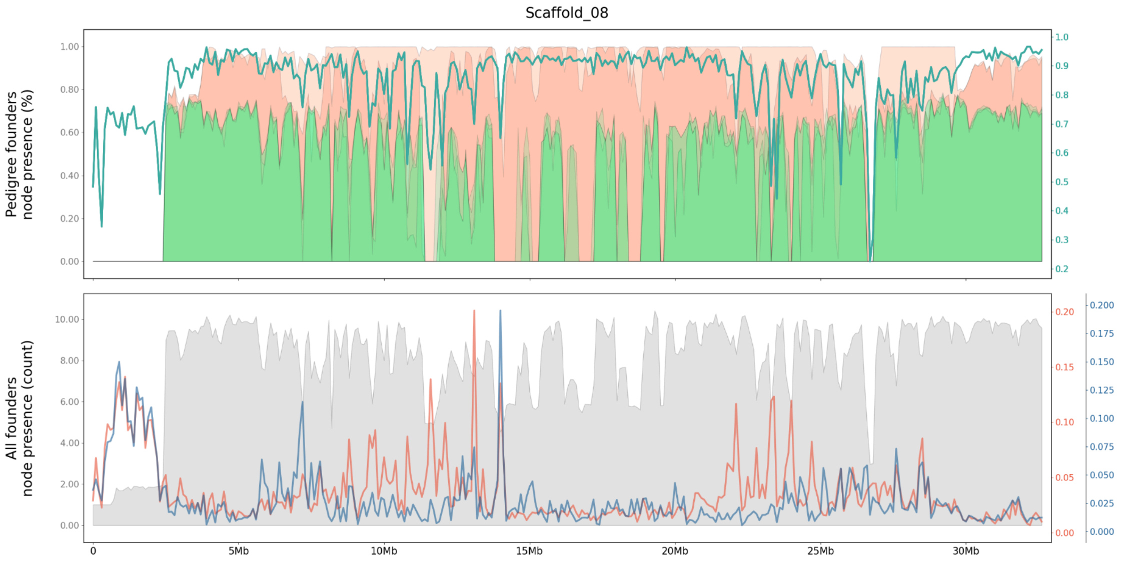


**Figure S28**. Relationship of founder presence in the graph nodes per 100 kb windows along chromosome 8. **(Top)** The node presence of pedigree founders (orange = mandarin; green = Australian lime) vs. the frequency of genotypes per window that are correct (green line). **(Bottom)** The number of founders present per-node vs. the rates of incorrect homozygous reference calls (red), and incorrect homozygous alternate calls (blue).

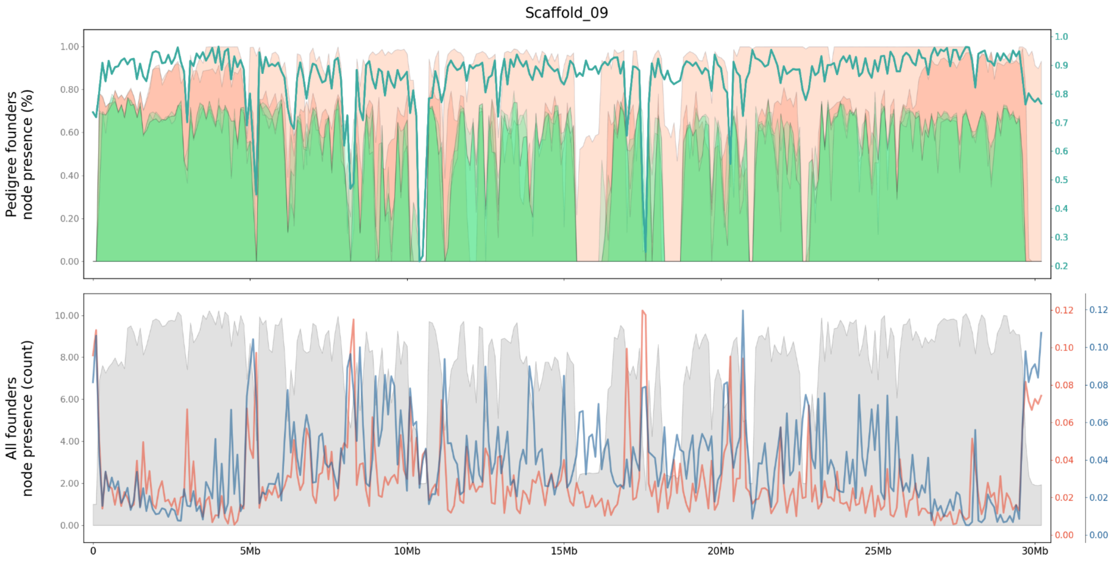


**Figure S29**. Relationship of founder presence in the graph nodes per 100 kb windows along chromosome 9. **(Top)** The node presence of pedigree founders (orange = mandarin; green = Australian lime) vs. the frequency of genotypes per window that are correct (green line). **(Bottom)** The number of founders present per-node vs. the rates of incorrect homozygous reference calls (red), and incorrect homozygous alternate calls (blue).

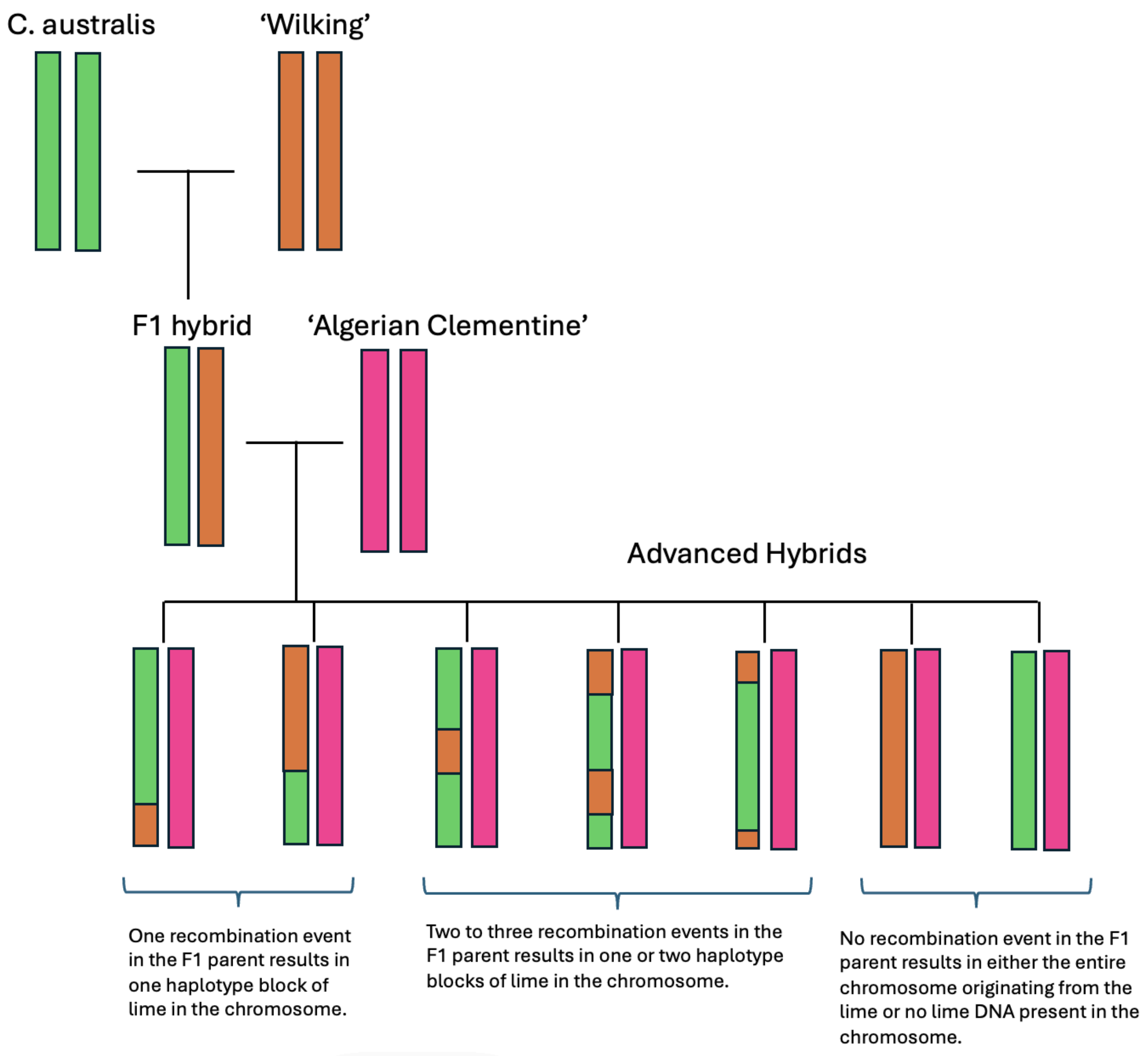


**Figure S30.** Representation of a single chromosome in the F1 and advanced hybrid breeding program.

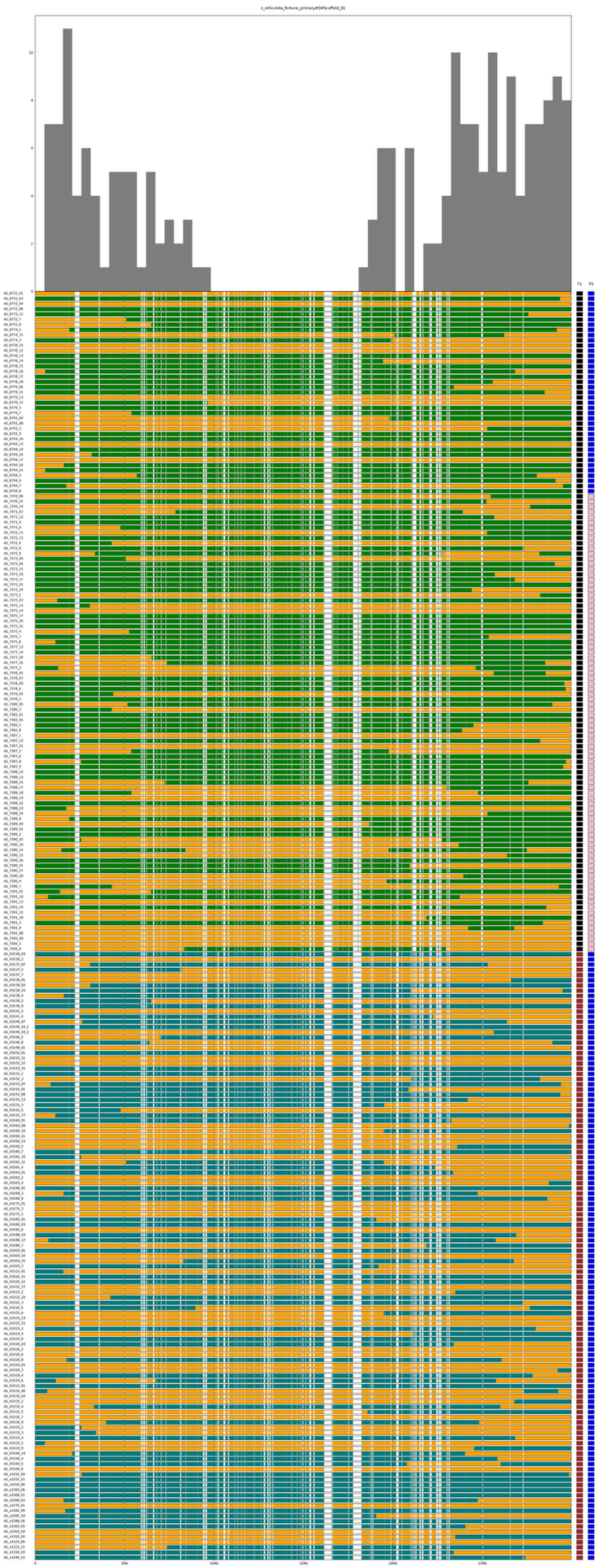
c_reticulata_fortune_primary#0#Scaffold_01
F1
F2
0
5Mb
10Mb
15Mb
20Mb
25Mb

**Figure S31.** Recombination frequencies on chromosome 1 (top bar chart) derived from parental haplotypes reconstruction in 244 advanced hybrids (bottom rows). Each of the three hybrid families studied are denoted with the F1 parent (black = 475_01; brown = 246_01) and P3 outcrossed parent (blue = 'Algerian Clementine'; pink = 'Kiyomi Tangor'). Orange blocks indicate the presence of a mandarin grandparent source and the Australian lime contribution is indicated by either green (*C. australasica*) or dark teal (*C. australis*).

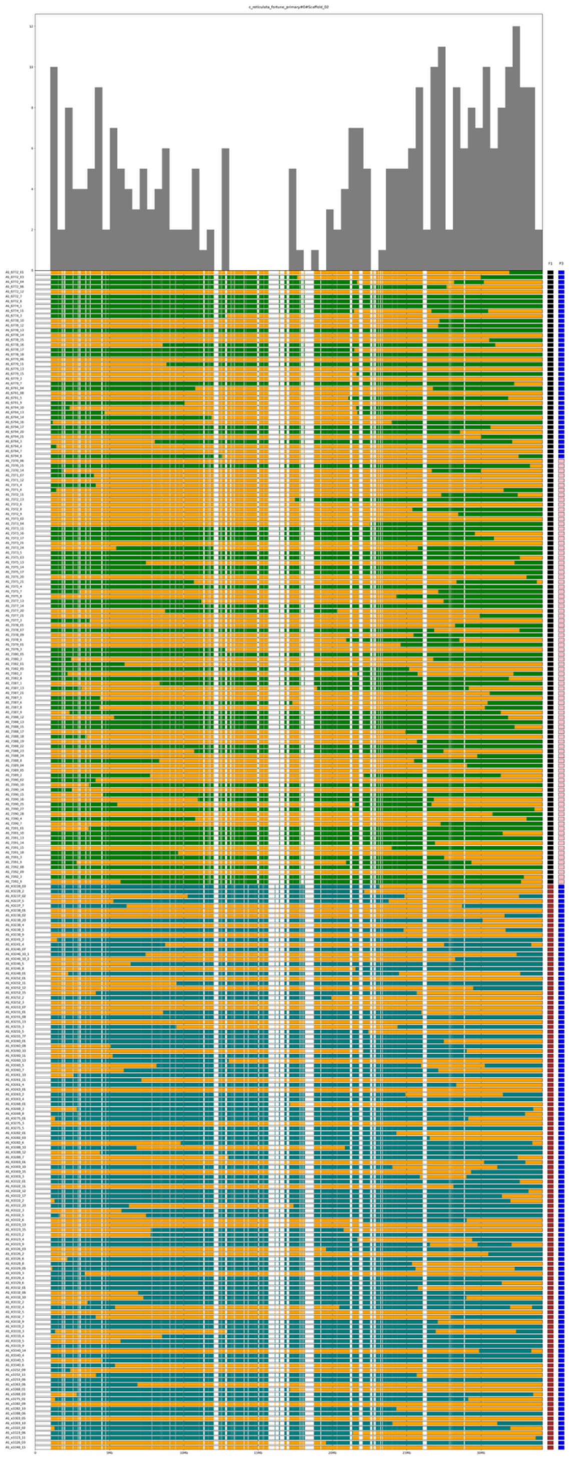

**Figure S32.** Recombination frequencies on chromosome 2 (top bar chart) derived from parental haplotypes reconstruction in 244 advanced hybrids (bottom rows). Each of the three hybrid families studied are denoted with the F1 parent (black = 475_01; brown = 246_01) and P3 outcrossed parent (blue = 'Algerian Clementine'; pink = 'Kiyomi Tangor'). Orange blocks indicate the presence of a mandarin grandparent source and the Australian lime contribution is indicated by either green (*C. australasica*) or dark teal (*C. australis*).

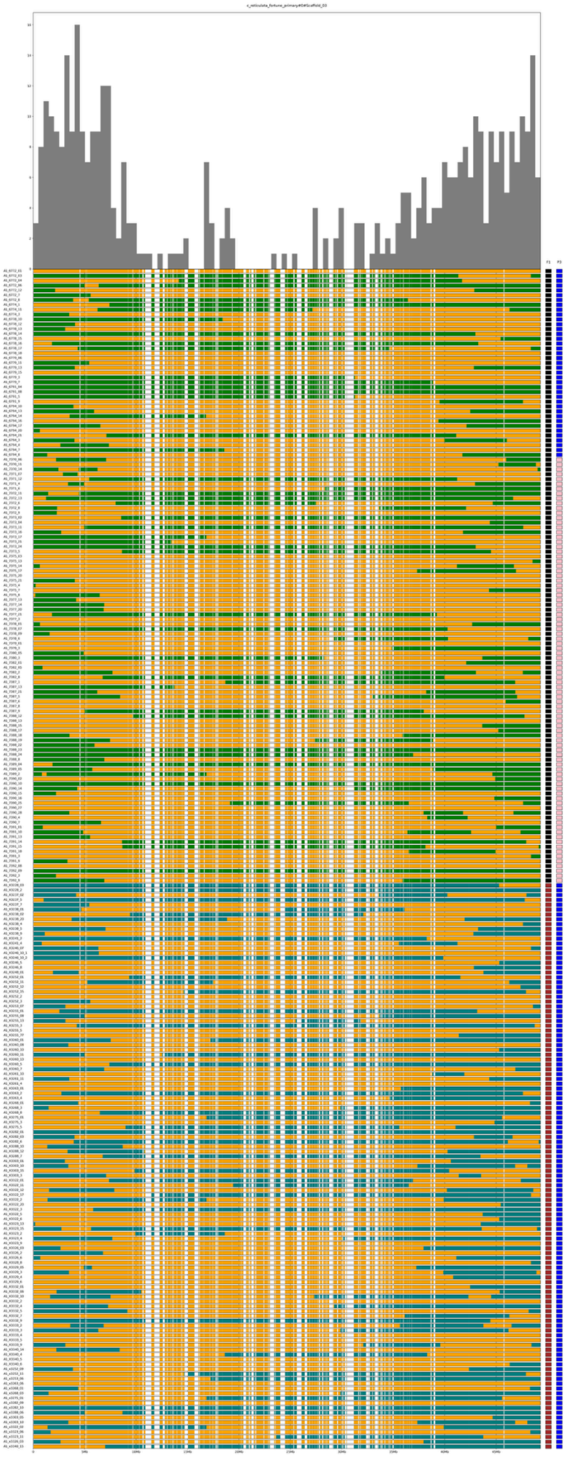

**Figure S33.** Recombination frequencies on chromosome 3 (top bar chart) derived from parental haplotypes reconstruction in 244 advanced hybrids (bottom rows). Each of the three hybrid families studied are denoted with the F1 parent (black = 475_01; brown = 246_01) and P3 outcrossed parent (blue = ‘Algerian Clementine’; pink = ‘Kiyomi Tangor’). Orange blocks indicate the presence of a mandarin grandparent source and the Australian lime contribution is indicated by either green (*C. australasica*) or dark teal (*C. australis*).

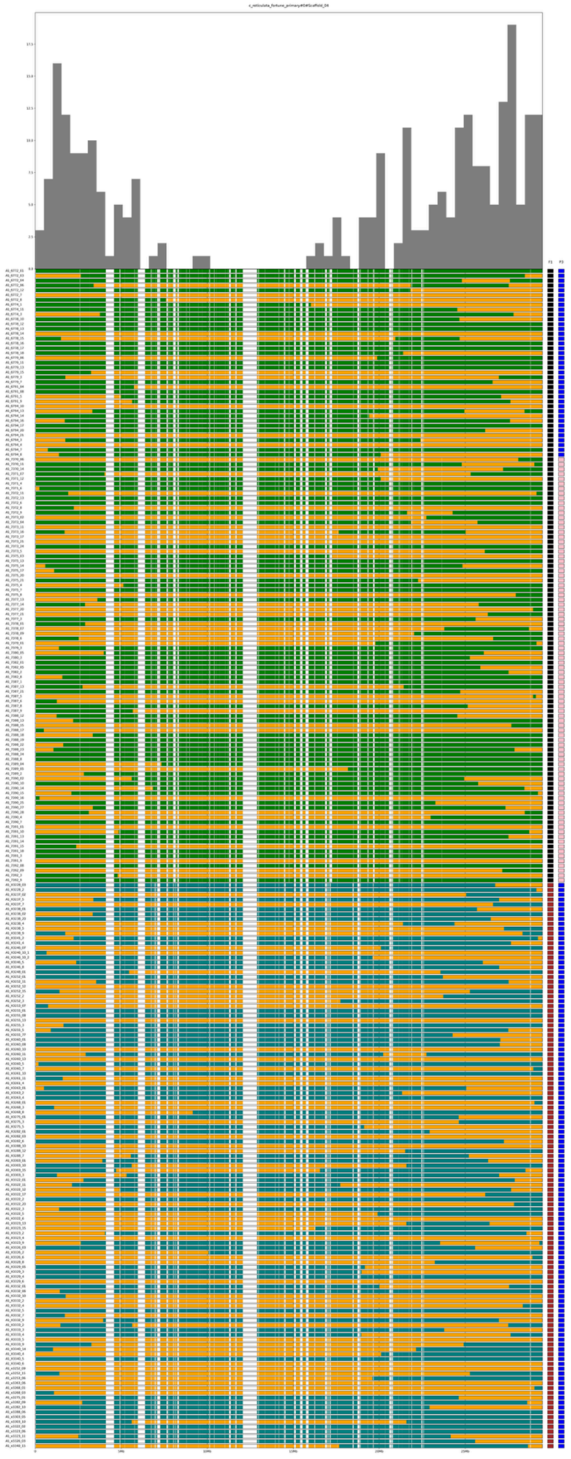

**Figure S34.** Recombination frequencies on chromosome 4 (top bar chart) derived from parental haplotypes reconstruction in 244 advanced hybrids (bottom rows). Each of the three hybrid families studied are denoted with the F1 parent (black = 475_01; brown = 246_01) and P3 outcrossed parent (blue = 'Algerian Clementine'; pink = 'Kiyomi Tangor'). Orange blocks indicate the presence of a mandarin grandparent source and the Australian lime contribution is indicated by either green (*C. australasica*) or dark teal (*C. australis*).

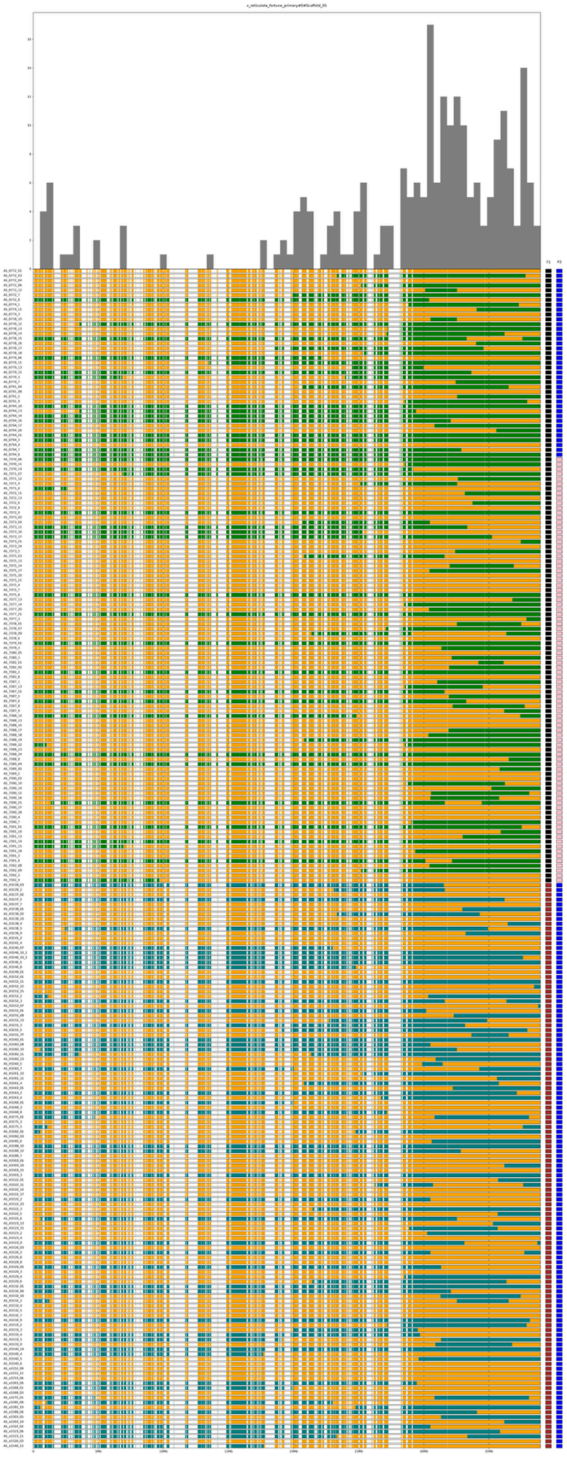

**Figure S35.** Recombination frequencies on chromosome 5 (top bar chart) derived from parental haplotypes reconstruction in 244 advanced hybrids (bottom rows). Each of the three hybrid families studied are denoted with the F1 parent (black = 475_01; brown = 246_01) and P3 outcrossed parent (blue = 'Algerian Clementine'; pink = 'Kiyomi Tangor'). Orange blocks indicate the presence of a mandarin grandparent source and the Australian lime contribution is indicated by either green (*C. australasica*) or dark teal (*C. australis*).

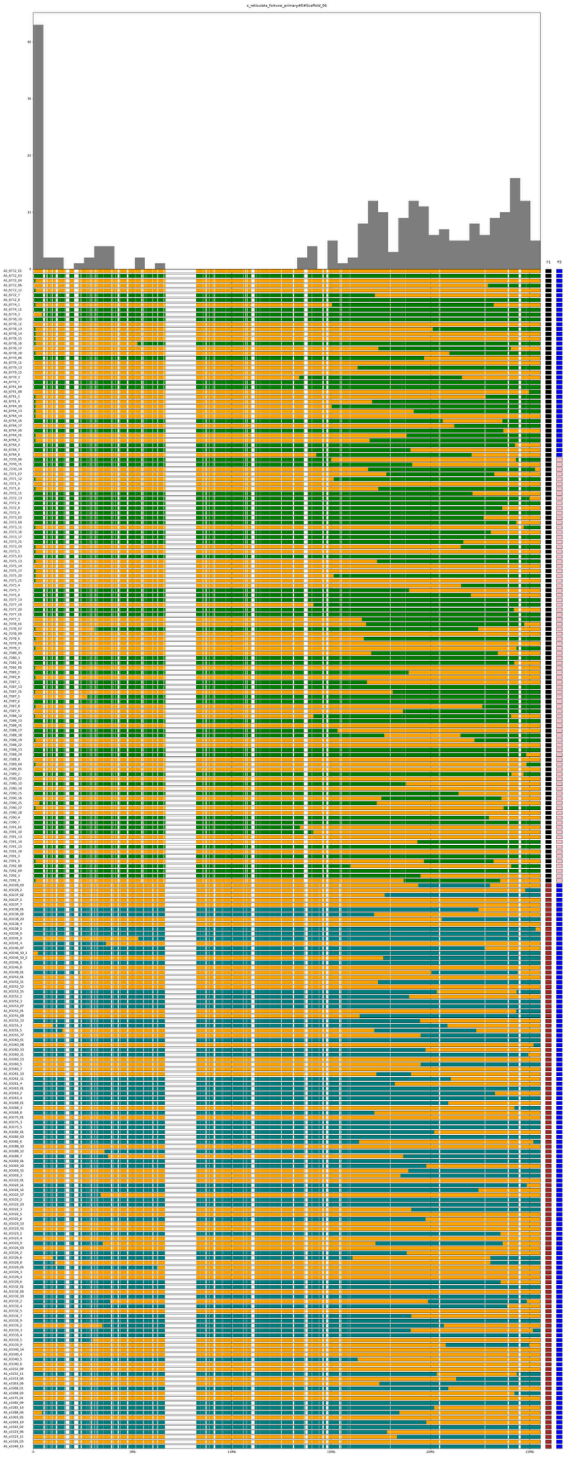

**Figure S36.** Recombination frequencies on chromosome 6 (top bar chart) derived from parental haplotypes reconstruction in 244 advanced hybrids (bottom rows). Each of the three hybrid families studied are denoted with the F1 parent (black = 475_01; brown = 246_01) and P3 outcrossed parent (blue = ‘Algerian Clementine’; pink = ‘Kiyomi Tangor’). Orange blocks indicate the presence of a mandarin grandparent source and the Australian lime contribution is indicated by either green (*C. australasica*) or dark teal (*C. australis*).

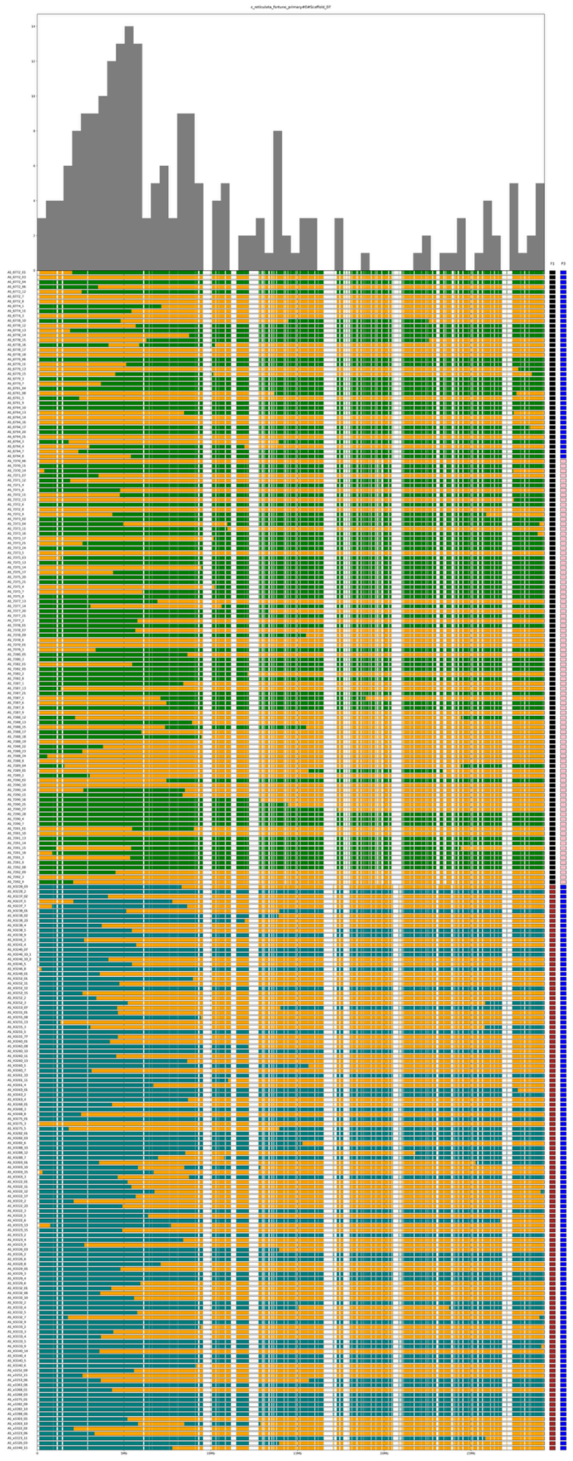

**Figure S37.** Recombination frequencies on chromosome 7 (top bar chart) derived from parental haplotypes reconstruction in 244 advanced hybrids (bottom rows). Each of the three hybrid families studied are denoted with the F1 parent (black = 475_01; brown = 246_01) and P3 outcrossed parent (blue = 'Algerian Clementine'; pink = 'Kiyomi Tangor'). Orange blocks indicate the presence of a mandarin grandparent source and the Australian lime contribution is indicated by either green (*C. australasica*) or dark teal (*C. australis*).

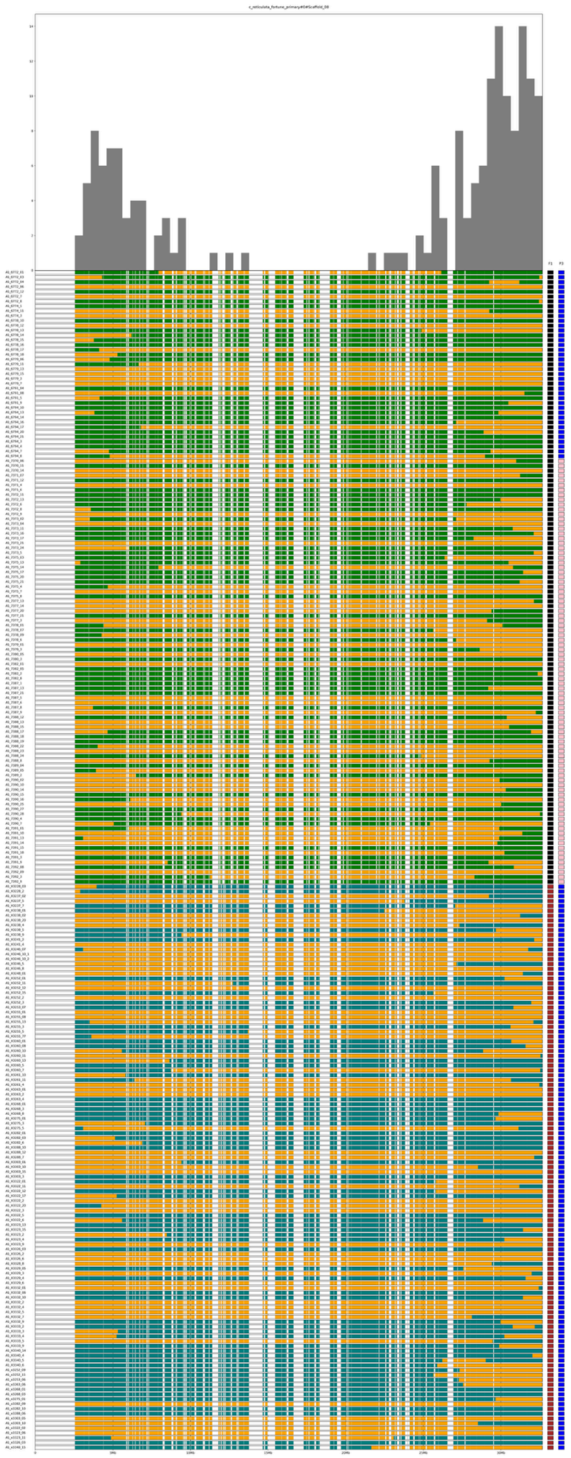

**Figure S38.** Recombination frequencies on chromosome 8 (top bar chart) derived from parental haplotypes reconstruction in 244 advanced hybrids (bottom rows). Each of the three hybrid families studied are denoted with the F1 parent (black = 475_01; brown = 246_01) and P3 outcrossed parent (blue = ‘Algerian Clementine’; pink = ‘Kiyomi Tangor’). Orange blocks indicate the presence of a mandarin grandparent source and the Australian lime contribution is indicated by either green (*C. australasica*) or dark teal (*C. australis*).

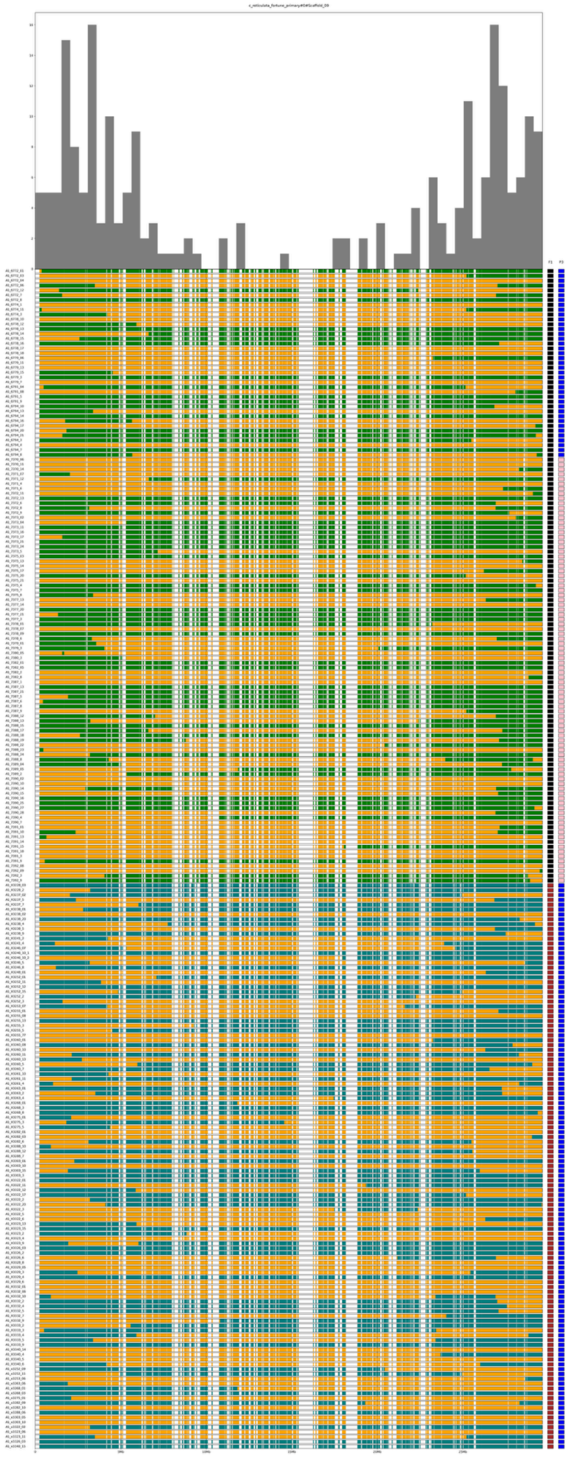

**Figure S39.** Recombination frequencies on chromosome 9 (top bar chart) derived from parental haplotypes reconstruction in 244 advanced hybrids (bottom rows). Each of the three hybrid families studied are denoted with the F1 parent (black = 475_01; brown = 246_01) and P3 outcrossed parent (blue = 'Algerian Clementine'; pink = 'Kiyomi Tangor'). Orange blocks indicate the presence of a mandarin grandparent source and the Australian lime contribution is indicated by either green (*C. australasica*) or dark teal (*C. australis*).
