## SupplementalTables for "Benchmarking SNP-Calling Accuracy Against Known *Citrus* Pedigrees Reveals Pangenome Advantages Over Linear References"

**Table S1.** Summary of haplotype resolved chromosome scaled genome assemblies statistics.

| Accession | Haplotype | Total Mb | Total Scaffolds | Scaffold N50 (Mb) | Mb in 9 Main Chromosomes | # Annotated Genes | Public Location (Citrus Genome Database) | NCBI Accession | Reference |
|---|---|---|---|---|---|---|---|---|---|
| *C. australasica* | Primary | 336.72 | 270 | 29.93 | 290.01 | 23,434 | CGD | PRJNA924094 | Singh et al. 2024 |
| | Alternate | 335.26 | 270 | 30.25 | 289.02 | 22,427 | CGD | PRJNA924095 | Singh et al. 2024 |
| *C. australis* | Primary | 321.53 | 127 | 30.33 | 291.56 | 24,602 | CGD26003 | PRJNA1418802 | Singh et al, unpublished data |
| | Alternate | 321.53 | 137 | 30.33 | 289.99 | 25,433 | CGD26003 | PRJNA1418803 | Singh et al, unpublished data |
| *C. inodora* | Primary | 303.76 | 299 | 28.92 | 276.93 | 28,176 | CGD | PRJNA924048 | Singh et al. 2024 |
| | Alternate | 298.85 | 299 | 28.92 | 272.04 | 27,665 | CGD | PRJNA924114 | Singh et al. 2024 |
| 'Fortune' | Primary | 321.79 | 170 | 30.27 | 299.81 | 24,939 | CGD26002 | PRJN1418788 | Singh et al, unpublished data |
| | Alternate | 320.26 | 144 | 30.33 | 293.01 | 24,107 | CGD26002 | PRJN1418787 | Singh et al, unpublished data |
| 'Fallglo' | Primary | 325.06 | 162 | 30.76 | 300.99 | 24,131 | CGD26004 | PRJNA1418798 | Singh et al, unpublished data |
| | Alternate | 322.39 | 166 | 30.43 | 295.33 | 24,876 | CGD26004 | PRJNA1418799 | Singh et al, unpublished data |
| 'Wilking' | Primary | 337.91 | 101 | 29.8 | 303.77 | 25,609 | CGD26001 | PRJNA1418772 | Singh et al, unpublished data |
| | Alternate | 337.95 | 106 | 30.16 | 303.76 | 25,566 | CGD26001 | PRJNA1418771 | Singh et al, unpublished data |

**Table S2.** F1 and advanced hybrid Illumina data. Coverage is based off of the length of the 'Fortune' primary accession.

| Sample Name | # of Reads | Coverage | Seed Parent | Pollen Parent | NCBI SRA Accession | BioProject number |
|---|---|---|---|---|---|---|
| **F1 Generation (n=30)** | | | | | | |
| F1_41_02 | 40,997,943 | 31.11 | 'Fortune mandarin' | *C. australis* | SAMN57102146 | PRJNA1448193 |
| F1_41_03 | 35,245,977 | 37.64 | 'Fortune mandarin' | *C. australis* | SAMN57102147 | |
| F1_191_03 | 38,118,017 | 40.70 | 'Wilking mandarin' | *C. inodora* | SAMN57102149 | |
| F1_196_01 | 36,567,126 | 39.05 | 'Wilking mandarin' | *C. inodora* | SAMN57102150 | |
| F1_199_01 | 33,901,691 | 36.20 | 'Wilking mandarin' | *C. inodora* | SAMN57102151 | |
| F1_199_02_90 | 43,356,344 | 46.30 | 'Wilking mandarin' | *C. inodora* | SAMN57102152 | |
| F1_242_01 | 40,642,406 | 43.40 | 'Wilking mandarin' | *C. australis* | SAMN57102153 | |
| F1_246_01 | 36,098,467 | 38.55 | 'Wilking mandarin' | *C. australis* | SAMN57102154 | |
| F1_247_01 | 36,991,508 | 39.50 | 'Wilking mandarin' | *C. australis* | SAMN57102155 | |
| F1_254_01 | 38,059,525 | 40.64 | 'Wilking mandarin' | *C. australis* | SAMN57102156 | |
| F1_313_01_95 | 46,964,002 | 50.15 | 'Wilking mandarin' | *C. australasica* | SAMN57102157 | |
| F1_334_01 | 33,669,982 | 35.95 | 'Wilking mandarin' | *C. australasica* | SAMN57102158 | |
| F1_359_01 | 35,708,076 | 38.13 | 'Wilking mandarin' | *C. inodora* | SAMN57102159 | |
| F1_411_02 | 36,032,227 | 33.99 | 'Wilking mandarin' | *C. australis* | SAMN57102160 | |
| F1_449_01 | 41,812,188 | 44.65 | 'Wilking mandarin' | *C. australasica* | SAMN57102161 | |
| F1_449_03 | 38,947,310 | 41.59 | 'Wilking mandarin' | *C. australasica* | SAMN57102162 | |
| F1_459_01 | 35,697,973 | 38.12 | 'Wilking mandarin' | *C. australasica* | SAMN57102163 | |
| F1_459_02 | 37,410,014 | 39.95 | 'Wilking mandarin' | *C. australasica* | SAMN57102164 | |
| F1_471_01 | 36,651,505 | 31.65 | 'Wilking mandarin' | *C. australasica* | SAMN57102165 | |
| F1_471_02 | 41,936,795 | 44.78 | 'Wilking mandarin' | *C. australasica* | SAMN57102166 | |
| F1_475_01 | 33,528,556 | 35.80 | 'Wilking mandarin' | *C. australasica* | SAMN57102167 | |
| F1_475_02 | 41,945,634 | 44.79 | 'Wilking mandarin' | *C. australasica* | SAMN57102168 | |
| F1_1682_01 | 34,565,450 | 36.91 | 'Fallglo mandarin' | *C. inodora* | SAMN57102169 | |
| F1_1682_02_96 | 39,806,600 | 42.51 | 'Fallglo mandarin' | *C. inodora* | SAMN57102170 | |
| F1_1689_01 | 33,212,207 | 35.46 | 'Fallglo mandarin' | *C. inodora* | SAMN57102171 | |
| F1_1689_02 | 34,477,949 | 36.82 | 'Fallglo mandarin' | *C. inodora* | SAMN57102172 | |
| F1_1729_02 | 36,007,349 | 38.45 | 'Fortune mandarin' | *C. inodora* | SAMN57102173 | |
| F1_1731_02_87 | 51,406,651 | 54.89 | 'Fortune mandarin' | *C. inodora* | SAMN57102174 | |
| F1_1735_01 | 34,681,385 | 37.03 | 'Fortune mandarin' | *C. inodora* | SAMN57102175 | |
| F1_1755_02 | 38,475,892 | 41.08 | 'Fortune mandarin' | *C. inodora* | SAMN57102176 | |
| **Advanced Hybrid Generation (n=244)** | | | | | | |
| Adv_6772_01 | 51,285,468 | 54.18 | 'Algerian Clementine' | 'Wilking' X *C. australasica* | SAMN56871903 | PRJNA1446964 |
| Adv_6772_03 | 44,421,989 | 46.48 | 'Algerian Clementine' | 'Wilking' X *C. australasica* | SAMN56871904 | |
| Adv_6772_04 | 42,409,659 | 44.01 | 'Algerian Clementine' | 'Wilking' X *C. australasica* | SAMN56871905 | |
| Adv_6772_06 | 41,311,411 | 42.77 | 'Algerian Clementine' | 'Wilking' X *C. australasica* | SAMN56871907 | |
| Adv_6772_07 | 44,736,608 | 46.94 | 'Algerian Clementine' | 'Wilking' X *C. australasica* | SAMN56871908 | |
| Adv_6772_08 | 47,828,652 | 47.59 | 'Algerian Clementine' | 'Wilking' X *C. australasica* | SAMN56871909 | |
| Adv_6772_12 | 49,171,984 | 52.23 | 'Algerian Clementine' | 'Wilking' X *C. australasica* | SAMN56871910 | |
| Adv_6774_01 | 44,938,499 | 46.73 | 'Algerian Clementine' | 'Wilking' X *C. australasica* | SAMN56871911 | |
| Adv_6774_03 | 41,430,287 | 43.37 | 'Algerian Clementine' | 'Wilking' X *C. australasica* | SAMN56871912 | |
| Adv_6774_11 | 51,716,365 | 54.63 | 'Algerian Clementine' | 'Wilking' X *C. australasica* | SAMN56871913 | |
| Adv_6778_10 | 41,527,221 | 42.98 | 'Algerian Clementine' | 'Wilking' X *C. australasica* | SAMN56871915 | |
| Adv_6778_12 | 51,861,918 | 54.37 | 'Algerian Clementine' | 'Wilking' X *C. australasica* | SAMN56871916 | |
| Adv_6778_13 | 42,588,857 | 44.66 | 'Algerian Clementine' | 'Wilking' X *C. australasica* | SAMN56871917 | |
| Adv_6778_14 | 48,966,119 | 51.29 | 'Algerian Clementine' | 'Wilking' X *C. australasica* | SAMN56871918 | |
| Adv_6778_15 | 42,566,534 | 44.64 | 'Algerian Clementine' | 'Wilking' X *C. australasica* | SAMN56871919 | |
| Adv_6778_16 | 41,050,477 | 43.16 | 'Algerian Clementine' | 'Wilking' X *C. australasica* | SAMN56871920 | |
| Adv_6778_17 | 45,634,776 | 48.11 | 'Algerian Clementine' | 'Wilking' X *C. australasica* | SAMN56871921 | |
| Adv_6778_18 | 52,174,697 | 54.23 | 'Algerian Clementine' | 'Wilking' X *C. australasica* | SAMN56871922 | |
| Adv_6779_03 | 55,316,631 | 57.82 | 'Algerian Clementine' | 'Wilking' X *C. australasica* | SAMN56871923 | |
| Adv_6779_06 | 45,887,076 | 48.14 | 'Algerian Clementine' | 'Wilking' X *C. australasica* | SAMN56871924 | |
| Adv_6779_07 | 40,057,321 | 41.39 | 'Algerian Clementine' | 'Wilking' X *C. australasica* | SAMN56871925 | |
| Adv_6779_11 | 55,777,075 | 58.40 | 'Algerian Clementine' | 'Wilking' X *C. australasica* | SAMN56871927 | |
| Adv_6779_13 | 48,153,663 | 50.75 | 'Algerian Clementine' | 'Wilking' X *C. australasica* | SAMN56871928 | |
| Adv_6779_15 | 49,221,041 | 51.79 | 'Algerian Clementine' | 'Wilking' X *C. australasica* | SAMN56871929 | |
| Adv_6791_04 | 47,898,016 | 50.24 | 'Algerian Clementine' | 'Wilking' X *C. australasica* | SAMN56871930 | |
| Adv_6791_05 | 41,469,898 | 43.53 | 'Algerian Clementine' | 'Wilking' X *C. australasica* | SAMN56871931 | |

| | | | | | | |
|---|---|---|---|---|---|---|
| Adv_6791_08 | 45,376,271 | 46.90 | 'Algerian Clementine' | 'Wilking' X *C. australasica* | SAMN56871932 | |
| Adv_6791_09 | 49,031,435 | 51.42 | 'Algerian Clementine' | 'Wilking' X *C. australasica* | SAMN56871933 | |
| Adv_6794_03 | 43,701,265 | 45.15 | 'Algerian Clementine' | 'Wilking' X *C. australasica* | SAMN56871934 | |
| Adv_6794_04 | 44,415,672 | 46.77 | 'Algerian Clementine' | 'Wilking' X *C. australasica* | SAMN56871943 | |
| Adv_6794_07 | 45,605,994 | 47.66 | 'Algerian Clementine' | 'Wilking' X *C. australasica* | SAMN56871935 | |
| Adv_6794_08 | 39,145,105 | 41.31 | 'Algerian Clementine' | 'Wilking' X *C. australasica* | SAMN56871944 | |
| Adv_6794_10 | 43,290,583 | 45.47 | 'Algerian Clementine' | 'Wilking' X *C. australasica* | SAMN56871936 | |
| Adv_6794_13 | 38,919,655 | 40.40 | 'Algerian Clementine' | 'Wilking' X *C. australasica* | SAMN56871937 | |
| Adv_6794_14 | 39,508,185 | 41.42 | 'Algerian Clementine' | 'Wilking' X *C. australasica* | SAMN56871938 | |
| Adv_6794_16 | 42,386,251 | 43.69 | 'Algerian Clementine' | 'Wilking' X *C. australasica* | SAMN56871939 | |
| Adv_6794_17 | 49,331,820 | 52.23 | 'Algerian Clementine' | 'Wilking' X *C. australasica* | SAMN56871940 | |
| Adv_6794_20 | 46,326,615 | 48.50 | 'Algerian Clementine' | 'Wilking' X *C. australasica* | SAMN56871941 | |
| Adv_6794_21 | 49,489,489 | 52.13 | 'Algerian Clementine' | 'Wilking' X *C. australasica* | SAMN56871942 | |
| Adv_7370_06 | 52,986,735 | 55.55 | 'Kiyomi Tangor-H12' | 'Wilking' X *C. australasica* | SAMN57079071 | |
| Adv_7370_11 | 49,935,058 | 52.27 | 'Kiyomi Tangor-H12' | 'Wilking' X *C. australasica* | SAMN57079072 | |
| Adv_7370_14 | 41,982,229 | 43.83 | 'Kiyomi Tangor-H12' | 'Wilking' X *C. australasica* | SAMN57079073 | |
| Adv_7371_04 | 44,691,507 | 47.15 | 'Kiyomi Tangor-H12' | 'Wilking' X *C. australasica* | SAMN57079076 | |
| Adv_7371_06 | 41,374,611 | 43.33 | 'Kiyomi Tangor-H12' | 'Wilking' X *C. australasica* | SAMN57079077 | |
| Adv_7371_07 | 43,435,855 | 45.08 | 'Kiyomi Tangor-H12' | 'Wilking' X *C. australasica* | SAMN57079074 | |
| Adv_7371_12 | 47,274,238 | 48.91 | 'Kiyomi Tangor-H12' | 'Wilking' X *C. australasica* | SAMN57079075 | |
| Adv_7372_06 | 42,511,345 | 44.45 | 'Kiyomi Tangor-H12' | 'Wilking' X *C. australasica* | SAMN57079080 | |
| Adv_7372_08 | 40,906,220 | 42.26 | 'Kiyomi Tangor-H12' | 'Wilking' X *C. australasica* | SAMN57079081 | |
| Adv_7372_09 | 46,845,491 | 48.32 | 'Kiyomi Tangor-H12' | 'Wilking' X *C. australasica* | SAMN57079082 | |
| Adv_7372_11 | 43,787,362 | 44.81 | 'Kiyomi Tangor-H12' | 'Wilking' X *C. australasica* | SAMN57079078 | |
| Adv_7372_13 | 42,317,588 | 43.20 | 'Kiyomi Tangor-H12' | 'Wilking' X *C. australasica* | SAMN57079079 | |
| Adv_7373_02 | 40,137,065 | 42.09 | 'Kiyomi Tangor-H12' | 'Wilking' X *C. australasica* | SAMN57079083 | |
| Adv_7373_04 | 47,129,185 | 49.26 | 'Kiyomi Tangor-H12' | 'Wilking' X *C. australasica* | SAMN57079084 | |
| Adv_7373_05 | 39,266,014 | 40.91 | 'Kiyomi Tangor-H12' | 'Wilking' X *C. australasica* | SAMN57079090 | |
| Adv_7373_11 | 53,484,455 | 56.13 | 'Kiyomi Tangor-H12' | 'Wilking' X *C. australasica* | SAMN57079085 | |
| Adv_7373_16 | 39,756,626 | 41.88 | 'Kiyomi Tangor-H12' | 'Wilking' X *C. australasica* | SAMN57079086 | |
| Adv_7373_17 | 50,369,254 | 53.44 | 'Kiyomi Tangor-H12' | 'Wilking' X *C. australasica* | SAMN57079087 | |
| Adv_7373_21 | 49,942,810 | 52.05 | 'Kiyomi Tangor-H12' | 'Wilking' X *C. australasica* | SAMN57079088 | |
| Adv_7373_24 | 46,349,347 | 48.22 | 'Kiyomi Tangor-H12' | 'Wilking' X *C. australasica* | SAMN57079089 | |
| Adv_7375_03 | 41,372,717 | 43.52 | 'Kiyomi Tangor-H12' | 'Wilking' X *C. australasica* | SAMN57079091 | |
| Adv_7375_04 | 51,296,643 | 54.19 | 'Kiyomi Tangor-H12' | 'Wilking' X *C. australasica* | SAMN57079097 | |
| Adv_7375_07 | 44,718,934 | 46.93 | 'Kiyomi Tangor-H12' | 'Wilking' X *C. australasica* | SAMN57079098 | |
| Adv_7375_08 | 49,396,728 | 52.15 | 'Kiyomi Tangor-H12' | 'Wilking' X *C. australasica* | SAMN57079099 | |
| Adv_7375_13 | 39,880,264 | 41.59 | 'Kiyomi Tangor-H12' | 'Wilking' X *C. australasica* | SAMN57079092 | |
| Adv_7375_14 | 48,010,651 | 50.50 | 'Kiyomi Tangor-H12' | 'Wilking' X *C. australasica* | SAMN57079093 | |
| Adv_7375_17 | 43,666,696 | 45.59 | 'Kiyomi Tangor-H12' | 'Wilking' X *C. australasica* | SAMN57079094 | |
| Adv_7375_20 | 57,912,782 | 60.69 | 'Kiyomi Tangor-H12' | 'Wilking' X *C. australasica* | SAMN57079095 | |
| Adv_7375_21 | 42,342,232 | 43.99 | 'Kiyomi Tangor-H12' | 'Wilking' X *C. australasica* | SAMN57079096 | |
| Adv_7377_03 | 51,976,655 | 55.05 | 'Kiyomi Tangor-H12' | 'Wilking' X *C. australasica* | SAMN57079104 | |
| Adv_7377_13 | 42,992,044 | 45.45 | 'Kiyomi Tangor-H12' | 'Wilking' X *C. australasica* | SAMN57079100 | |
| Adv_7377_14 | 45,629,257 | 47.77 | 'Kiyomi Tangor-H12' | 'Wilking' X *C. australasica* | SAMN57079101 | |
| Adv_7377_20 | 69,634,210 | 69.15 | 'Kiyomi Tangor-H12' | 'Wilking' X *C. australasica* | SAMN57079102 | |
| Adv_7377_21 | 46,301,423 | 48.77 | 'Kiyomi Tangor-H12' | 'Wilking' X *C. australasica* | SAMN57079103 | |
| Adv_7378_01 | 39,422,357 | 40.51 | 'Kiyomi Tangor-H12' | 'Wilking' X *C. australasica* | SAMN57079105 | |
| Adv_7378_06 | 40,195,770 | 42.32 | 'Kiyomi Tangor-H12' | 'Wilking' X *C. australasica* | SAMN57079108 | |
| Adv_7378_07 | 42,088,488 | 43.52 | 'Kiyomi Tangor-H12' | 'Wilking' X *C. australasica* | SAMN57079106 | |
| Adv_7378_09 | 39,847,599 | 41.69 | 'Kiyomi Tangor-H12' | 'Wilking' X *C. australasica* | SAMN57079107 | |
| Adv_7379_01 | 58,001,299 | 60.97 | 'Kiyomi Tangor-H12' | 'Wilking' X *C. australasica* | SAMN57079109 | |
| Adv_7379_03 | 48,234,776 | 50.54 | 'Kiyomi Tangor-H12' | 'Wilking' X *C. australasica* | SAMN57079110 | |
| Adv_7380_03 | 59,469,616 | 62.29 | 'Kiyomi Tangor-H12' | 'Wilking' X *C. australasica* | SAMN57079112 | |
| Adv_7380_05 | 44,902,023 | 46.66 | 'Kiyomi Tangor-H12' | 'Wilking' X *C. australasica* | SAMN57079111 | |
| Adv_7382_01 | 41,458,242 | 43.14 | 'Kiyomi Tangor-H12' | 'Wilking' X *C. australasica* | SAMN57079113 | |
| Adv_7382_02 | 49,122,426 | 51.62 | 'Kiyomi Tangor-H12' | 'Wilking' X *C. australasica* | SAMN57079115 | PRJNA1447590 |
| Adv_7382_05 | 41,067,393 | 41.66 | 'Kiyomi Tangor-H12' | 'Wilking' X *C. australasica* | SAMN57079114 | |
| Adv_7382_08 | 48,009,424 | 50.68 | 'Kiyomi Tangor-H12' | 'Wilking' X *C. australasica* | SAMN57079117 | |
| Adv_7387_01 | 46,348,643 | 48.19 | 'Kiyomi Tangor-H12' | 'Wilking' X *C. australasica* | SAMN57079118 | |

| | | | | | |
|---|---|---|---|---|---|
| Adv_7387_05 | 41,795,634 | 43.57 | 'Kiyomi Tangor-H12' | 'Wilking' X *C. australasica* | SAMN57079122 |
| Adv_7387_06 | 40,391,037 | 42.26 | 'Kiyomi Tangor-H12' | 'Wilking' X *C. australasica* | SAMN57079123 |
| Adv_7387_08 | 40,908,828 | 42.85 | 'Kiyomi Tangor-H12' | 'Wilking' X *C. australasica* | SAMN57079120 |
| Adv_7387_09 | 49,854,648 | 52.71 | 'Kiyomi Tangor-H12' | 'Wilking' X *C. australasica* | SAMN57079124 |
| Adv_7387_13 | 41,116,282 | 42.99 | 'Kiyomi Tangor-H12' | 'Wilking' X *C. australasica* | SAMN57079119 |
| Adv_7387_21 | 47,987,508 | 47.87 | 'Kiyomi Tangor-H12' | 'Wilking' X *C. australasica* | SAMN57079121 |
| Adv_7388_08 | 40,445,671 | 42.45 | 'Kiyomi Tangor-H12' | 'Wilking' X *C. australasica* | SAMN57079134 |
| Adv_7388_12 | 43,097,417 | 43.70 | 'Kiyomi Tangor-H12' | 'Wilking' X *C. australasica* | SAMN57079125 |
| Adv_7388_13 | 58,683,038 | 61.28 | 'Kiyomi Tangor-H12' | 'Wilking' X *C. australasica* | SAMN57079126 |
| Adv_7388_15 | 43,348,087 | 44.80 | 'Kiyomi Tangor-H12' | 'Wilking' X *C. australasica* | SAMN57079127 |
| Adv_7388_17 | 49,152,444 | 51.86 | 'Kiyomi Tangor-H12' | 'Wilking' X *C. australasica* | SAMN57079128 |
| Adv_7388_18 | 46,958,720 | 49.83 | 'Kiyomi Tangor-H12' | 'Wilking' X *C. australasica* | SAMN57079129 |
| Adv_7388_19 | 52,177,724 | 54.66 | 'Kiyomi Tangor-H12' | 'Wilking' X *C. australasica* | SAMN57079130 |
| Adv_7388_22 | 52,536,686 | 55.54 | 'Kiyomi Tangor-H12' | 'Wilking' X *C. australasica* | SAMN57079131 |
| Adv_7388_23 | 42,725,464 | 44.57 | 'Kiyomi Tangor-H12' | 'Wilking' X *C. australasica* | SAMN57079132 |
| Adv_7388_24 | 43,806,593 | 45.84 | 'Kiyomi Tangor-H12' | 'Wilking' X *C. australasica* | SAMN57079133 |
| Adv_7389_02 | 46,511,821 | 49.16 | 'Kiyomi Tangor-H12' | 'Wilking' X *C. australasica* | SAMN57079137 |
| Adv_7389_04 | 45,712,058 | 47.75 | 'Kiyomi Tangor-H12' | 'Wilking' X *C. australasica* | SAMN57079135 |
| Adv_7389_05 | 52,342,163 | 54.95 | 'Kiyomi Tangor-H12' | 'Wilking' X *C. australasica* | SAMN57079136 |
| Adv_7390_02 | 41,248,701 | 42.88 | 'Kiyomi Tangor-H12' | 'Wilking' X *C. australasica* | SAMN57079138 |
| Adv_7390_04 | 39,195,338 | 41.38 | 'Kiyomi Tangor-H12' | 'Wilking' X *C. australasica* | SAMN57079146 |
| Adv_7390_07 | 38,059,466 | 39.91 | 'Kiyomi Tangor-H12' | 'Wilking' X *C. australasica* | SAMN57079147 |
| Adv_7390_10 | 48,092,347 | 49.36 | 'Kiyomi Tangor-H12' | 'Wilking' X *C. australasica* | SAMN57079139 |
| Adv_7390_14 | 41,908,227 | 43.73 | 'Kiyomi Tangor-H12' | 'Wilking' X *C. australasica* | SAMN57079140 |
| Adv_7390_15 | 46,282,894 | 48.19 | 'Kiyomi Tangor-H12' | 'Wilking' X *C. australasica* | SAMN57079141 |
| Adv_7390_16 | 40,652,936 | 40.99 | 'Kiyomi Tangor-H12' | 'Wilking' X *C. australasica* | SAMN57079142 |
| Adv_7390_25 | 47,072,002 | 49.16 | 'Kiyomi Tangor-H12' | 'Wilking' X *C. australasica* | SAMN57079143 |
| Adv_7390_27 | 46,428,313 | 48.98 | 'Kiyomi Tangor-H12' | 'Wilking' X *C. australasica* | SAMN57079144 |
| Adv_7390_28 | 50,333,640 | 53.08 | 'Kiyomi Tangor-H12' | 'Wilking' X *C. australasica* | SAMN57079145 |
| Adv_7391_01 | 48,135,406 | 50.44 | 'Kiyomi Tangor-H12' | 'Wilking' X *C. australasica* | SAMN57079148 |
| Adv_7391_03 | 47,850,854 | 50.58 | 'Kiyomi Tangor-H12' | 'Wilking' X *C. australasica* | SAMN57079156 |
| Adv_7391_09 | 44,041,004 | 45.43 | 'Kiyomi Tangor-H12' | 'Wilking' X *C. australasica* | SAMN57079157 |
| Adv_7391_10 | 41,431,261 | 43.84 | 'Kiyomi Tangor-H12' | 'Wilking' X *C. australasica* | SAMN57079150 |
| Adv_7391_13 | 42,788,071 | 44.88 | 'Kiyomi Tangor-H12' | 'Wilking' X *C. australasica* | SAMN57079151 |
| Adv_7391_14 | 41,884,018 | 44.16 | 'Kiyomi Tangor-H12' | 'Wilking' X *C. australasica* | SAMN57079152 |
| Adv_7391_15 | 48,689,328 | 51.11 | 'Kiyomi Tangor-H12' | 'Wilking' X *C. australasica* | SAMN57079153 |
| Adv_7391_18 | 50,384,025 | 52.89 | 'Kiyomi Tangor-H12' | 'Wilking' X *C. australasica* | SAMN57079155 |
| Adv_7392_03 | 51,654,405 | 54.16 | 'Kiyomi Tangor-H12' | 'Wilking' X *C. australasica* | SAMN57079160 |
| Adv_7392_08 | 46,141,493 | 48.07 | 'Kiyomi Tangor-H12' | 'Wilking' X *C. australasica* | SAMN57079158 |
| Adv_7392_09 | 50,192,543 | 52.67 | 'Kiyomi Tangor-H12' | 'Wilking' X *C. australasica* | SAMN57079159 |
| Adv_7392_9 | 50,590,533 | 53.50 | 'Kiyomi Tangor-H12' | 'Wilking' X *C. australasica* | SAMN57079161 |
| Adv_X3228_03 | 50,006,570 | 52.51 | 'Algerian Clementine' | 'Wilking' X *C. australis* | |
| Adv_X3228_2 | 53,090,973 | 55.91 | 'Algerian Clementine' | 'Wilking' X *C. australis* | |
| Adv_X3237_02 | 44,976,394 | 47.37 | 'Algerian Clementine' | 'Wilking' X *C. australis* | |
| Adv_X3237_5 | 43,812,977 | 46.11 | 'Algerian Clementine' | 'Wilking' X *C. australis* | |
| Adv_X3237_7 | 42,196,134 | 44.35 | 'Algerian Clementine' | 'Wilking' X *C. australis* | |
| Adv_X3238_01 | 43,904,970 | 46.10 | 'Algerian Clementine' | 'Wilking' X *C. australis* | |
| Adv_X3238_02 | 43,158,592 | 45.52 | 'Algerian Clementine' | 'Wilking' X *C. australis* | |
| Adv_X3238_4 | 42,168,738 | 44.37 | 'Algerian Clementine' | 'Wilking' X *C. australis* | |
| Adv_X3238_5 | 45,626,197 | 47.86 | 'Algerian Clementine' | 'Wilking' X *C. australis* | |
| Adv_X3238_9 | 48,625,224 | 50.97 | 'Algerian Clementine' | 'Wilking' X *C. australis* | |
| Adv_X3238_20 | 40,457,967 | 42.42 | 'Algerian Clementine' | 'Wilking' X *C. australis* | |
| Adv_X3241_2 | 43,388,158 | 45.64 | 'Algerian Clementine' | 'Wilking' X *C. australis* | |
| Adv_X3241_4 | 58,814,751 | 61.75 | 'Algerian Clementine' | 'Wilking' X *C. australis* | |
| Adv_X3246_07 | 45,245,279 | 47.43 | 'Algerian Clementine' | 'Wilking' X *C. australis* | |
| Adv_X3246_5 | 40,955,747 | 42.91 | 'Algerian Clementine' | 'Wilking' X *C. australis* | |
| Adv_X3246_8 | 46,193,761 | 48.48 | 'Algerian Clementine' | 'Wilking' X *C. australis* | |
| Adv_X3246_10_1 | 48,018,881 | 50.33 | 'Algerian Clementine' | 'Wilking' X *C. australis* | |
| Adv_X3246_10_2 | 41,089,261 | 43.13 | 'Algerian Clementine' | 'Wilking' X *C. australis* | |
| Adv_X3248_01 | 42,765,435 | 44.96 | 'Algerian Clementine' | 'Wilking' X *C. australis* | |

| | | | | |
|---|---|---|---|---|
| Adv_X3252_01 | 46,771,857 | 49.17 | 'Algerian Clementine' | 'Wilking' X *C. australis* |
| Adv_X3252_2 | 43,824,128 | 45.80 | 'Algerian Clementine' | 'Wilking' X *C. australis* |
| Adv_X3252_3 | 39,416,147 | 41.57 | 'Algerian Clementine' | 'Wilking' X *C. australis* |
| Adv_X3252_09 | 64,875,302 | 67.80 | 'Algerian Clementine' | 'Wilking' X *C. australis* |
| Adv_X3252_11 | 48,246,298 | 50.81 | 'Algerian Clementine' | 'Wilking' X *C. australis* |
| Adv_X3252_12 | 48,533,634 | 51.05 | 'Algerian Clementine' | 'Wilking' X *C. australis* |
| Adv_X3252_15 | 39,260,344 | 41.03 | 'Algerian Clementine' | 'Wilking' X *C. australis* |
| Adv_X3252_15 | 46,305,318 | 48.03 | 'Algerian Clementine' | 'Wilking' X *C. australis* |
| Adv_X3253_06 | 54,647,933 | 56.34 | 'Algerian Clementine' | 'Wilking' X *C. australis* |
| Adv_X3253_07 | 43,583,137 | 45.77 | 'Algerian Clementine' | 'Wilking' X *C. australis* |
| Adv_X3255_01 | 44,691,351 | 46.85 | 'Algerian Clementine' | 'Wilking' X *C. australis* |
| Adv_X3255_3 | 46,271,653 | 48.64 | 'Algerian Clementine' | 'Wilking' X *C. australis* |
| Adv_X3255_5 | 41,520,928 | 43.51 | 'Algerian Clementine' | 'Wilking' X *C. australis* |
| Adv_X3255_08 | 44,427,149 | 46.61 | 'Algerian Clementine' | 'Wilking' X *C. australis* |
| Adv_X3255_13 | 42,834,649 | 44.87 | 'Algerian Clementine' | 'Wilking' X *C. australis* |
| Adv_X3255_77 | 47,180,308 | 49.67 | 'Algerian Clementine' | 'Wilking' X *C. australis* |
| Adv_X3260_01 | 45,311,296 | 47.51 | 'Algerian Clementine' | 'Wilking' X *C. australis* |
| Adv_X3260_5 | 42,916,761 | 45.00 | 'Algerian Clementine' | 'Wilking' X *C. australis* |
| Adv_X3260_7 | 44,542,507 | 46.93 | 'Algerian Clementine' | 'Wilking' X *C. australis* |
| Adv_X3260_08 | 48,984,179 | 51.38 | 'Algerian Clementine' | 'Wilking' X *C. australis* |
| Adv_X3260_10 | 42,885,112 | 44.98 | 'Algerian Clementine' | 'Wilking' X *C. australis* |
| Adv_X3260_11 | 43,419,685 | 45.84 | 'Algerian Clementine' | 'Wilking' X *C. australis* |
| Adv_X3260_13 | 47,223,934 | 49.57 | 'Algerian Clementine' | 'Wilking' X *C. australis* |
| Adv_X3261_10 | 40,989,739 | 43.06 | 'Algerian Clementine' | 'Wilking' X *C. australis* |
| Adv_X3261_11 | 42,965,697 | 45.13 | 'Algerian Clementine' | 'Wilking' X *C. australis* |
| Adv_X3261_4 | 51,177,586 | 53.76 | 'Algerian Clementine' | 'Wilking' X *C. australis* |
| Adv_X3263_01 | 53,644,433 | 56.34 | 'Algerian Clementine' | 'Wilking' X *C. australis* |
| Adv_X3263_2 | 42,508,036 | 44.81 | 'Algerian Clementine' | 'Wilking' X *C. australis* |
| Adv_X3263_4 | 43,856,364 | 45.95 | 'Algerian Clementine' | 'Wilking' X *C. australis* |
| Adv_X3263_06 | 41,531,471 | 43.48 | 'Algerian Clementine' | 'Wilking' X *C. australis* |
| Adv_X3268_01 | 45,557,725 | 47.77 | 'Algerian Clementine' | 'Wilking' X *C. australis* |
| Adv_X3268_03 | 45,633,632 | 48.06 | 'Algerian Clementine' | 'Wilking' X *C. australis* |
| Adv_X3268_8 | 43,837,717 | 46.05 | 'Algerian Clementine' | 'Wilking' X *C. australis* |
| Adv_X3268_01 | 44,620,314 | 46.74 | 'Algerian Clementine' | 'Wilking' X *C. australis* |
| Adv_X3268_03 | 51,268,343 | 53.79 | 'Algerian Clementine' | 'Wilking' X *C. australis* |
| Adv_X3275_01 | 44,849,856 | 47.24 | 'Algerian Clementine' | 'Wilking' X *C. australis* |
| Adv_X3275_3 | 39,463,457 | 41.44 | 'Algerian Clementine' | 'Wilking' X *C. australis* |
| Adv_X3275_5 | 42,730,743 | 44.94 | 'Algerian Clementine' | 'Wilking' X *C. australis* |
| Adv_X3275_01 | 62,268,286 | 65.69 | 'Algerian Clementine' | 'Wilking' X *C. australis* |
| Adv_X3282_01 | 51,146,610 | 53.84 | 'Algerian Clementine' | 'Wilking' X *C. australis* |
| Adv_X3282_03 | 45,166,351 | 47.30 | 'Algerian Clementine' | 'Wilking' X *C. australis* |
| Adv_X3282_6 | 47,303,644 | 49.90 | 'Algerian Clementine' | 'Wilking' X *C. australis* |
| Adv_X3282_09 | 47,496,471 | 49.75 | 'Algerian Clementine' | 'Wilking' X *C. australis* |
| Adv_X3282_10 | 43,355,913 | 45.29 | 'Algerian Clementine' | 'Wilking' X *C. australis* |
| Adv_X3288_06 | 42,255,487 | 44.33 | 'Algerian Clementine' | 'Wilking' X *C. australis* |
| Adv_X3288_10 | 47,782,648 | 50.14 | 'Algerian Clementine' | 'Wilking' X *C. australis* |
| Adv_X3288_12 | 53,180,133 | 55.51 | 'Algerian Clementine' | 'Wilking' X *C. australis* |
| Adv_X3288_7 | 43,065,030 | 45.41 | 'Algerian Clementine' | 'Wilking' X *C. australis* |
| Adv_X3303_01 | 45,682,453 | 47.99 | 'Algerian Clementine' | 'Wilking' X *C. australis* |
| Adv_X3303_05 | 38,966,321 | 40.82 | 'Algerian Clementine' | 'Wilking' X *C. australis* |
| Adv_X3303_10 | 41,552,932 | 43.28 | 'Algerian Clementine' | 'Wilking' X *C. australis* |
| Adv_X3303_15 | 47,915,012 | 50.34 | 'Algerian Clementine' | 'Wilking' X *C. australis* |
| Adv_X3303_3 | 44,361,847 | 46.69 | 'Algerian Clementine' | 'Wilking' X *C. australis* |
| Adv_X3303_10 | 57,075,733 | 60.23 | 'Algerian Clementine' | 'Wilking' X *C. australis* |
| Adv_X3322_01 | 43,749,887 | 45.91 | 'Algerian Clementine' | 'Wilking' X *C. australis* |
| Adv_X3322_02 | 41,609,664 | 51.11 | 'Algerian Clementine' | 'Wilking' X *C. australis* |
| Adv_X3322_11 | 44,554,194 | 46.58 | 'Algerian Clementine' | 'Wilking' X *C. australis* |
| Adv_X3322_12 | 50,165,122 | 52.50 | 'Algerian Clementine' | 'Wilking' X *C. australis* |
| Adv_X3322_17 | 42,794,607 | 45.01 | 'Algerian Clementine' | 'Wilking' X *C. australis* |
| Adv_X3322_20 | 43,090,176 | 45.17 | 'Algerian Clementine' | 'Wilking' X *C. australis* |

| | | | | |
|---|---|---|---|---|
| Adv_X3322_2 | 48,642,412 | 51.11 | 'Algerian Clementine' | 'Wilking' X *C. australis* |
| Adv_X3322_3 | 45,504,367 | 47.91 | 'Algerian Clementine' | 'Wilking' X *C. australis* |
| Adv_X3322_5 | 53,508,264 | 56.39 | 'Algerian Clementine' | 'Wilking' X *C. australis* |
| Adv_X3322_6 | 46,293,973 | 48.80 | 'Algerian Clementine' | 'Wilking' X *C. australis* |
| Adv_X3323_06 | 53,239,249 | 55.26 | 'Algerian Clementine' | 'Wilking' X *C. australis* |
| Adv_X3323_11 | 54,027,428 | 56.59 | 'Algerian Clementine' | 'Wilking' X *C. australis* |
| Adv_X3323_13 | 42,947,509 | 45.21 | 'Algerian Clementine' | 'Wilking' X *C. australis* |
| Adv_X3323_15 | 44,826,100 | 46.99 | 'Algerian Clementine' | 'Wilking' X *C. australis* |
| Adv_X3323_2 | 41,746,796 | 43.87 | 'Algerian Clementine' | 'Wilking' X *C. australis* |
| Adv_X3323_4 | 43,086,952 | 45.14 | 'Algerian Clementine' | 'Wilking' X *C. australis* |
| Adv_X3323_9 | 42,006,184 | 44.16 | 'Algerian Clementine' | 'Wilking' X *C. australis* |
| Adv_X3326_03 | 42,761,338 | 44.92 | 'Algerian Clementine' | 'Wilking' X *C. australis* |
| Adv_X3326_2 | 44,367,424 | 46.77 | 'Algerian Clementine' | 'Wilking' X *C. australis* |
| Adv_X3326_6 | 43,071,847 | 45.34 | 'Algerian Clementine' | 'Wilking' X *C. australis* |
| Adv_X3326_03 | 48,143,008 | 50.38 | 'Algerian Clementine' | 'Wilking' X *C. australis* |
| Adv_X3328_8 | 43,135,583 | 45.50 | 'Algerian Clementine' | 'Wilking' X *C. australis* |
| Adv_X3329_05 | 42,810,075 | 44.90 | 'Algerian Clementine' | 'Wilking' X *C. australis* |
| Adv_X3329_3 | 43,509,958 | 45.73 | 'Algerian Clementine' | 'Wilking' X *C. australis* |
| Adv_X3329_4 | 40,909,273 | 42.91 | 'Algerian Clementine' | 'Wilking' X *C. australis* |
| Adv_X3329_6 | 61,902,842 | 65.20 | 'Algerian Clementine' | 'Wilking' X *C. australis* |
| Adv_X3332_01 | 48,016,699 | 50.38 | 'Algerian Clementine' | 'Wilking' X *C. australis* |
| Adv_X3332_06 | 42,044,960 | 44.40 | 'Algerian Clementine' | 'Wilking' X *C. australis* |
| Adv_X3332_10 | 47,062,060 | 49.55 | 'Algerian Clementine' | 'Wilking' X *C. australis* |
| Adv_X3332_2 | 46,485,617 | 48.99 | 'Algerian Clementine' | 'Wilking' X *C. australis* |
| Adv_X3332_4 | 43,681,957 | 45.84 | 'Algerian Clementine' | 'Wilking' X *C. australis* |
| Adv_X3332_5 | 39,536,101 | 41.69 | 'Algerian Clementine' | 'Wilking' X *C. australis* |
| Adv_X3332_7 | 51,880,512 | 54.67 | 'Algerian Clementine' | 'Wilking' X *C. australis* |
| Adv_X3332_9 | 50,241,521 | 52.82 | 'Algerian Clementine' | 'Wilking' X *C. australis* |
| Adv_X3333_2 | 42,274,082 | 44.04 | 'Algerian Clementine' | 'Wilking' X *C. australis* |
| Adv_X3333_3 | 49,014,411 | 51.31 | 'Algerian Clementine' | 'Wilking' X *C. australis* |
| Adv_X3333_4 | 43,326,656 | 45.38 | 'Algerian Clementine' | 'Wilking' X *C. australis* |
| Adv_X3333_5 | 40,881,912 | 43.00 | 'Algerian Clementine' | 'Wilking' X *C. australis* |
| Adv_X3333_9 | 42,931,630 | 45.04 | 'Algerian Clementine' | 'Wilking' X *C. australis* |
| Adv_X3340_14 | 43,452,666 | 45.74 | 'Algerian Clementine' | 'Wilking' X *C. australis* |
| Adv_X3340_15 | 51,319,809 | 53.82 | 'Algerian Clementine' | 'Wilking' X *C. australis* |
| Adv_X3340_4 | 45,654,695 | 47.75 | 'Algerian Clementine' | 'Wilking' X *C. australis* |
| Adv_X3340_5 | 42,500,925 | 44.84 | 'Algerian Clementine' | 'Wilking' X *C. australis* |
| Adv_X3340_6 | 45,573,840 | 47.90 | 'Algerian Clementine' | 'Wilking' X *C. australis* |
| Other Samples Included with the 244 Advanced Hybrids (n=7) | | | | |
| F1_246_01 | 36,098,467 | 36.30 | 'Wilking' | *C. australis* |
| F1_475_01 | 33,528,556 | 41.95 | 'Wilking' | *C. australasica* |
| *'Kiyomi Tangor-H12'* | 40,100,970 | 41.04 | | |
| *'Algerian Clementine'* | 33,507,185 | 34.51 | | |
| *C. australasica* | 36,943,740 | 35.41 | | |
| *C. australis* | 37,879,389 | 37.13 | | |
| 'Wilking' | 36,280,782 | 36.72 | | |

**Supplemental Table 3**. Comparison of genotype frequency and average depth in regions of the masked vcf (mask) vs. regions only within masking ranges (mask inverse). Percentages calculated from the total loci within each vcf. Asterisks denote samples present as founder lines in the graph pangenome.

| Sample | Reference | Region | Depth | Hom_ref | Hom_alt | Het | Missing |
|---|---|---|---|---|---|---|---|
| **C. reticulata*** | graph | mask | 22.8 | 79.61% | 4.94% | 13.01% | 2.45% |
| | | mask inverse | 26.1 | 68.06% | 8.67% | 14.35% | 8.92% |
| | linear | mask | 15.3 | 79.06% | 4.20% | 14.20% | 2.54% |
| | | mask inverse | 21.8 | 77.80% | 3.12% | 12.72% | 6.36% |
| **C. australis*** | graph | mask | 19.9 | 44.30% | 32.70% | 14.49% | 8.51% |
| | | mask inverse | 12.5 | 15.95% | 5.97% | 3.66% | 74.42% |
| | linear | mask | 13.5 | 45.03% | 25.33% | 21.30% | 8.34% |
| | | mask inverse | 22.8 | 45.65% | 8.42% | 18.95% | 26.98% |
| **C. australasica*** | graph | mask | 15.8 | 46.36% | 28.58% | 15.31% | 9.75% |
| | | mask inverse | 13 | 17.81% | 6.55% | 4.31% | 71.32% |
| | linear | mask | 10.8 | 47.16% | 21.34% | 21.10% | 10.41% |
| | | mask inverse | 22.8 | 46.14% | 7.87% | 17.44% | 28.55% |
| **Kiyomi Tangor** | graph | mask | 17.1 | 66.36% | 7.91% | 19.62% | 6.11% |
| | | mask inverse | 29.8 | 53.16% | 12.60% | 22.68% | 11.56% |
| | linear | mask | 23.4 | 65.89% | 6.15% | 21.13% | 6.83% |
| | | mask inverse | 33.4 | 66.52% | 4.94% | 20.27% | 8.27% |
| **Algerian Clementine** | graph | mask | 16.8 | 79.92% | 2.68% | 15.91% | 1.49% |
| | | mask inverse | 25 | 73.53% | 4.77% | 17.72% | 3.98% |
| | linear | mask | 10.9 | 79.87% | 2.12% | 16.42% | 1.60% |
| | | mask inverse | 20 | 81.04% | 1.73% | 13.94% | 3.29% |
| **F1: 475_01** C. reticulata x C. australasica | graph | mask | 20.9 | 54.30% | 7.03% | 35.39% | 3.28% |
| | | mask inverse | 15.3 | 65.19% | 11.41% | 7.20% | 16.20% |
| | linear | mask | 11 | 54.88% | 5.49% | 36.25% | 3.37% |
| | | mask inverse | 24.7 | 68.25% | 4.81% | 19.10% | 7.84% |
| **F1: 246_01** C. reticulata x C. australis | graph | mask | 19.7 | 53.16% | 6.92% | 35.31% | 4.60% |
| | | mask inverse | 16 | 65.59% | 10.84% | 6.95% | 16.62% |
| | linear | mask | 13.2 | 53.51% | 5.34% | 36.58% | 4.57% |
| | | mask inverse | 17.3 | 67.63% | 4.46% | 19.65% | 8.25% |
